# INTS12 Bridges Integrator and NELF to Prevent the Release of Non-processive RNA Polymerase II Complexes

**DOI:** 10.64898/2026.01.16.699819

**Authors:** Laura D. Corso, Isela Sarahi Rivera, Aleen Al Halawani, Chloe A. N. Gerak, Dáire Gannon, Oliver Ozaydin, Mengbo Li, Kedhar Thyagarajan, Lukasz Niezabitowski, Ngee Kiat Chua, Winnie Tan, Susanne I. Wudy, Aoife McLysaght, Gordon K. Smyth, Shabih Shakeel, Rebecca Feltham, Stephin J. Vervoort

## Abstract

Promoter-proximal RNA Polymerase II (RNAPII) pausing and the processivity are controlled by distinct modules of the Integrator complex, which together fine-tune transcription and protect against the accumulation of defective RNAPII complexes. Compromised activity of individual Integrator modules has been linked to human disease including cancer and developmental disorders, caused by defective transcription of protein-coding or small-nuclear RNAs. Despite extensive characterisation of the Integrator complex both genetically and structurally, the role of smallest member of the complex, INTS12, has remained enigmatic. Here, we uncover that INTS12 loss acts to stabilise the association between NELF and Integrator via its PHD domain and N-terminus, respectively, thus safeguarding against the release of defective RNAPII complexes. Acute degradation of INTS12 results in the selective dissociation of Integrator from the NELF-RNAPII complex which subsequently convert to their canonical paused form from which they can be released by CDK9. In the absence of INTS12 excess release of defective RNAPII via P-TEFb/SEC, loss of the ARMC5 salvage pathway and deletion of the catalytic and core Integrator subunits is toxic to cells. These findings demonstrate that there is interconversion between canonical paused RNAPII and paused-Integrator, and highlight the critical interplay between these processes and P-TEFb mediated pause-release to ensure that only transcription competent complexes are released into elongation.

- INTS12 degradation confers CDK9 inhibitor resistance and triggers cellular stress through a phosphatase module-independent mechanism.
- INTS12 stabilizes the Integrator-NELF complex through its N-terminus and PHD domain.
- Acute INTS12 degradation promotes aberrant release of promoter-proximal RNA polymerase II complexes.
- RNA polymerase II complexes released upon INTS12 loss exhibit defective elongation and reduced processivity.
- INTS12 loss removes Integrator from RNAPII resulting in aberrant paused-state from which it can be released by CDK9.
- Excess CDK9 activity and ARMC5 loss are synthetically lethal with INTS12 deficiency.

## Introduction

The RNA Polymerase II (RNAPII) transcription cycle is dynamically governed by transcriptional CDKs, which license progression of messenger RNA synthesis through discrete checkpoints (1). The controlled release of RNAPII at each stage ensures tight control over spatial-temporal gene expression patterns in response to intrinsic and extrinsic cellular cues (2). As such, each checkpoint serves as an integration hub for these signals to coordinate the sequential exchange of RNAPII-interacting complexes at discrete stages, ensuring proper control over processivity, termination, splicing, and transcript quality.

In metazoans, RNAPII comes to a halt post-initiation at Transcription Start Site (TSS) proximal regions (40-100 bp), where it adopts a “paused” conformation in complex with the Negative Elongation Factor (NELF; comprising NELFA, NELFE and NELFCD) and the DRB Sensitivity Inducing Factor (DSIF) containing SPT5 and SPT4. At this pause-release checkpoint RNAPII remains engaged in transcription but is unable to enter productive elongation until NELF, DSIF, and the RNAPII C-Terminal Domain (CTD) are phosphorylated by CDK9 in complex with Cyclin T, also known as the Positive Transcription Elongation Factor b (P-TEFb) (3). In concert, these phosphorylation events enact the release of NELF, a conformational change in DSIF, and the subsequent association with elongation factors such as the PAF complex and SPT6 (4). Expanding on this paradigm, recent discoveries have highlighted that TSS-proximal termination is more widespread than a single defined pausing-site, instead occurring within a broader pausing-zone in which RNAPII has an increased likelihood of pausing or terminating (5). Indeed, even upon acute NELF depletion, RNAPII fails to enter the elongation phase of transcription and instead halts at a discrete secondary pause-site, insensitive to P-TEFb activity (6).

The Integrator complex was originally discovered the key regulator of small nuclear RNA (snRNA) 3’-end processing (7, 8) mediated via its catalytic endonuclease activity of its subunit INTS11 (CPSF3L). Since then, it has been demonstrated to play critical roles in the early stages of the transcription cycle as a regulator of pausing (9, 10) termination and release of RNAPII. Integrator consists of 15 subunits organised into distinct functional modules: the enhancer (arm/tail/auxiliary), scaffold, phosphatase, and cleavage (endonuclease) modules (11). Recent, biochemical and structural work has demonstrated that Integrator further associates with the SOSS complex (INIP, NABP1/2) which facilitates single-strand DNA binding at R-loops as well as the formation of a “paused” Integrator state in complex with NELF and DSIF (4, 12–15).

Mechanistically, these functions depend on the endonuclease activity of Integrator, which is required for the release of stalled, non-productive RNAPII at TSS proximal regions thus preventing the accumulation of defective transcriptional complexes. Accordingly, rapid depletion of INTS11 results in an accumulation of defective RNAPII concomitant with widespread disruption of normal transcription. Recent work has revealed that these stalled TSS-proximal RNAPII complexes can be cleared by ARMC5-mediated ubiquitination and degradation, the disruption of which is synthetically lethal with Integrator dysfunction (16, 17). The second catalytic activity of Integrator lies in its phosphatase module, composed of INTS6, INTS8 together with the scaffold (A) and catalytic (C) subunits of PP2A, which enforces a paused state by dephosphorylating Serine 2 (Ser2) and Serine 5 (Ser5) of RNAPII’s CTD, thus counteracting early transcription cycle kinases such as CDK7 and CDK9. Indeed, depletion of INTS6 and INTS8 culminated in the accumulation of phosphorylated Ser2 and Ser5 on RNAPII’s CTD, reduced sensitivity to CDK9 inhibition (CDK9i), and notably promoted the release of RNAPII into elongation (18–22).

In line with its key role in transcription, genetic mutations within integrator components such as INTS1, INTS8 and INTS11 lead to developmental disorders (23–26). Disease phenotypes for INTS1 and INTS8 have recently been linked to the activation of the cellular stress response and subsequent induction of inflammatory signals (27). Aberrant integrator function can also contribute to cancer, and INTS6 loss has been observed in a variety of cancers and has been demonstrated to impact tumorigenesis (28, 29). Additional human diseases may be connected to Integrator including COPD which was been linked to INTS12 in GWAS studies (30).

Although structural and functional studies have resolved the function of the key Integrator modules, the role of INTS12 has remained largely unknown. INTS12 contains a PHD domain and a conserved N-terminal α-helical domain that mediates interaction with INTS1 and is sufficient to maintain snRNA processing in *Drosophila melanogaster* (31). Here, through integrated genomics, proteomics, and CRISPR-screening approaches, combined with endogenously tagged TurboID and dTAG^V^-1 alleles, we comprehensively characterised the role of INTS12 in the paused Integrator state. Rapid depletion of INTS12 elicited a strong induction of an intracellular stress response and resulted in an increased release of RNAPII at TSS-proximal regions with reduced overall processivity, indicative of the release of defective RNAPII complexes into elongation. Mechanistically, INTS12 depletion did not disrupt the Integrator complex but resulted in the dissociation between Integrator and NELF, a phenotype dependent on both its N-terminal and PHD domains, the latter of which mediates the interaction with NELFA. The NELF complex remains bound to chromatin in the absence of INTS12 indicating the conversion to a canonical paused state or the release of defective complexes. Genetically and pharmacologically, the impact of INTS12 loss can be partially negated by reduced release of RNAPII from the paused-state via inhibition of CDK9 or the deletion of ELL/MED8, with synthetic lethality arising from the deletion of the ARMC5 salvage pathway, core Integrator components, or negative regulators of pause-release.

## Results

### INTS12 degradation triggers a cellular stress response and opposes CDK9 activity

The Integrator complex forms a stable paused state with NELF and DSIF that is mutually exclusive with the canonical paused state (13), but the recruitment of Integrator to RNAPII and the interconversion between these distinct paused RNAPII states remains unknown. Although structurally unresolved and poorly characterised, our previous work suggested that INTS12 could play a critical role in the promoter-proximal regulation of RNAPII as its depletion mediated resistance to CDK9 inhibition (18), **Supplementary Figure 1A**).

To investigate the role of INTS12 in the Integrator complex and the transcription cycle, we knocked-in the FKBP12^F36V^ degron tag into the C-terminus of the INTS12 locus in eHAP1 cells (**Figure 1A**). Treatment with the von Hippel-Lindau (VHL) recruiting heterobifunctional dTAG^V^-1 ligand (250nM) resulted in near complete loss of INTS12 in 2 hours. Acute INTS12 degradation did not impact the expression of other Integrator components, including INTS1, INTS3, INTS5, INTS6, and INTS10, nor did affect the expression of NELFA (**Figure 1B**). To measure the impact of INTS12 degradation on cell growth, we performed a competition assay between INTS12 degron-tagged cells and wild-type (WT) cells in the presence of dTAG^V^-1 or DMSO (**Supplementary Figure 1B**). dTAG^V^-1 treatment resulted in the steady outgrowth of WT cells over their degron-tagged counterparts, indicating that INTS12 negatively affected cell growth (**Figure 1C**). Further analysis of cell division using Cell-Trace Violet (CTV) staining revealed modest decreases in the division rate upon INTS12 degradation (**Supplementary Figure 1C-D**). To assess the transcriptome-wide impact of INTS12 depletion we performed RNA-sequencing in the presence or absence of dTAG^V^-1 (**Supplementary Figure 1E**). This revealed that INTS12 degradation resulted in 185 differentially expressed genes (LFC |1|. Adj. P value < 0.01), the majority of which were induced (**Figure 1D**). Among these, stress responsive genes such *ATF3*, *DDIT3*, and *GADD45B* were significantly upregulated. Pathway and gene-set enrichment analysis revealed that the induction of stress responsive genes was concomitant with the activation of genes involved in pro-inflammatory NFkB/TNF signalling, UV response, hypoxia, and P53-signalling (**Figure 1E, Supplementary Figure 1F-G**). Activation of cellular stress was further underscored by immunoblotting ATF3 and demonstrating its time-dependent induction post-INTS12 degradation, whereas transcriptional changes in *HEXIM1* did not translate into increased overall protein levels. In addition, the loss of INTS12 did not impact the levels of RNAPII even after 8 hours of dTAG^V^-1 treatment (**Figure 1F**).

**Figure 1:**
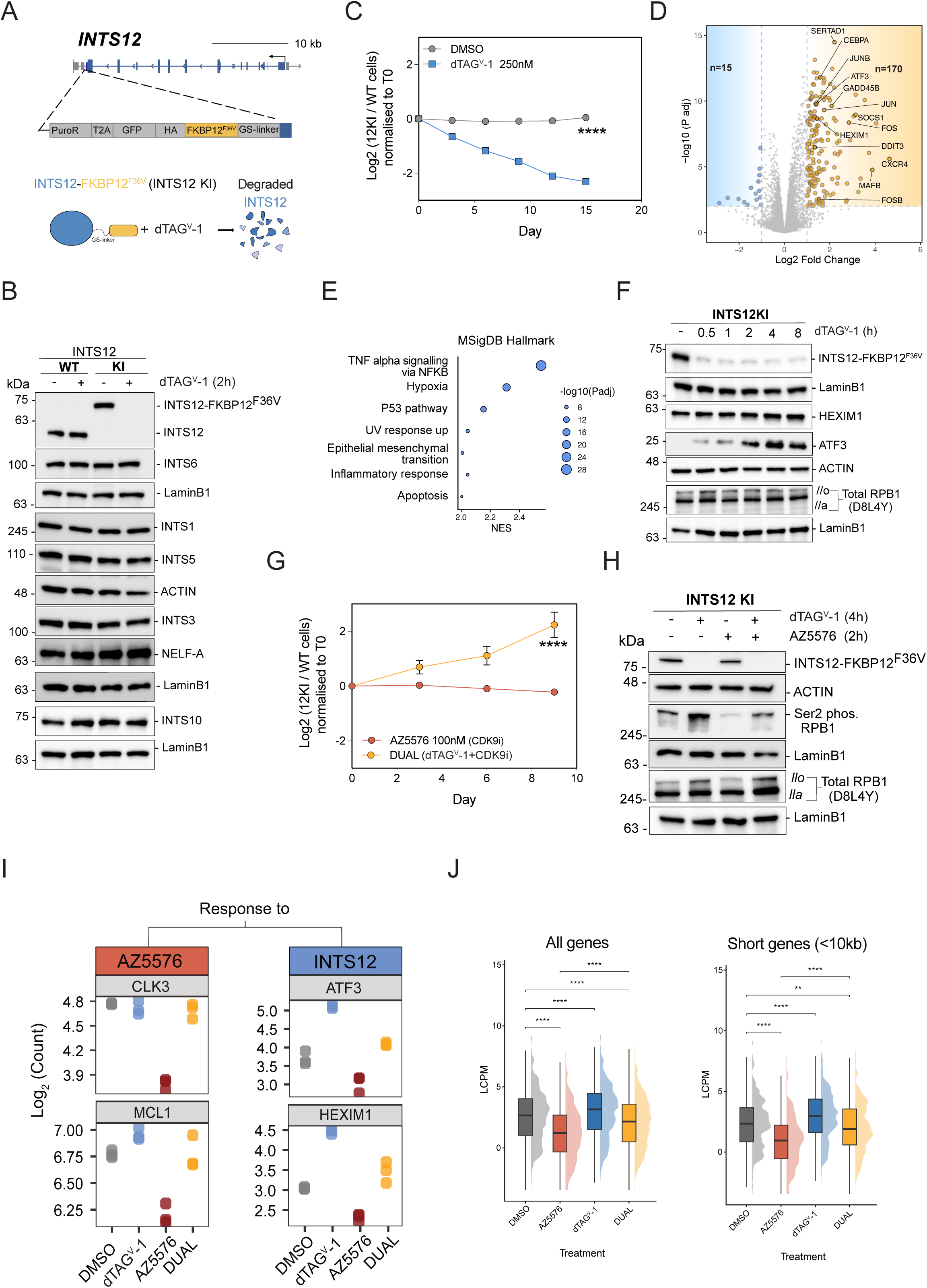
INTS12 degradation triggers a cellular stress response and opposes CDK9 activity. A) Schematic of INTS12 degron knock-in; The FKBP12^F36V^ degron tag was inserted at the C-terminus of the endogenous INTS12 locus in eHAP1 cells. The addition of dTAG^V^-1 induces the degradation of INTS12 degron-tagged. B) Immunoblots of WT and INTS12-FKBP12^F36V^ eHAP1 cells probing for Integrator components and NELFA after treatment with dTAG^V^-1 for 2h to induce degradation of INTS12. LaminB1 and actin are loading controls. C) Competitive proliferation assay; INTS12-FKBP12^F36V^ cells put in competition 1:1 with WT eHAP1s treated with DMSO or dTAG^V^-1. This is representative of n = 2 replicates. D) Volcano plot showing differentially expressed genes upon 2h dTAG^V^-1 treatment. Significantly enriched (*n = 170*) and depleted (*n = 15*) genes are based on a log2FC cut-off >|1| and adjusted p value (p.adj) < 0.01. E) Gene ontology analysis of enriched genes defined in (D). The top 7 MSigDB hallmark gene sets are shown. NES, Normalised Enrichment Score. F) Immunoblots from a time course of dTAG^V^-1 treatment probing for INTS12, HEXIM1, ATF3 and RNAPII-NTD. LaminB1 and actin are loading controls. G) INTS12-KI cells put in competition with WT eHAP1s treated with CDK9i or dual dTAG-V1 and CDK9i (AZ5576). Data is representative of *n = 2* replicates. (C) and (G) were analysed using 2-way ANOVA, ****p < 0.0001, *p < 0.05. H) Immunoblots from untreated, dTAG^V^-1, AZ5576, and dual treated cells probing for INTS12 RNAPII Ser2P and NTD showcasing the hyper-phosphorylated (RNAPIIo) and hypo-phosphorylated (RNAPIIa) form of RNAPII (*n = 2*). I) RNA Expression levels of CDK9i responsive genes (*SERTAD1* and *MCL1*) and INTS12 responsive genes (*ATF3* and *HEXIM1*) in DMSO, dTAG^V^-1, AZ5576, and dual treated cells (*n = 3*). J) Distribution of aggregated counts for all genes and a subset of short genes (<10kb) across the four different treatments. LCPM, Log Counts Per Million; **p < 0.01, ^∗∗∗∗^p < 0.0001.

As we had previously discovered that INTS12 loss confers resistance to CDK9 inhibition in genome-wide CRISPR screens (**Supplementary Figure 1A**), we next studied the role of INTS12 in this context. To this end, we performed competition assays between INTS12 degron-tagged and WT cells in the presence or absence of the selective CDK9 inhibitor, AZ5576 (100nM) (**Supplementary Figure 1H**). This revealed that INTS12 loss conferred robust resistance to CDK9 inhibition with degron-tagged lines rapidly outcompeting WT counterparts (**Figure 1G**). To assess the impact of INTS12-depletion on CDK9 dependent phosphorylation of RNAPII we measured the phosphorylation levels of the Ser2 residue in the 52 heptad repeats within the CTD of RNAPII (1, 32).This demonstrated that acute INTS12 depletion increases Ser2^P^ and RNAPIIo levels without affecting its hypo-phosphorylated counterpart (RNAPIIa) (**Figure 1H**). Importantly, Ser2^P^ and RNAPIIo levels in dTAG^V^-1 treated cells persist even after inhibiting CDK9, whereas DMSO treated cells exhibit rapid loss of phosphorylation. Together these observations indicate the functional and molecular antagonism between INTS12 and CDK9, akin to that observed with INTS6/PP2A (18). To study whether this antagonism was also reflected on the transcriptional level, we performed RNA-Seq analysis in the presence and absence of AZ5576 (**Supplementary Figure 1I**). These data demonstrated the reciprocal nature between CDK9 responses and INTS12, as evidenced by the opposed regulation of key target genes, *HEXIM1*, *ATF3*, *CLK3*, *MCL*1, and others (**Figure 1I, Supplementary Figure 1J**). Transcriptome-wide analysis of this effect demonstrated that responsive genes to CDK9 inhibition (Adj. Pvalue < 0.01 and LogFC < -1) were significantly induced upon INTS12 degradation, with a diminished reduction in their expression upon dual targeting as compared to CDK9 inhibition alone (**Figure 1J**). The transcriptional interplay between INTS12 and CDK9 was most evident for short transcripts (<10kb) in the genome compared to medium (10-50 kb) or long (>50 kb) ones, suggesting that the rescue is skewed, possibly owing to downstream transcriptional defects for CDK9-released RNAPII complexes (**Supplementary Figure 1L**).

### INTS12 controls the stable assembly of the paused Integrator-NELF complex

To investigate how INTS12, as a non-catalytic component of Integrator, can control the transcriptional response and how it can mechanistically oppose CDK9 activity, we comprehensively profiled the INTS12 interactome using three orthogonal proteomics approaches; including antibody mediated immunoprecipitation (IP) of the unmodified endogenous protein, SLF-biotin mediated affinity purification (AP) of the FKBP12^F36V^ endogenously-tagged INTS12, and proximity biotin-labelling via endogenously-tagged INTS12-TurboID (**Figure 2A, Supplementary Figure 2A, B and C**).

**Figure 2:**
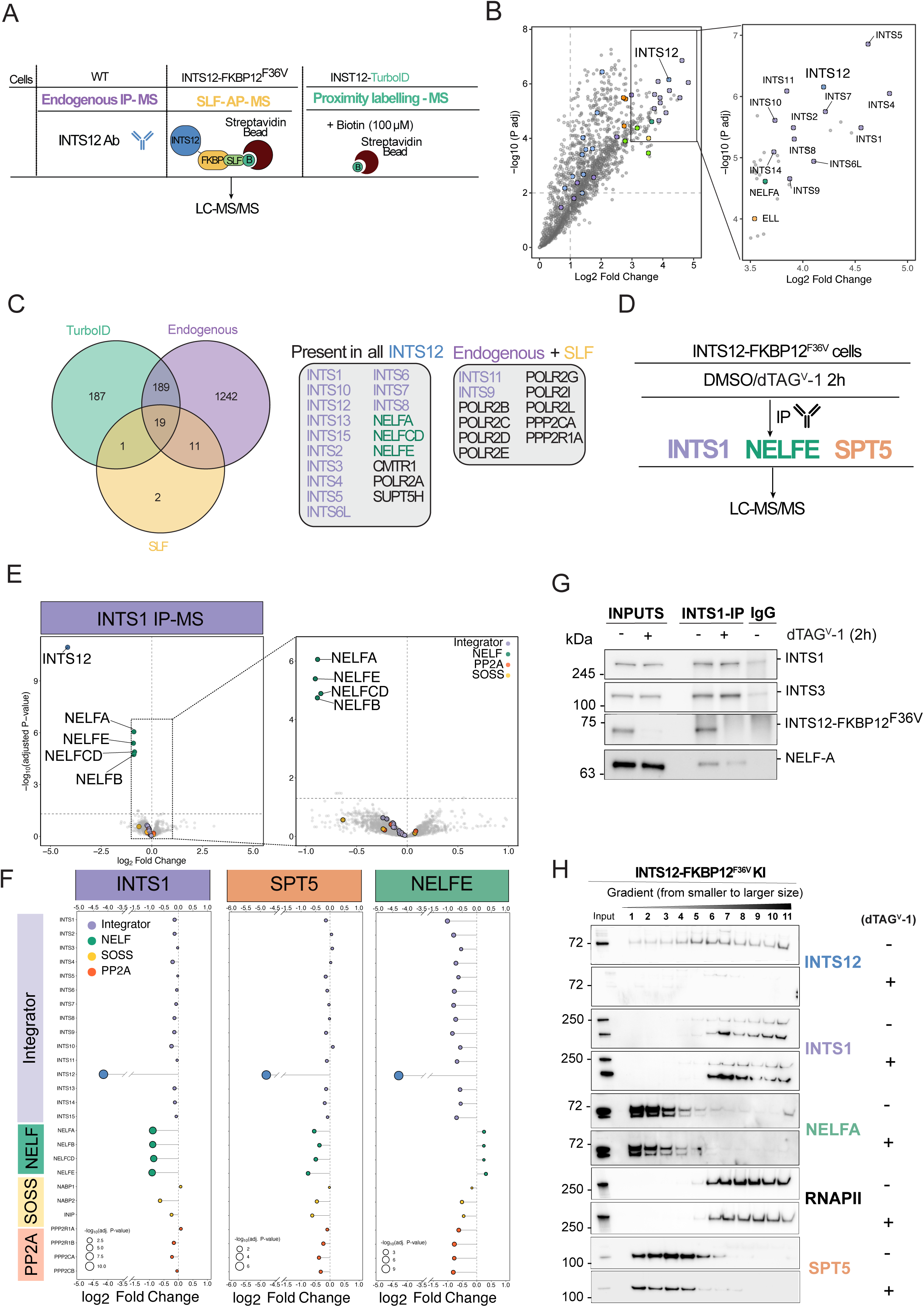
INTS12 controls the stable assembly of the Paused Integrator-NELF complex. A) Schematic of the three different MS techniques used to investigate the INTS12 interactome; Endogenous IP-mass spectrometry (IP-MS), Selective Ligand for FKBP-Affinity purification (SLF-AP), and proximity labelling (TurboID). B) Volcano plot of quantitative IP-mass spectrometry (IP-MS) of endogneous INTS12 (*n = 4*). C) Venn diagram showing the identified interactors in the TurboID, endogenous and SLF data sets and the common factors. D) Schematic of the INTS1, NELFE and SPT5 IP-MS performed after 2h of INTS12 degradation. E) Volcano plot of quantitative IP-MS of endogenous INTS1 in prescense of dTAG^V^-1 or vehicle (2h, *n = 4)*. F) Lollipop diagram of INTS1, NELFE and SPT5 IP-MS highlighting Integrator, NELF, SOSS and PP2A components upon INTS12 degradation (2h). G) Immunoblots of the INTS1 IP-MS -/+ dTAG^V^-1 (250 nM) probing for INTS1, INTS3, INTS12 and NELFA *(n = 2)*. H) Immunoblots of whole lysate glycerol gradient samples from INTS12-FKBP12^F36V^ cells treated with dTAG^V^-1 (2h) then probed for NELFA, INTS12, INTS1, RNAPII (4H8) and SPT5 (*n = 2*).

Analysis of the resulting mass spectrometry (MS) data revealed the co-purification of the entire NELF complex, comprising NELFA, NELFB, NELFCD, and NELFE, as well as the entire Integrator complex. The PP2A holoenzyme, comprising PPP2R1A and PPP2CA, and RNAPII subunits were identified in the endogenous IP and SLF-biotin AP, but not in the TurboID-based proximity labelling, possibly owing to the positioning of the TurboID tag, physical distance from the INTS12 protein, or steric hindrance to the proximity labelling (**Figure 2B and C**). Proximity labelling additionally indicated the proximity between INTS12 and numerous factors involved in pause-release and the RNAPII transcription cycle more broadly such as components of Mediator, the super-elongation complex (SEC), and transcriptional CDKs (**Supplementary Figure 2D**).

To determine the impact of INTS12 degradation on the Integrator complex and its interactors, we performed IPs using antibodies against INTS1, NELFE, and SPT5 in cells treated with dTAG^V^-1 for 2 hours (**Figure 2D**). The IP-MS results demonstrated the strong and specific enrichment of each of these components relative to a matched IgG control (**Supplementary Figure 2E, F and G**). Statistical comparison between the dTAG^V^-1 treated condition and the DMSO control in the INTS1 IP-MS data using the LIMPA analysis suite, revealed that INTS12 depletion led to the selective loss of the entire NELF complex from Integrator (**Figure 2E**). INTS12 loss did not affect the co-purification of other Integrator components with INTS1 (**Figure 2E and Figure 2F**), nor did it significantly impact its association with auxiliary subunits including SOSS and PP2A (**Figure 2E and F**), suggesting that the Integrator complex remains intact. SPT5 IP-MS data mirrored INTS1 albeit less clearly, possibly owing to its facultative association with Integrator (**Figure 2F and Supplementary Figure 2F**). With respect to NELFE; however, binding was reduced with all Integrator components and the PP2A submodule (**Figure 2F and Supplementary Figure 2G**). We next validated the INTS12-dependent interaction between NELF and Integrator by IP-Immunoblot in eHAP1 cells, which confirmed the reduced association between these two complexes in the absence of INTS12 (**Figure 2G**). This direct association is supported by crosslinks detected between INTS12, INTS1, and the NELF complex (NELFA in particular) in re-analysed cross-linking MS data from (13) (**Supplementary Figure 2H**). Furthermore, assessment of the structural integrity of the Integrator complex using density based separation revealed no major shifts in the migration of the integrator complex upon INTS12 degradation (fractions 6-11), although a modest increase in the smaller INTS1-comprising complexes could be observed (fractions 6-8). Under the same conditions, NELFA was lost from the largest complexes (fraction 11) where it co-migrates with RNAPII and Integrator, with no changes seen for SPT5 in those fractions (**Figure 2H**).

Together, these data indicate that INTS12 is required for the stable association between the Integrator complex and NELF. Acute loss of INTS12 results in a rapid dissociation between the intact Integrator and the NELF complex. Despite INTS12 having no defined structure in current Integrator complex structures, using the crosslinking data we predict that INTS12 interacts strongly with the N-terminal region of INTS1 and resides between INTS1 and the NELF complex (**Supplementary Figure 2I**).

### The INTS12 PHD domain mediates its interaction with NELFA and is required for its function in promoter-proximal transcription

In evolutionary terms, INTS12 has been described to be the youngest member of the Integrator complex; it contains a single defined central domain, called the plant homeodomain (PHD) (33).To generate insights into the importance of the INTS12 protein PHD domain and adjacent regions, we looked through an evolutionary lens by comparing its sequence similarity across orthologs from *Chordata* to *Placozoa,* as well as with other PHD domain containing genes (*PHF1*, *PHF19* and *MTF2)* (**Figure 3A**). This analysis revealed that ancestral versions of INTS12 can be found dating back to *Placozoa*. Clear sequence similarity and conserved domain organisation can be observed between the human protein sequence and INTS12 in *Spiralia*, including its PHD domain, the N-terminus and a small fraction of the C-terminus regions. Sequence similarity was less apparent with other PHD domain containing proteins, even within this subdomain itself, suggesting early divergence in function (**Figure 3A**). The importance of these three distinct subdomains of INTS12 is further underscored by analysis of the rate of evolutionary change (rate4seq) using MSA alignments as determined in the predictomes database (34), which demonstrates resistance to changes in these regions over time. Reciprocally, AlphaMissense analysis (35) showed a higher likelihood for mutations in these regions to be deleterious to INTS12 function (**Supplementary Figure 3A**). Predictomes analysis correctly identified that the N-terminal domain of INTS12 mediates an interaction with INTS1 via two small α-helices making up this region, which is in line with previously reported observations from the Wagner laboratory (31) (**Supplementary Figure 3B**). A second interface was predicted between the INTS12 PHD domain and the C-terminus of NELFA, suggesting a direct interaction between INTS12 and this component of the NELF complex (**Supplementary Figure 3B**). The same interface was predicted with the human protein-protein interaction database although with less confidence (**Supplementary Figure 3B**).

**Figure 3:**
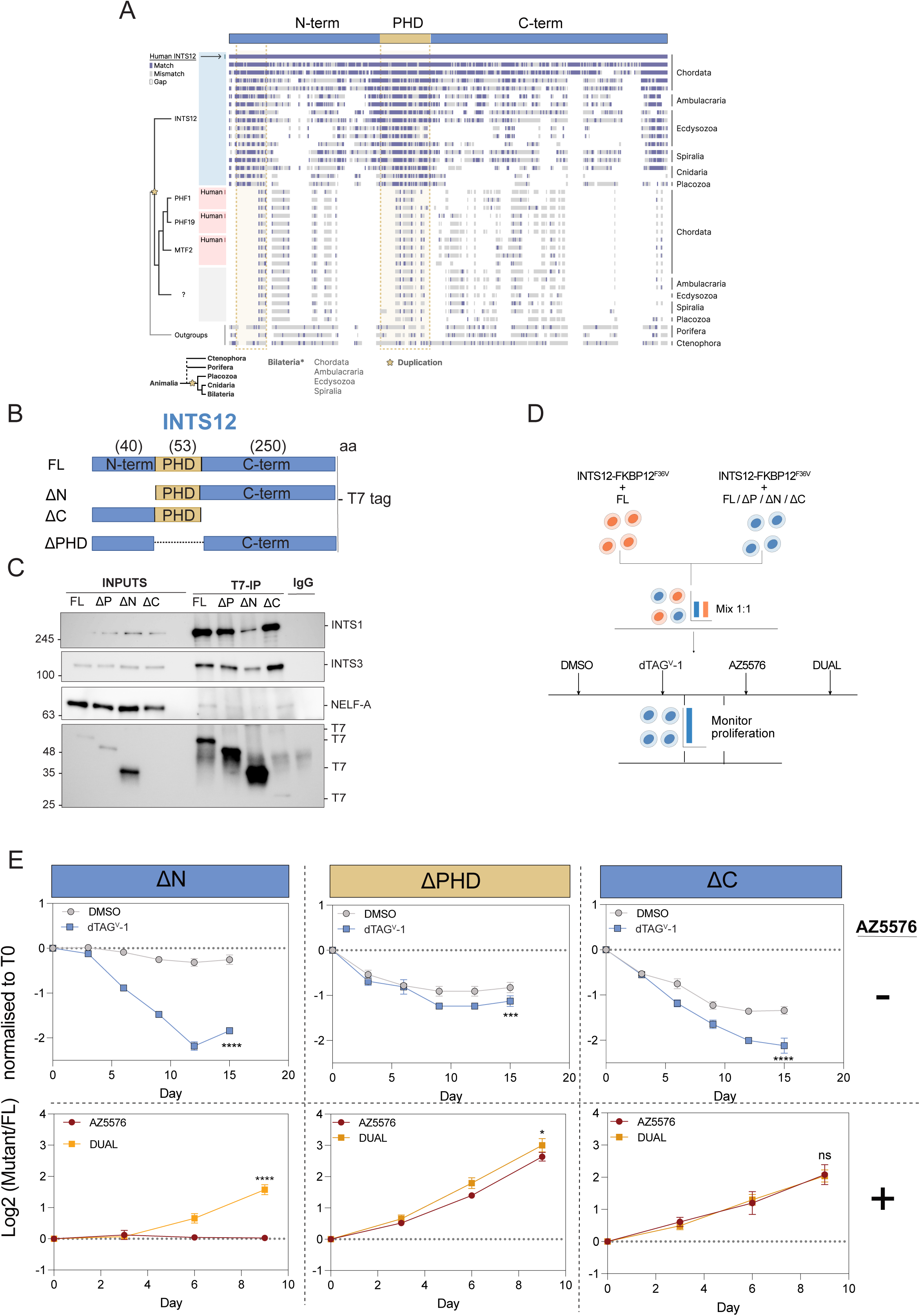
The INTS12 PHD domain mediates its interaction with NELFA is required for its function in promoter-proximal transcription. A) Alignment of INTS12 homolog sequences. Horizontal bars represent protein sequences that have been aligned to human INTS12. Each band represents individual amino acids; Blue bands indicate an exact amino acid match, grey bands indicate a mismatch. Empty space indicates a gap in the alignment. The black outline above the alignment represents a simplified diagram of human INTS12, showing the positions of the N-terminal and PHD domains as yellow regions. The positions of these regions have been extended downwards using broken yellow lines for easier reference. Human sequences in the alignment are labelled on the left; The inferred evolutionary relationship between each of these sequences is illustrated as a tree. The question mark indicates all outgroup sequences to PHF1, PHF19, and MTF2. An inferred duplication event in this tree is marked with a yellow star. This duplication is also marked on a tree below the alignment, which shows its timing relative to the branching of major animal lineages. Labels on the right of the alignment indicate the animal lineage that each sequence belongs to. B) Schematic of the INTS12 domains and the protein mutants generated. N-term, N-terminal region; C-term, C-terminal region; FL, full length, ΔN, deletion of N-terminal region; ΔC, deletion of C-terminal region; ΔPHD (or ΔP), deletion of PHD domain. C) Immunoblots of samples from the T7-IP probed for INTS1, INTS3, NELFA and T7 (*n = 4*). D) Schematic of the competitive proliferative assay between the INTS12-FKBP12^F36V^ cells with FL overexpression put in competition 1:1 with the INTS12-FKBP12^F36V^ cells with the overexpression of the mutants. E) Competitive proliferation assays result of the mutants -/+ dTAG^V^-1 (top panels) (*n = 2*) and -/+ AZ5576 and DUAL (bottom panels) (*n = 3*). *p < 0.05, ***p < 0.0005, ^∗∗∗∗^p < 0.0001, ns = non significant; 2-way ANOVA.

To determine the molecular and functional importance of these domains for INTS12 biology, we generated deletion mutants of the N-terminal, C-terminal, and PHD domains fused to the T7 epitope tag (**Figure 3B**). We then stably reconstituted these mutants in the INTS12 degron-tagged eHAP1 cells (**Supplementary Figure 3C**). Firstly, we sought to examine the importance of each of these domains for the interaction between INTS12, INTS1, and NELFA by IP of the T7-tag. The endogenous protein was degraded for 2 hours using dTAG^V^-1 (250 nM) to prevent competition in binding between mutants and endogenous wild-type protein and IP of the T7-tagged exogenous constructs was then performed.. This demonstrated that deletion of the N-terminal region of INTS12 nearly completely abrogated its ability to interact with INTS1 and NELFA (**Figure 3C**). As this domain is not predicted to mediate interactions with the NELF complex directly, this observation suggests that association with Integrator precedes, and is required for, the interaction between INTS12 and NELFA. Deletion of the PHD domain in contrast did not affect the ability for INTS12 to interact with INTS1, however resulted in reduced association with NELFA. No discernible defect in binding between INTS12 and either INTS1 or NELFA was observed for the C-terminal deletion mutant, despite its evolutionary conservation, likely indicating that it plays key roles in other aspects of INTS12 biology (**Figure 3C**). To determine the functional impact of the deletion mutants, we next performed a series of genetic complementation assays in INTS12 degron-tagged eHAP1 cells, leveraging the INTS12 essentiality and resistance phenotypes to attempt to separate functions. The eHAP1 cells carrying deletion mutants were placed in direct competition with eHAP1 cells expressing full-length exogenous INTS12 in the presence or absence of dTAG^V^-1 and/or the CDK9 inhibitor (**Figure 3D**). In line with the IP data results, the INTS12 N-terminal deletion mutant failed to rescue the depletion of the endogenous protein and was rapidly outcompeted by the full-length rescue line (**Figure 3E top left**). In concordance, the N-terminal mutant behaved as a null allele in the presence of CDK9 inhibition, resulting in the rapid and conditional outgrowth of the N-terminal mutant in the presence of dTAG^V^-1 and AZ5576, but not when treated only with AZ5576 where the wild-type endogenous protein is still expressed (**Figure 3E bottom left**). In contrast, the PHD deletion mutant negatively impacted cell growth even in the presence of the wild-type INTS12 protein, indicated by the steady loss of this condition in both the DMSO and dTAG^V^-1 treated conditions, suggesting that it may act as a dominant negative form (**Figure 3E top centre**). Indeed, expression of the PHD-deletion mutant mediated resistance to CDK9 inhibition even in the presence of the wild-type protein as compared to the full-length control lines (**Figure 3E bottom centre**). This further underscores its dominant negative activity, and highlights a critical role for the INTS12 PHD domain in CDK9-regulated transcription cycle checkpoints. Although we could not find any interaction defects with the C-terminal deletion mutant, it largely phenocopied the PHD-deletion mutant in these assays (**Figure 3E right**), indicating that it must play a role in promoter-proximal regulation of Integrator. These observations clearly pinpoint INTS12 as the central bridge between Integrator and NELF, via interactions between its N-terminal and PHD domains with INTS1 and NELFA, respectively.

### INTS12 is required for the recruitment and stable formation of the NELF-Integrator complex on chromatin

The Integrator complex has recently been described to exist in multiple diverse states: A “paused” or pre-termination state as part of a NELF/RNAPII complex on chromatin, a post-termination state that lacks DSIF and NELF, and a “free” state where the integrator complex is no longer tethered to chromatin (13). To determine how acute degradation of INTS12 impacts the occupancy of the constituents of these complexes on chromatin, we performed chromatin-immunoprecipitation sequencing (ChIP-seq) on INTS12, INTS1, NELFE, NELFC/D, pan-phospho (4H8) and Ser2^P^ forms of RNAPII (**Figure 4A**).

**Figure 4:**
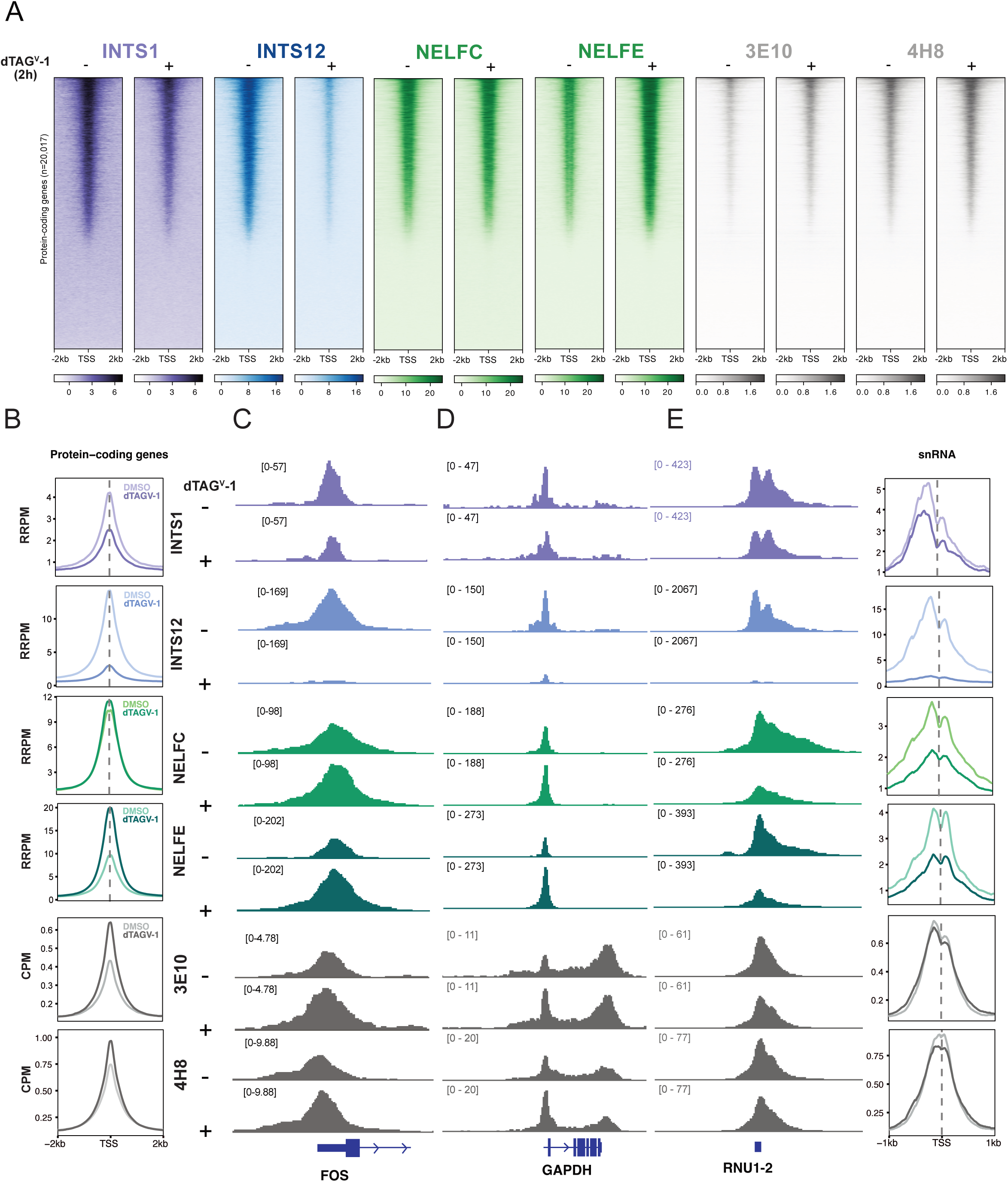
INTS12 is required for the recruitment and stable formation of the NELF-Integrator complex on chromatin. A) Tornado plots depicting the occupancy of INTS1, INTS12, NELFC, NELFE and RNAPII (3E10 and 4H8) in eHAP1 cells treated with DMSO (-) or dTAG^V^-1 (+) to induce INTS12 degradation. ChIP-seq signal is centred on the TSS of protein-coding genes (*n = 20,017*), with TSSs defined as the gene level start coordinates. B) Average ChIP-seq profiles centred on transcript level TSSs of protein-coding genes(+/- dTAG^V^-1). TSSs were defined as the transcript level start. C) Genome browser view of ChIP-seq tracks for each probed factor at the *FOS* D) and *GAPDH* loci. E) Average ChIP-seq profiles centred on transcript-level TSSs of snRNA genes (*n = 1,901*, right panel) with genome browser view of ChIP-seq tracks at the RNU1-2 locus (left panel). RRPM, reference adjusted reads per million; CPM, counts per million. One representative replicate is shown for each mark.

The acute degradation of INTS12 resulted in the global loss of its binding at all TSS-proximal regions in the genome (**Figure 4B-E**). INTS1 occupancy was also reduced although to a lesser extent than INTS12 itself (**Figure 4B**), which is possibly due to interactions with other chromatin-bound interfaces including those on RNAPII. This suggests that loss of INTS1 in the absence of INTS12 is the result of reduced *de novo* association with RNAPII, or the eviction of paused-Integrator forms from TSS-proximal regions rather than the immediate dissociation of already assembled paused-Integrator complexes. In contrast to INTS1, NELFE binding to chromatin was markedly increased in the absence of INTS12 at TSS-proximal regions for protein coding genes, with occupancy for NELFC/D largely unaffected (**Figure 4B and Supplementary Figure 4B**). Conversely, an opposing trend was noted for snRNA loci where both NELFC/D and NELFE were lost from chromatin (**Figure 4E**).

These data suggest that Integrator complexes differ in composition. More likely the absence of INTS12 leads to NELF retention on chromatin as a result of conversion to a canonical paused state with DSIF (4, 36) or through electrostatic interactions with unprocessed RNAPII-bound nascent RNA (37, 38). Finally, in line with early Immunoblots (**Figure 1H**), the acute degradation of INTS12 resulted in increased TSS-proximal signal for the pan-phospho RNAPII and to a greater extent Ser2^P^ (**Figure 4B-D and Supplementary Figure 4B**). Together with the proteomics data, these findings demonstrate that INTS12 is required for the recruitment of Integrator to NELF-bound RNAPII, and that its acute loss results in eviction of the intact Integrator complex but not NELF from chromatin.

### INTS12 degradation drives the release of defective, non-processive RNAPII complexes

To determine the consequence of INTS12 degradation on nascent transcription we performed transient transcriptome sequencing (TT-seq). INTS12 was degraded for 1.5 hours with 4-thiouridine labelling in the last 15 minutes to capture nascent RNA (**Figure 5A**). Under homeostatic, asynchronous conditions, INTS12 degradation resulted in a modest increase of signal at the 5’-end of the transcriptional unit (TU) and a slight but reproducible reduction at the 3’-end (**Figure 5B**), indicative of the increased release of RNAPII complexes, that are unable to effectively complete the transcription cycle. To quantify this observation more accurately, we calculated a processivity index by taking the ratio the TT-seq signal of the first 1.25kb of the TU from TSS and the first 1.25kb from the TES. This revealed a robust and statistically significant increase in the processivity index, indicative of a diminished capacity for RNAPII to make it to the end of the transcription cycle (**Figure 5B and C, and Supplementary Figure 5A and B**). To investigate the impact of INTS12 depletion on promoter-proximal release of RNAPII, we next analysed TT-seq data under the same conditions as before, adding AZ5576 or the vehicle in the final 30min of the treatment. CDK9 inhibition prevented the release of RNAPII from TSS-proximal regions, but left RNAPII complexes past this point unaffected. This resulted in a characteristic retreating-wave signal in TT-seq, which is proportional to the rate at which RNAPII complexes clear the promoter-proximal regions in the genome (**Figure 5D-F**). Notably, prior INTS12 depletion impaired the ability of the CDK9 inhibitor to enforce transcriptional pausing. This is indicated by the continued release of RNAPII into the elongation phase as evidenced by TSS-proximal signal in the dTAG^V^-1 plus CDK9i treated condition (**Figure 5D**) and an increase in the processivity index (**Figure 5E** and **Supplementary Figure 5C and D**).

**Figure 5:**
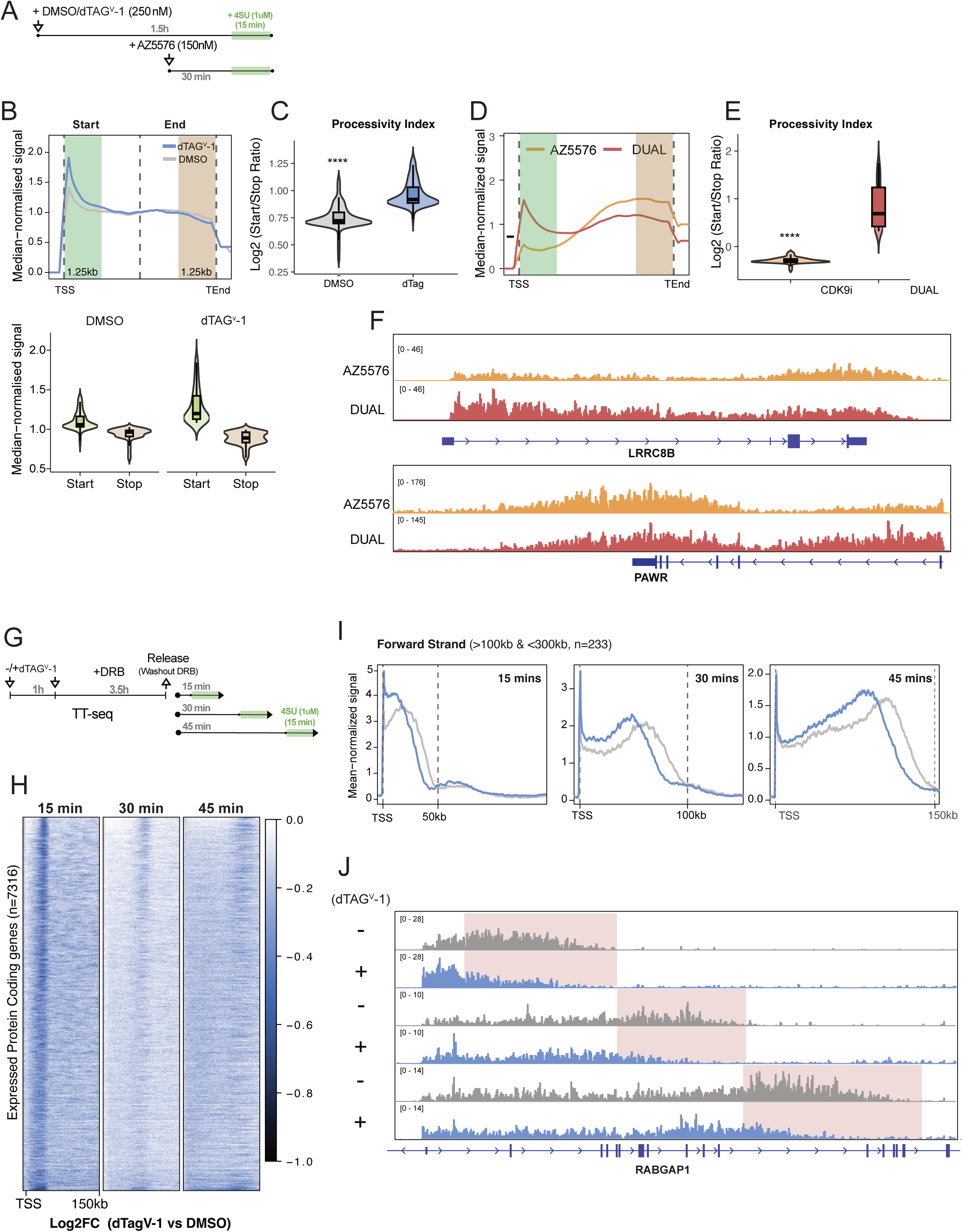
INTS12 degradation drives the release of defective, non-processive RNAPII complexes. A) Schematic of TT-seq experiments performed in eHAP1 cells treated with DMSO or dTAG^V^-1. The green box highlights the addition of 4SU. B) Median-normalised metagene profiles of TT-seq signal across protein-coding genes (*n =1458*; gene length >100kb and <1Mb). The gene body was scaled to 5kb, with a 1.25kb window highlighted around the TSS and TES. The highlighted regions are plotted in the lower panel, and C) The processivity index is expressed as log2(start/stop ratio); ^∗∗∗∗^p < 0.0001 paired T-test. D) Equivalent analyses are shown for CDK9i and dual (CDK9i + dTAG^V^-1) treatments (*n = 1,494*; gene length >100kb and <1Mb), with a 1.25 kb highlighted window at the TSS and TES and E) the processivity index is shown; ^∗∗∗∗^p < 0.0001 paired T-test. F) Genome browser view of TT-seq signal at *LRRC8B* locus on the forward strand and *PAWR* on the reverse strand in CDK9i and dual (CDK9i + dTAG^V^-1) treated cells. G) DRB-TT-seq schematic; arrows mark the addition of the different compounds, and release after washing out DRB. Samples are collected at each of the denoted timepoints after the addition of 4SU in the last 15 mins. H) Heatmap showing genome-wide Log2FC(dTAG^V^-1/DMSO) signal at each timepoint for all expressed protein coding genes (*n = 7,316*). I) Mean normalised DRB-TT-seq profile plots centered at TSSs; dashed lines indicate signal extension to 50kb, 100kb and 150kb at 15, 30 and 45 mins, respectively. J) Genome browser views for DRB TT-seq signal at 15, 30 and 45 mins. Boxes highlight regions exhibiting reduced processivity.

As transcriptional defects can be obscured in the asynchronous state, we next sought to determine the impact of INTS12 depletion after transcription cycle synchronisation with DRB. To this end, we treated INTS12 degron-tagged lines with dTAG^V^-1 for 1h, followed by DRB treatment for 3.5h, followed by the timed release for 15, 30, and 45 min with labelling in the final 15min with 4sU (**Figure 5G** and **Supplementary Figure 5E and F**). The DRB-release TT-seq experiments revealed that INTS12 loss resulted in a profound and global loss in processivity of RNAPII as compared to vehicle treated control conditions. This is evidenced by the decreased signal at 3’-end loci in dTAG^V^-1 conditions compared to DMSO controls, which was observed at each of the time points post-release at loci comprising all expressed protein coding genes (**Figure 5H**). Visualisation of protein-coding genes between 100kb and 300kb confirmed this phenotype with consistently reduced processivity at each of the timepoints (**Figure 5I and J)**. Taken together, these findings indicate that in response to INTS12 loss and concomitant reduced paused-Integrator stability, RNAPII can be released into defective, non-processive elongation, likely from a canonical or aberrant NELF-bound state.

### INTS12 loss genetically interacts with pause-release factors, Integrator, and the ARMC5 degradation pathway

To contextualise these observations in a broader genetic context we decided to profile the synthetic lethality and resistance following INTS12 degradation in genome-wide CRISPR-screens. We introduced Cas9 into the INTS12 degron-tagged eHAP1 cells and subsequently transduced these with a bespoke PROTAC-compatible CRISPR-library lacking sgRNAs against Cereblon, VHL, and other mediators of dTAG/PROTAC-induced degradation. Following transduction and selection with hygromycin, cells were cultured in the presence or absence of either DMSO or dTAG^V^-1 for 20 days after which the guide abundance was analysed relative to a T0 control for each of the treatment conditions as well as to each other (**Figure 6A**). This screen revealed that INTS12 loss was synthetically lethal with inactivation of INTS1, ARMC5, and components of the CDK9 sequestering HEXIM1 complex (**Figure 6B and Supplementary Figure 6A,B)**. Indeed, validation of ARMC5 and HEXIM1 in competition assays revealed that KO lines comprising these deletions were rapidly outcompeted by their AAVS sgRNA counterparts only when INTS12 is degraded (**Figure 6C-E**). These observations fit previous publications demonstrating synthetic lethal interaction between Integrator dysfunction and ARMC5 (16, 17) and suggest that increased activity of CDK9 mediated pause-release is toxic in the absence of INTS12, likely owing to the release of defective, non-processive RNAPII inducing a cellular stress response. In agreement with this observation, deletion of ELL and MED8 resulted in desensitisation to INTS12 depletion (**Figure 6D-E),** possibly due to reduced defective elongation. The later is in line with the CDK9 inhibitor treatment results, and it could be equally argued that diminished CDK9 activity enhances the cells tolerance to INTS12 depletion. In addition, analysis of public dataset from the Dependency Map (DepMap) portal showed a positive correlation between INTS12, NELFA and HEXIM1, albeit moderate, and a negative correlation between INTS12 and ELL (**Supplementary Figure 6C**). Finally, the synthetic lethality with INTS1 is noteworthy as one would consider that their phenotypes would be overlapping, either pointing to additive effects or additional INTS12-independent functions for Integrator, possibly in its free or post-termination state.

**Figure 6:**
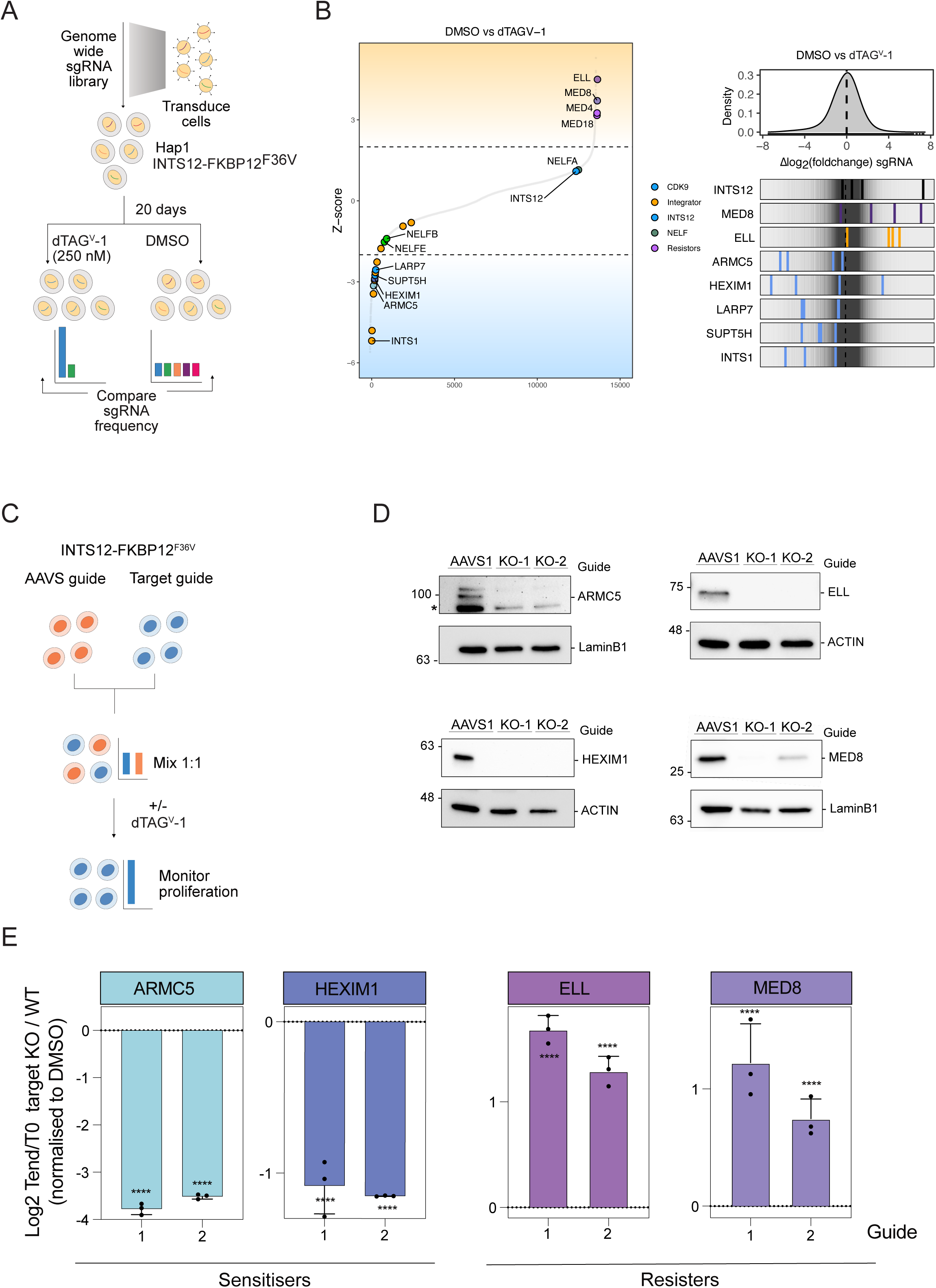
INTS12 loss genetically interacts with pause-release factors, Integrator and the ARMC5 degradation pathway. A) Schematic of the CRISPR-Cas9 genome-wide screen: INTS12-FKBP12^F36V^ eHAP1 cells were transduced with the PROTAC-compatible CRISPR sgRNA library and cultured with DMSO or dTAG^V^-1 for 20 days. B) Enriched and depleted sgRNAs from the screen (left side) and their abundance (right side). C) Schematic of the competitive proliferation assay for (E); INTS12-FKBP12^F36V^ cells nucleofected with AAVS1 sgRNA and INTS12-FKBP12^F36V^ cells nucleofected with either ARMC5, ELL, HEXIM1 or MED8 guides (2 different guides/gene) were mixed 1:1 and treated with DMSO or dTAG^V^-1. D) Immunoblots from INTS12-FKBP12^F36V^ cells nucleofected with AAVS1 sgRNA and INTS12-FKBP12^F36V^ cells nucleofected with either ARMC5, ELL, HEXIM1 or MED8 guides (2 guides/gene) (n = 2; * represents non-specific bands). E) Bar plots showing the results of the competition assays for ARMC5, ELL, HEXIM1 and MED8. Data shown is Tend normalised to T0 and dTAG^V^-1 condition normalised to DMSO This result is representative of *n = 4* replicates. ****p < 0.0001.

Taken together, our observations clearly define the role of INTS12 in Integrator recruitment to the RNAPII-NELF complex, elucidate the reversible nature of this state and unravel an intriguing interplay with the canonical CDK9 mediated pause-release.

## Discussion

Here, we reveal that INTS12 mediates the association between Integrator and NELF-bound RNAPII to stabilise the “paused” form of Integrator. The acute depletion of INTS12 abrogates effective recruitment and destabilises this complex resulting in the eviction of the entire Integrator complex from RNAPII and NELF. Disruption of paused Integrator increases the release of RNAPII complexes from TSS-proximal paused-states into elongation (**Figure 7**). However, these released complexes are largely non-processive, as evidenced by diminished occupancy toward the distal 5’ end of the transcriptional unit. Indeed, the disruption of paused-Integrator via INTS12 degradation led to the entire transcriptionally engaged RNAPII pool becoming non-processive in time-series DRB-release experiments. In contrast to the asynchronous setting, this suggests that paused forms of RNAPII go through an additional INTS12/Integrator mediated pausing step after release by CDK9. Alternatively, it is possible that prolonged DRB exposure increases the ratio between the canonical paused RNAPII state and the Integrator paused form, thus resulting in a near complete penetrance of the processivity defect. This raises important questions about the order of pausing events, the involvement of CDK9 in diverse states of paused RNAPII and the possible interconversion between these states. The observation that DRB sensitive RNAPII complexes can still be impacted by INTS12 depletion would minimally suggest that there is a crosstalk between the Integrator paused state and the canonical form and may indicate that the Integrator quality control step can arise after classical paused forms. In agreement with this extensive crosstalk, we observe that the depletion of INTS12 confers potent resistance to the catalytic inhibition of CDK9. One possible explanation is that CDK9 plays a role in the release of Integrator from NELF. To date, the biochemical and structural evidence for this direct model is lacking.

**Figure 7:**
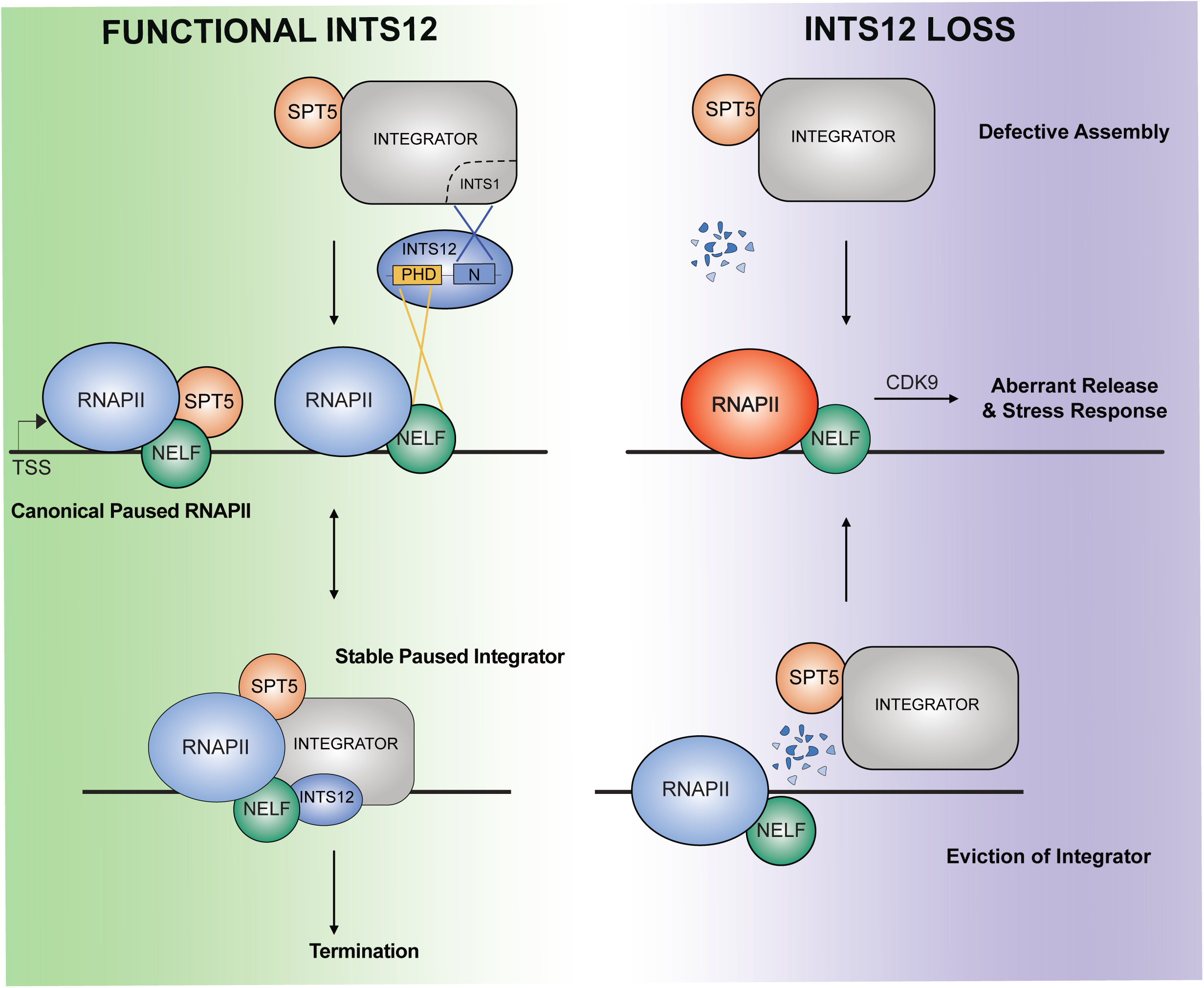
INTS12 loss results in release of aberrant RNAPII molecules into elongation. INTS12 binds to the Integrator complex through direct interaction with INTS1. Further, INTS12 associates with the NELF complex via its PHD domain by direct interaction with NELFA and possibly other interactors. These INTS12 interactions result in a stable paused complex prior to Integrator-mediated termination. When INTS12 is depleted, either the de novo recruitment of Integrator to the RNAPII-NELF complexes is perturbed or INTS12 loss causes the disruption of already chromatin-bound paused complexes. Regardless of the type of disruption, loss of INTS12 results in increased RNAPII-NELF association on chromatin, which exposes this improperly assembled complex to CDK9-dependent release from pausing. This presumed aberrant release leads to non-processive RNAPII molecules that fail to reach the end of genes, thus driving a cellular stress response, paralleling INTS1 loss (27).

An alternative mode of action could be that the loss of paused-Integrator results in the conversion of the now exposed RNAPII interface to its canonical paused form from which can then be released by CDK9. Indeed, we observe that the depletion of INTS12 does not result in a dissociation of NELF from chromatin at protein coding regions. How NELF remains bound to chromatin needs further investigation: its binding may be mediated via its association with RNAPII in the absence of DSIF or alternatively it may be facilitated by electrostatic interactions with promoter-proximal RNA which persists in the absence of Integrator-mediated cleavage.

Indeed, several NELF components have been demonstrated to have RNA binding affinity including NELFA and NELFE (37–39). Although we did not see an increased association of NELF with SPT5 by mass-spectrometry, NELF has been shown to interact more with SPT5 upon loss of INTS1, providing evidence that interconversion is possible when the entire Integrator complex is perturbed (27). A final explanation could be that there is activity *in trans* whereby paused Integrator complexes act upon canonical paused RNAPII in TSS proximal regions via the phosphatase module. Our work also highlights a difference in the RNAPII-NELF complexes composition and INTS12 dependency at snRNA loci versus mRNA loci, with the removal of INTS12 mediating the near complete loss of NELF occupancy on chromatin at the former. The acute depletion of NELF has been demonstrated to cause RNAPII accumulation close to the canonical paused form, a state that is insensitive to CDK9 and may be mediated by Integrator (6). The relationship between CDK9 and INTS12 can also be viewed as reciprocal with reduced release of RNAPII through CDK9 inhibition compensating for the excessive release of aberrant RNAPII complexes upon INTS12 depletion. In agreement with this observation, we find that the knock-out of ELL conferred moderate resistance to INTS12 depletion, with HEXIM1 loss conferring sensitisation, presumably through decreased or increased CDK9 activity, respectively. The Genome-wide CRISPR screening confirmed recent literature on the ARMC5 salvage pathway and additionally demonstrated that core Integrator components are the key sensitisers in INTS12 depleted cells (16, 17). This may indicate that the free Integrator or non-paused forms of the complex play critical roles in gene regulation.

We demonstrate that the interactions between INTS12 and Integrator are mediated by its conserved N-terminal domain via interactions with INTS1, in agreement with previous observations from the Wagner laboratory (31). The INTS12 PHD domain is required for stable binding to the NELF complex via an interface with NELFA, which is supported by biochemical evidence and modelling. PHD domains classically have been viewed as histone binding domains, but they have recently been demonstrated in non-histone and DNA binding (40, 41). The INTS12 PHD domain thus appears to have acquired divergent functions compared to the PHD domains of proteins such as PHF1, PHF19 and MTF2 which bind to methylated lysines on histone tails (42). The functional importance of the N-terminus and PHD domain is underscored by our genetic complementation experiments which revealed a clear dichotomy between these two domains. The N-terminal deletion was unable to compensate for the loss of INTS12 upon dTAG^V^-1 treatment in stably growing cells and did not confer re-sensitisation to CDK9 inhibition, indicative of a complete loss of function, fitting with the importance of the N-terminal domain in the association between INTS12 and Integrator. The loss of NELF association with the N-terminal deletion mutant of INTS12, in the absence of evidence for direct association between this domain and NELF, suggests that the recruitment of INTS12 to Integrator precedes its association with NELF and paused RNAPII. Interestingly, the PHD domain deletions decreased association with NELF, but were able to functionally compensate for the loss of INTS12 in steady-state conditions, indicating that the N-terminus is possibly sufficient to rescue the growth deficiency caused by INTS12 degradation. This fits past observation focused on snRNA transcription (31). However, upon CDK9 inhibition, PHD deletion mutants acted as dominant negative forms, mediating resistance even in the presence of WT INTS12. These observations indicate that retention of the N-terminus allows delta PHD mutants to dock into the Integrator complex, but fails in stably associating these mutants with NELF and they are thus able to outcompete the WT protein. These data also provide evidence that the phenotypic impact caused by INTS12 depletion on proliferation mainly arises from deficiencies in snRNA transcription.

INTS12 degradation resulted in the induction of a cellular stress response concomitant with a pro-inflammatory NFkB signature, similar to what has been observed upon depletion of INTS1 (27). Despite the similarity in transcriptional response, INTS12 has not been linked to the same developmental disorders that have been associated with INTS1 and INTS8. There is evidence that INTS12 can play a role in other disease such as COPD as evidenced by GWAS associations and the modulation of inflammatory signals (30). Our findings suggest that disruption of Integrator-NELF pausing may be at the heart of transcriptional deficiencies linked to human diseases, and possibly of resistance to targeted therapeutic exploiting this process. Taken together, our work uncovers the functional importance of the paused-Integrator complex with a key role for INTS12 herein.

## Limitations of the study

Our work is based on cell line models and steady state transcription. Although powerful for their reproducibility and ease of access to material, these models do not always capture the full complexity of gene regulation in a dynamic developmental or non-homeostatic condition. Moreover, we primarily focused on the acute rapid depletion of INTS12 and paused-Integrator, but have not evaluated compensatory functions upon prolonged depletion. Future work is necessary to confirm the role of INTS12 in human disease and development, using separation of function mutants such as the PHD deletion variants to distinguish between the impact on snRNA transcription and the regulation of protein coding genes.

## Supporting information

CRISPRscreen_sgRNAsummary

CRISPRscreen_genesummary

## Supplementary Data Figures

**Supplementary data Figure1:**
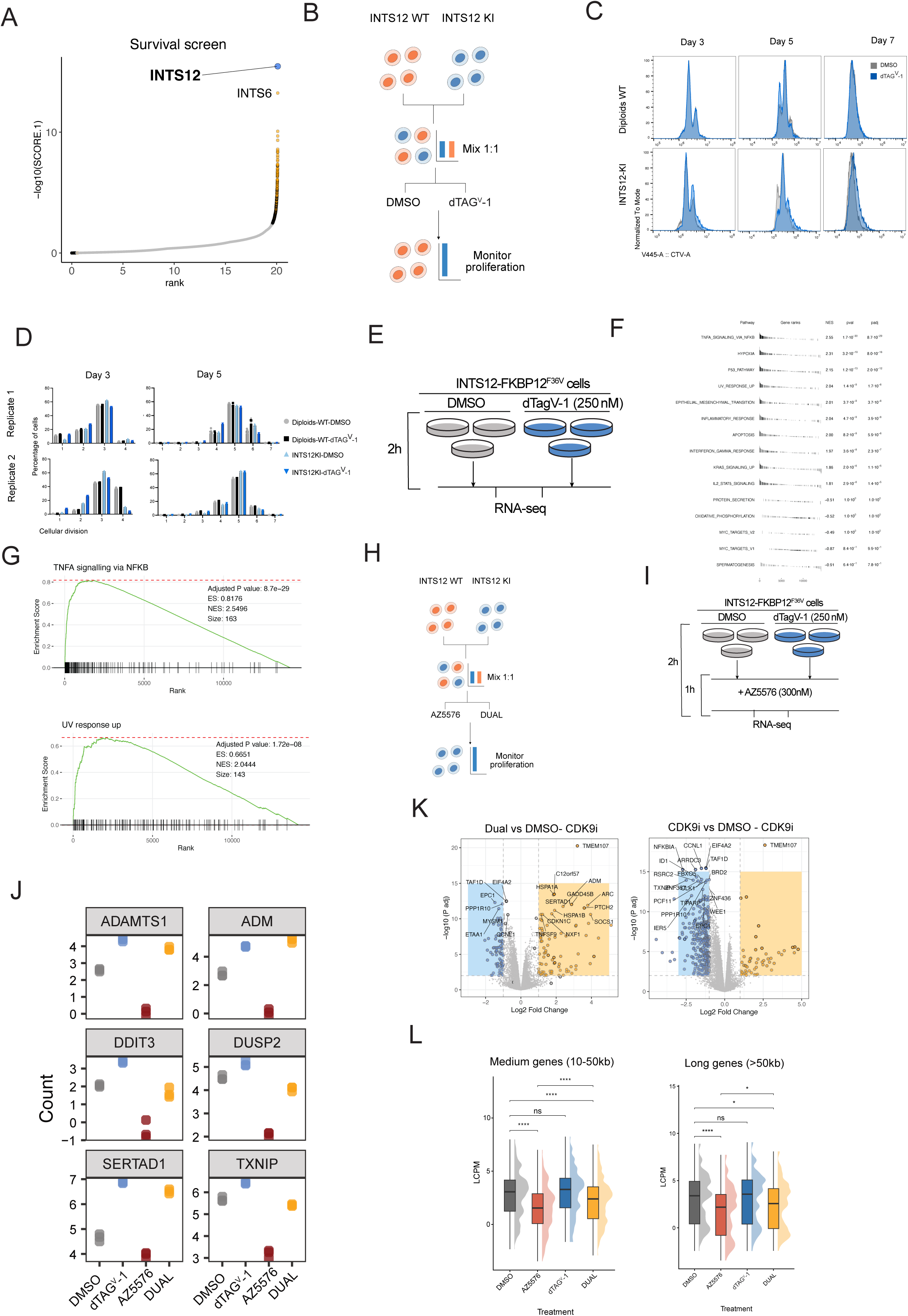
A) CRISPR screen result from Vervoort et al in THP1 cells highlighting INTS12 as a resistance hit to CDK9 inhibition with AZ5576. B) Schematic of competitive proliferation assay between INTS12 KI and WT cells with DMSO or dTAG^V^-1. C) Cellular division tracking experiments. WT diploid or INTS12-KI cells were stained with CellTrace Violet and sorted. Density plot of a representative replicate after sorting cells in presence of dTAG^V^-1 or vehicle on Day 3, 5 and 7 *n = 2*. D) Percentage of cells present at each cellular division in days 3 and 5. E) Schematic of RNA-seq performed in INTS12 KI line with DMSO or F) Top 10 Upregulated and top 5 downregulated Hallmark pathways upon INTS12 depletion. G) Gene set enrichment analysis (GSEA) showing INTS12 the upregulated hallmark pathways “TNFA signalling via NFKB” and “UV response up”. H) Schematic of competition assay between INTS12 KI and WT cells with AZ5576 or DUAL. I) Schematic of RNA-seq preformed in INTS12 KI line with AZ5576 or DUAL. J) Example genes that respond to INTS12 depletion and CDK9 inhibition. K) Volcano plot of RNA-seq samples treated with dTAGV-1 or vehicle and AZ5576. L) RNA-seq results with CDK9 inhibition and how this affects medium (10-50kb) and long (>50kb) genes.

**Supplementary data Figure 2:**
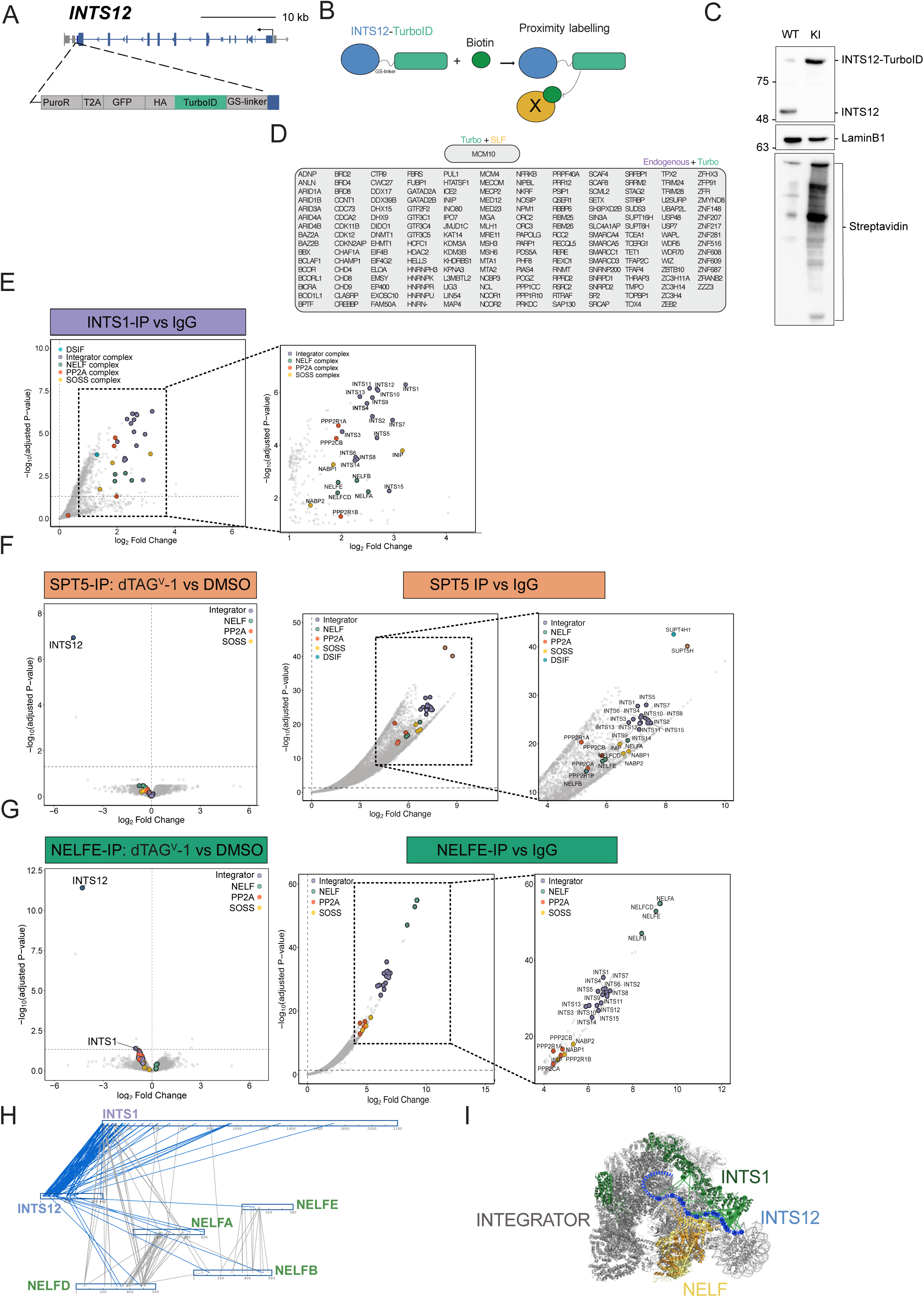
A) Schematic of INTS12-TurboID KI in the INTS12 locus. B) Schematic of mechanism of action of proximity labelling via TurboID. C) Immunoblotting showing WT eHAP1 or INTS12-TurboID tagged cells using INTS12 antibody (top panel). Labelling experiment showing streptavidin binding in INTS12-TurboID KI line. D) List of common INTS12 interactors between endogenous and TurboID or between SLF and the TurboID methods. E) Volcano plot of the INTS1 IP-MS experiment between IP samples and IgG. F) Volcano plot of the SPT5 IP-MS experiment between IP samples and IgG. G) Volcano plot of the NELFE IP-MS experiment between IP samples and IgG. H) Re-visualisation of XL-MS data from Fianu et al 2024 showing interactions between INTS12, INTS1 and NELF components. I) Mapping of INTS12 crosslinks to INTS1 and NELF in the Integrator complex (PDB 8RBX).

**Supplementary data Figure 3:**
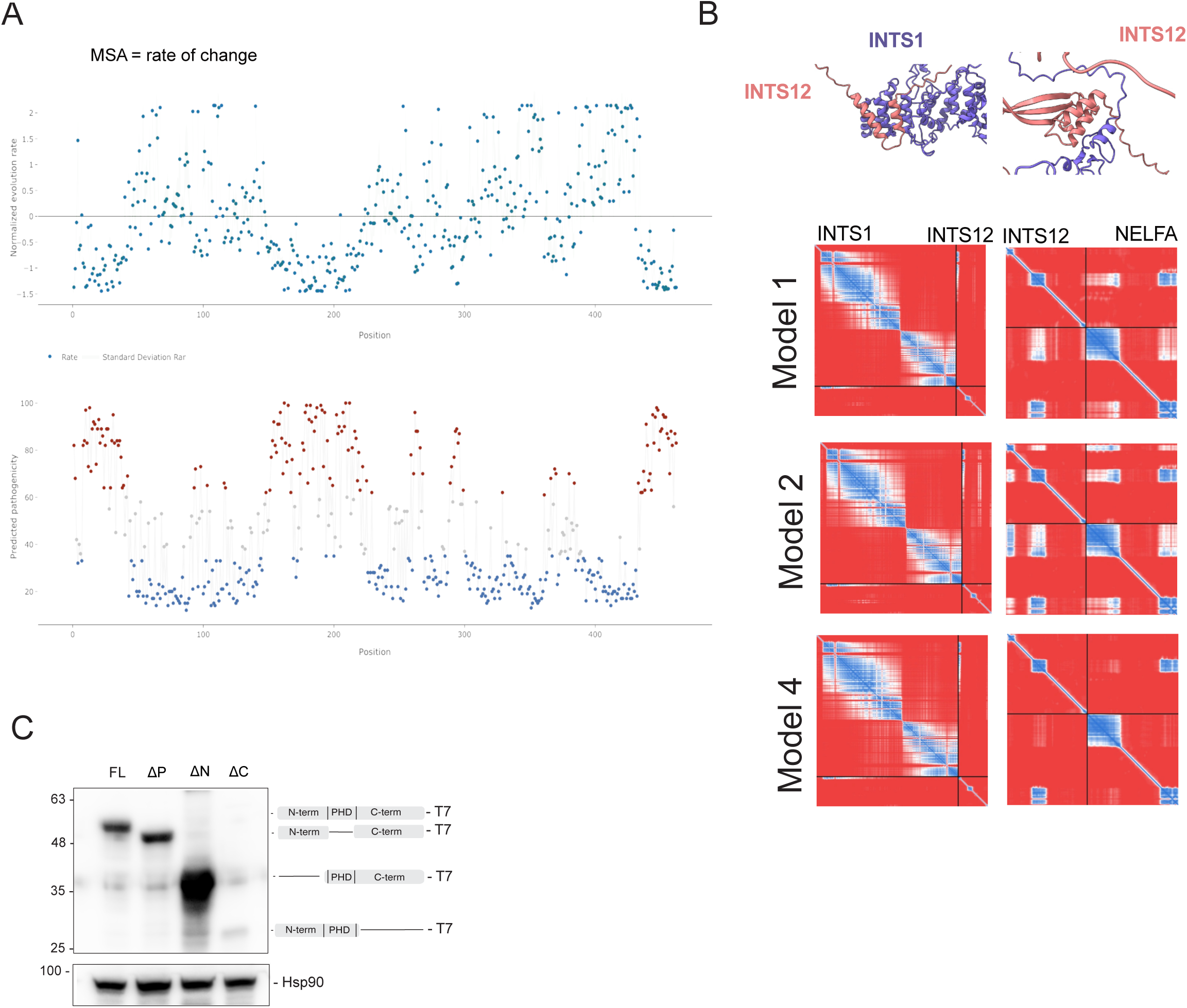
A) Scatter plots showing the evolution rate (top panel) and predicted pathogenicity (bottom panel) of the different regions of INTS12 protein. B) Prediction of interaction between INTS12 and INTS1 (left panels) and NELF (right panels). C) Immunoblotting of INTS12 over expressing mutants in the INTS12-FKBP KI line using T7 antibody.

**Supplementary data Figure 4:**
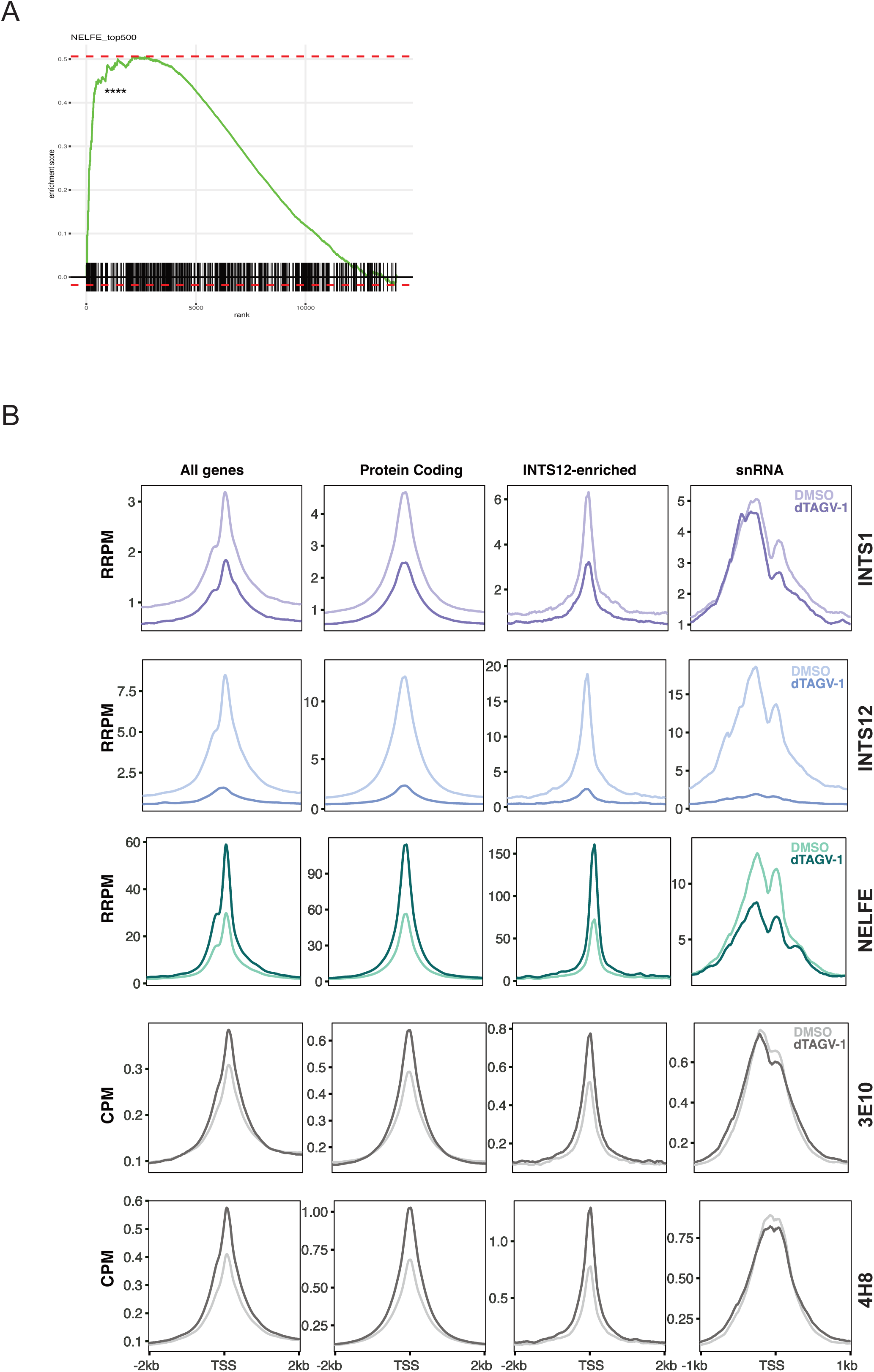
A) GSEA was performed on a ranked list of genes comparing dTAG^V^-1 treated and untreated (DMSO) samples. A gene set of the Top 500 NELFE bound sites identified from NELFE ChIP-seq signal in DMSO was defined. The vertical black bars indicate the position of genes belonging to the NELFE bound set. NES, Normalized Enrichment Score; \*\*\*\**p.adj* < 0.0001. B) Average profile plots of replicate 2 for INTS1, INTS12, NELFE, 3E10 and 4H8 ChIP-seq with DMSO and dTAG^V^-1 for all genes, protein coding genes, INTS12-enriched genes or snRNA genes.

**Supplementary data Figure 5:**
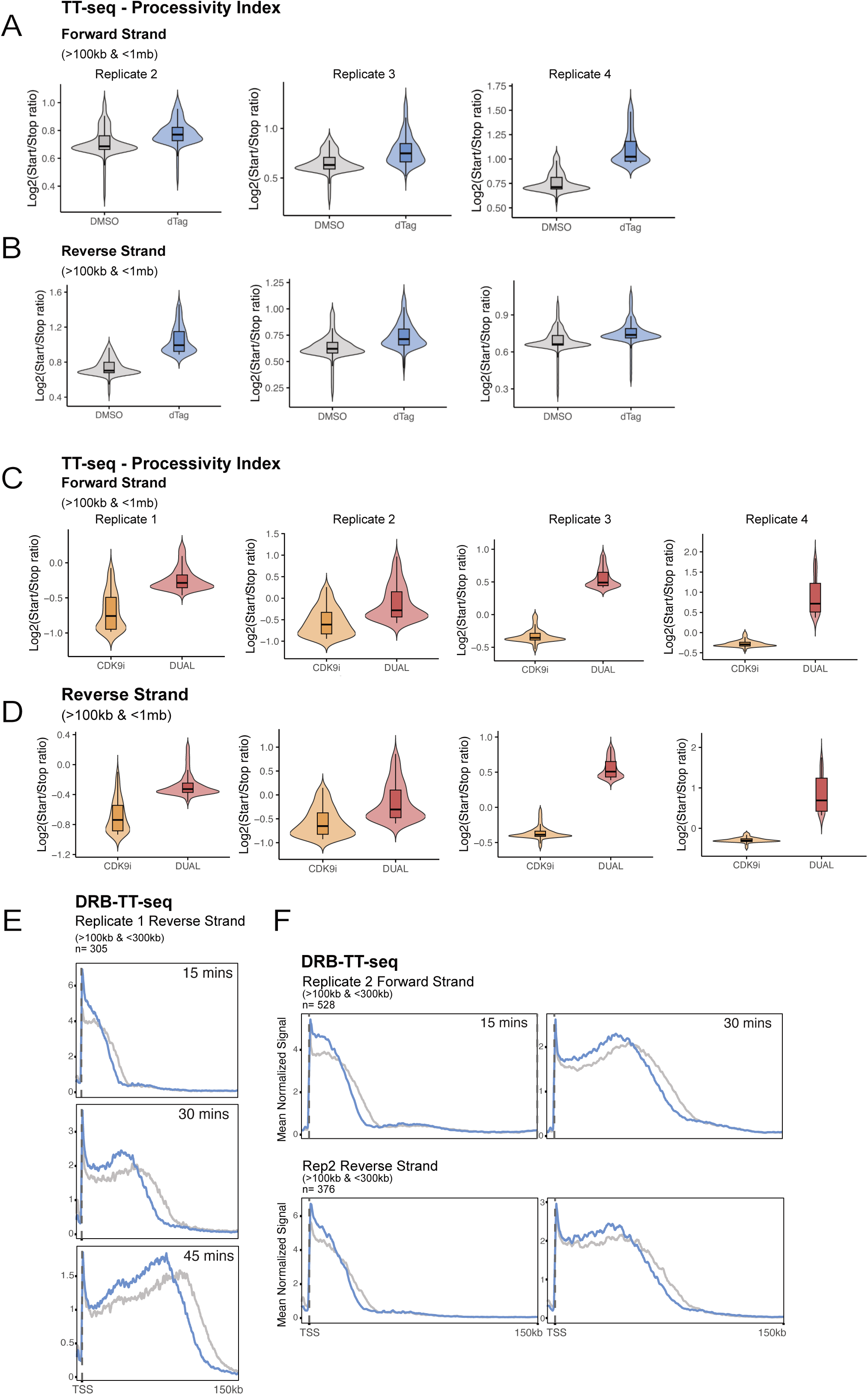
A) Quantified Processivity Index for replicates 2-4 for the forward and B) Reverse strands in DMSO/dTAG^V^-1 TT-seq. C) Quantified Processivity Index for replicates 1-4 for the forward and D) Reverse strands in AZ5576/dual TT-seq. Only genes between 100kb and 1Mb are quantified. Two batches of AZ5576 were used and IC-50 was determined for each batch. Replicates 1 and 2 were treated with 300nM of AZ5576 from batch1, replicates 3 and 4 were treated with 150nM of AZ5576 from batch2. E) Reverse strand of replicate 1 in DMSO/dTAG^V^-1 DRB-TT-seq. F) Forward and reverse strands for replicate 2 in DMSO/dTAG^V^-1 DRB-TT-seq.

**Supplementary data Figure 6:**
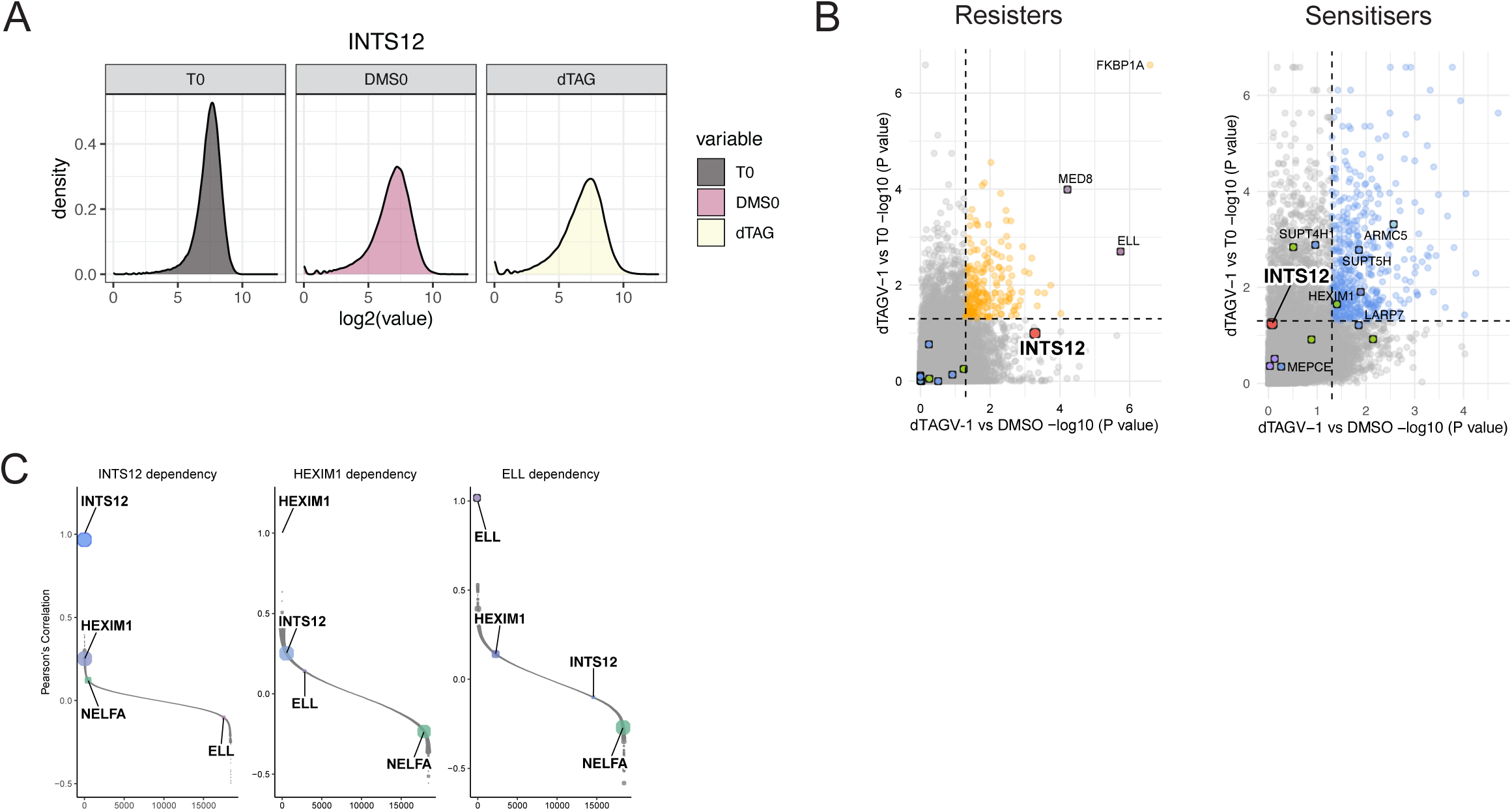
A) Guide distribution at T0 and Tend for DMSO and dTAG^V^-1 for CRISPR-screen showing a decrease of the gini index over time. B) Resisters and Sensitisers scatter plots from the screen showing the degron-tag FKBP12^F36V^ as a hit like INTS12 (positive control for INTS12 degradation) in the dropout screen and LARP7 in the sensitisers. C) Ranked plots visualising DepMap public dependency data 2025Q4 and showing a moderate positive correlation between INTS12, HEXIM1 and NELFA and a negative correlation between INTS12 and ELL.

**Supplementary Data Table 1 -** table with the library minus protacs genes and IDs

## Methods

### Key Resource Table

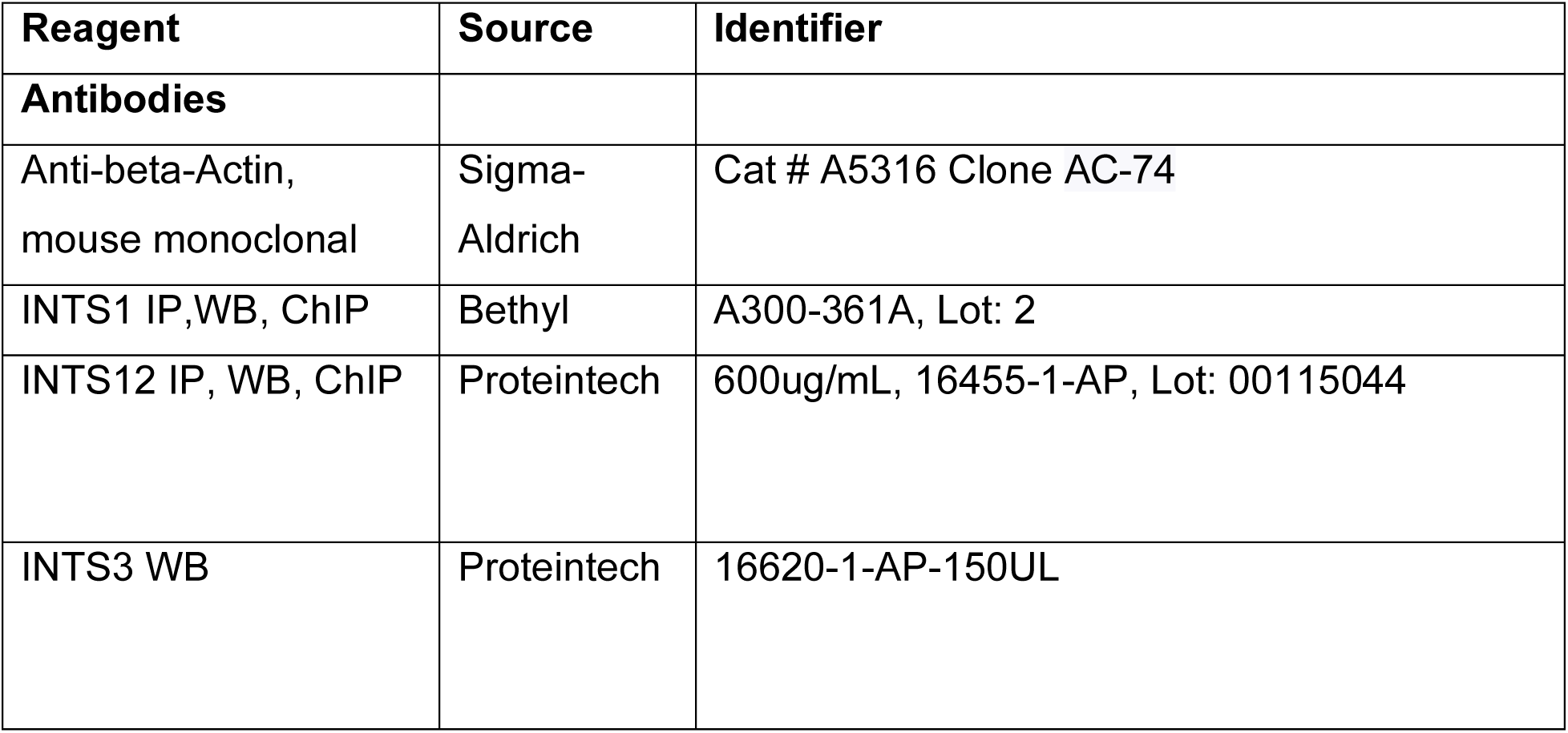
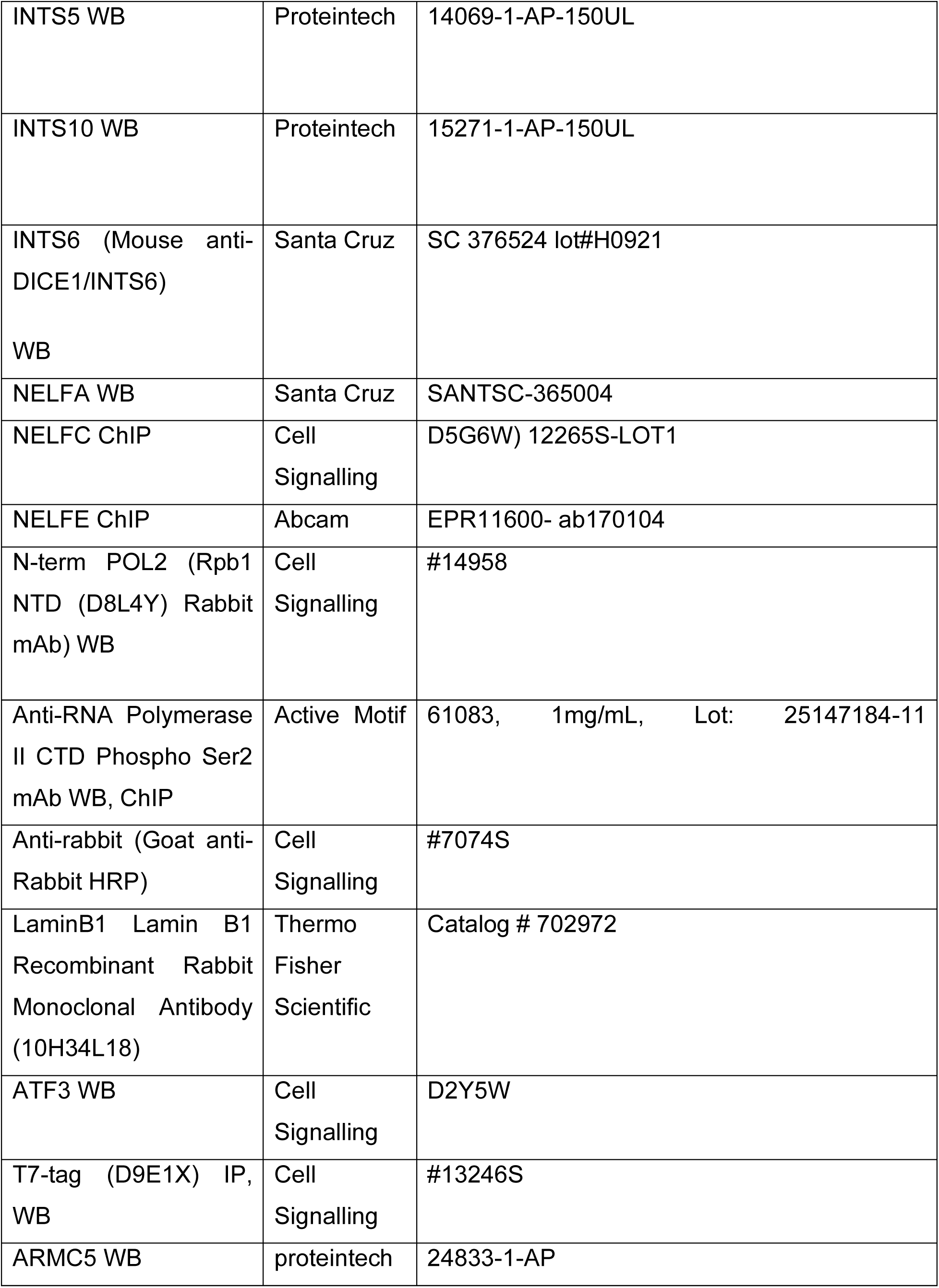
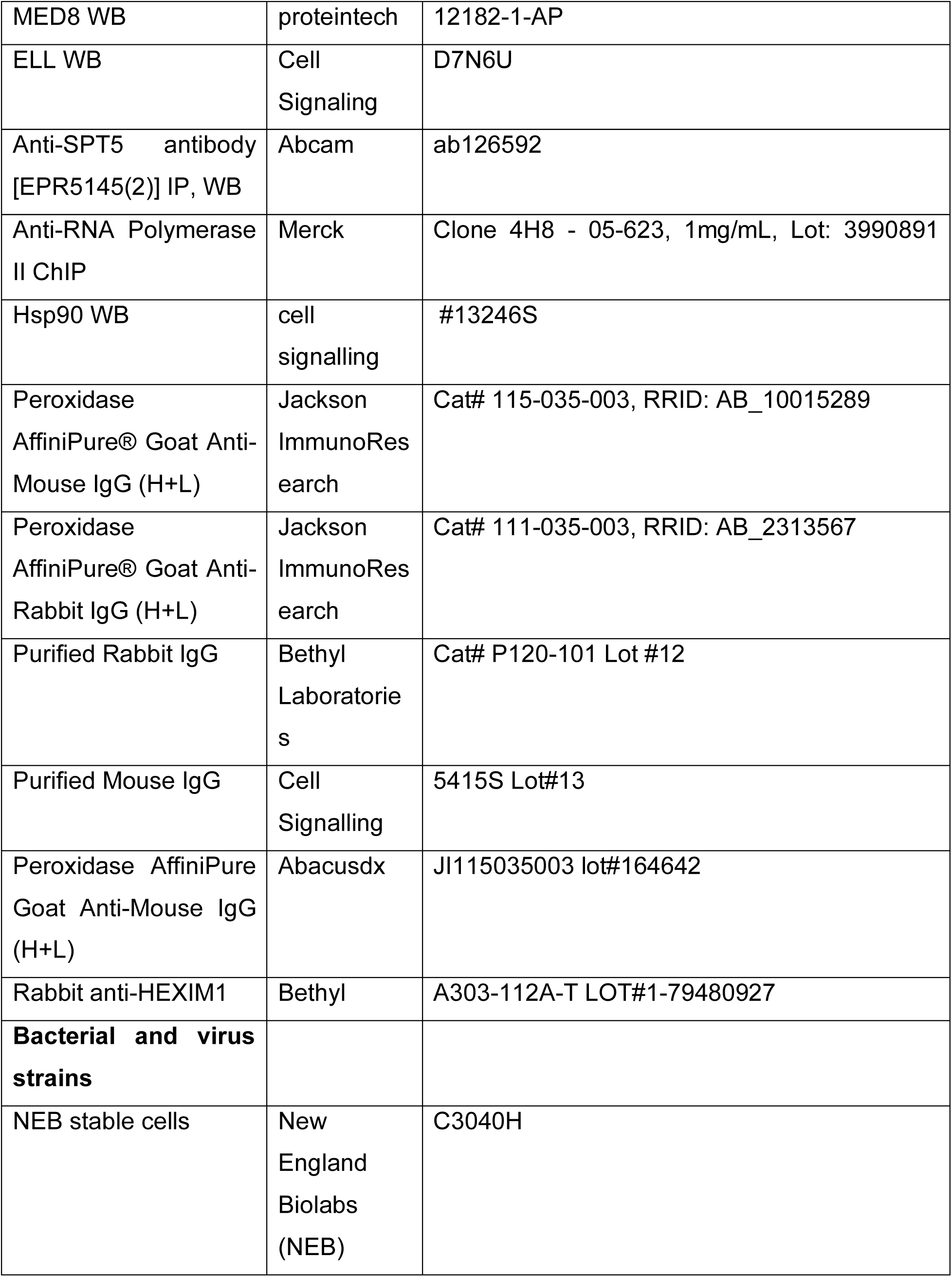
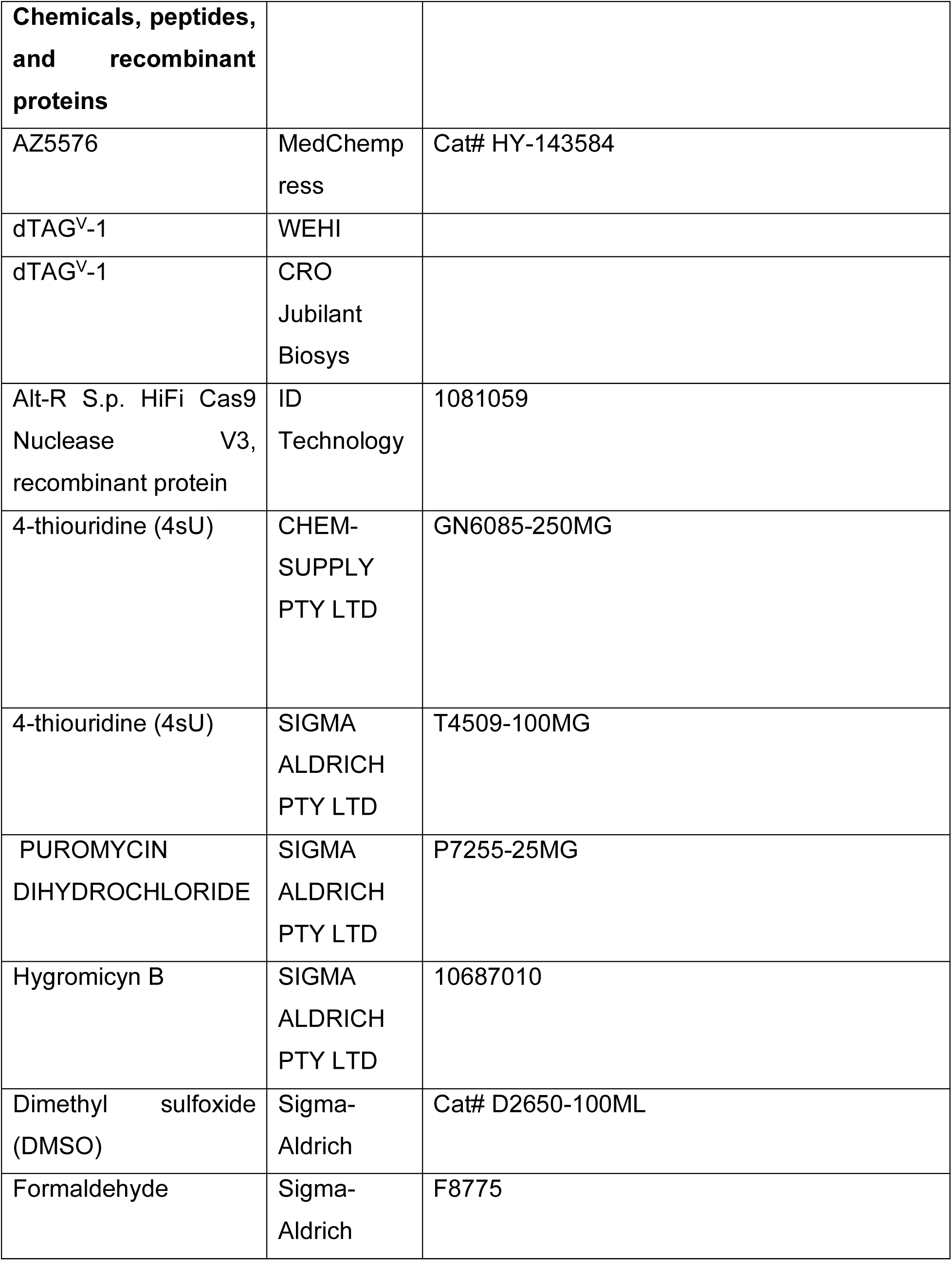
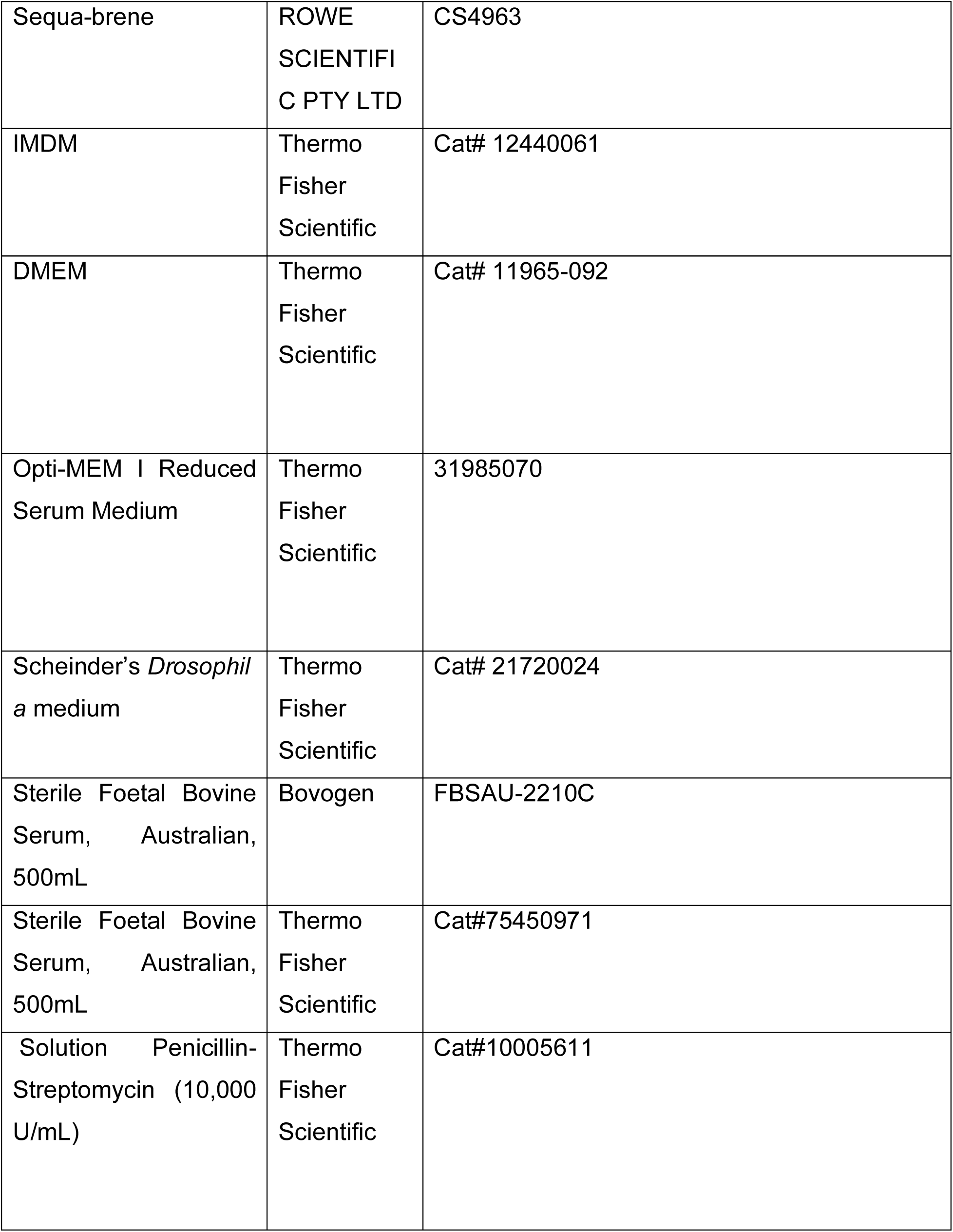
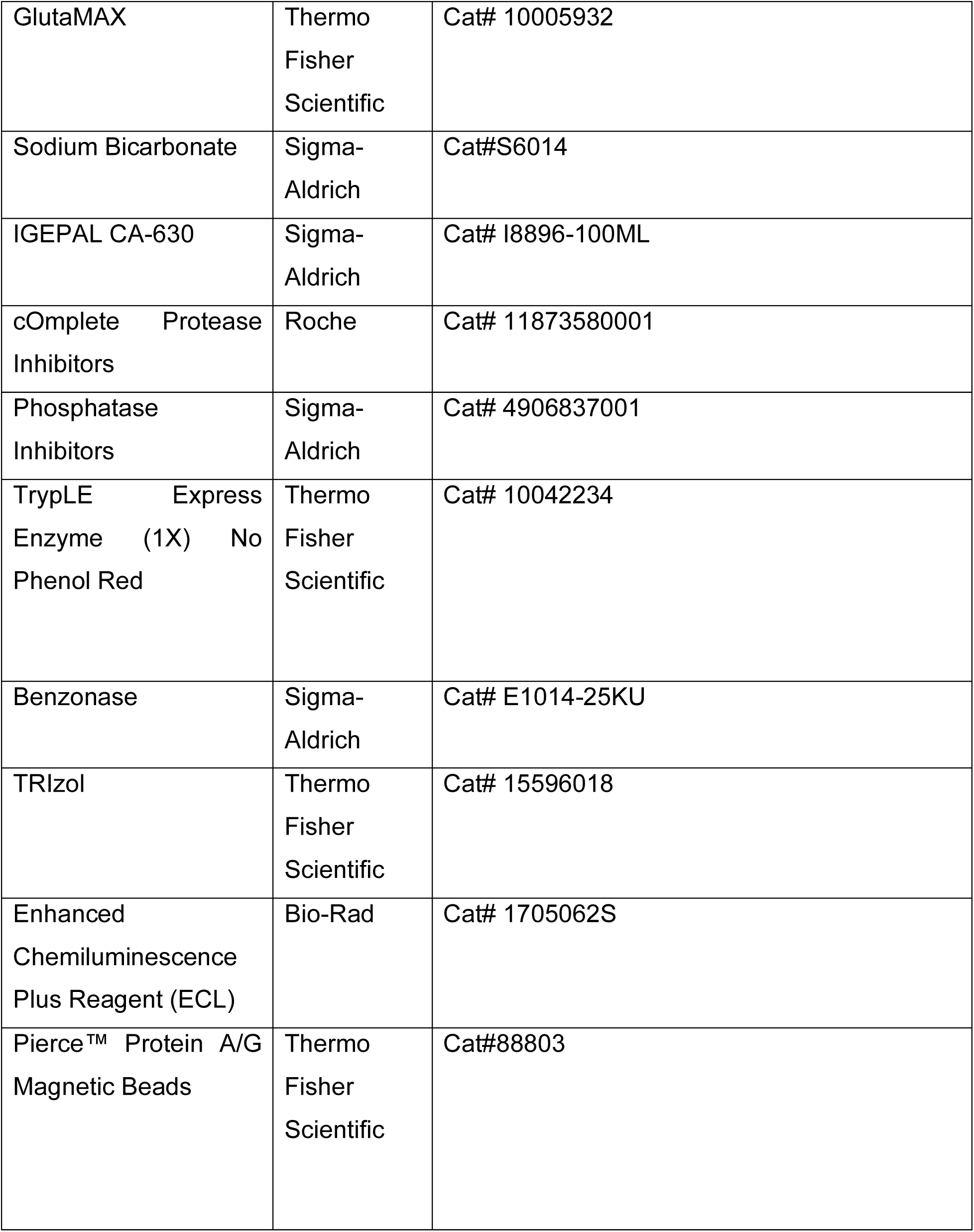
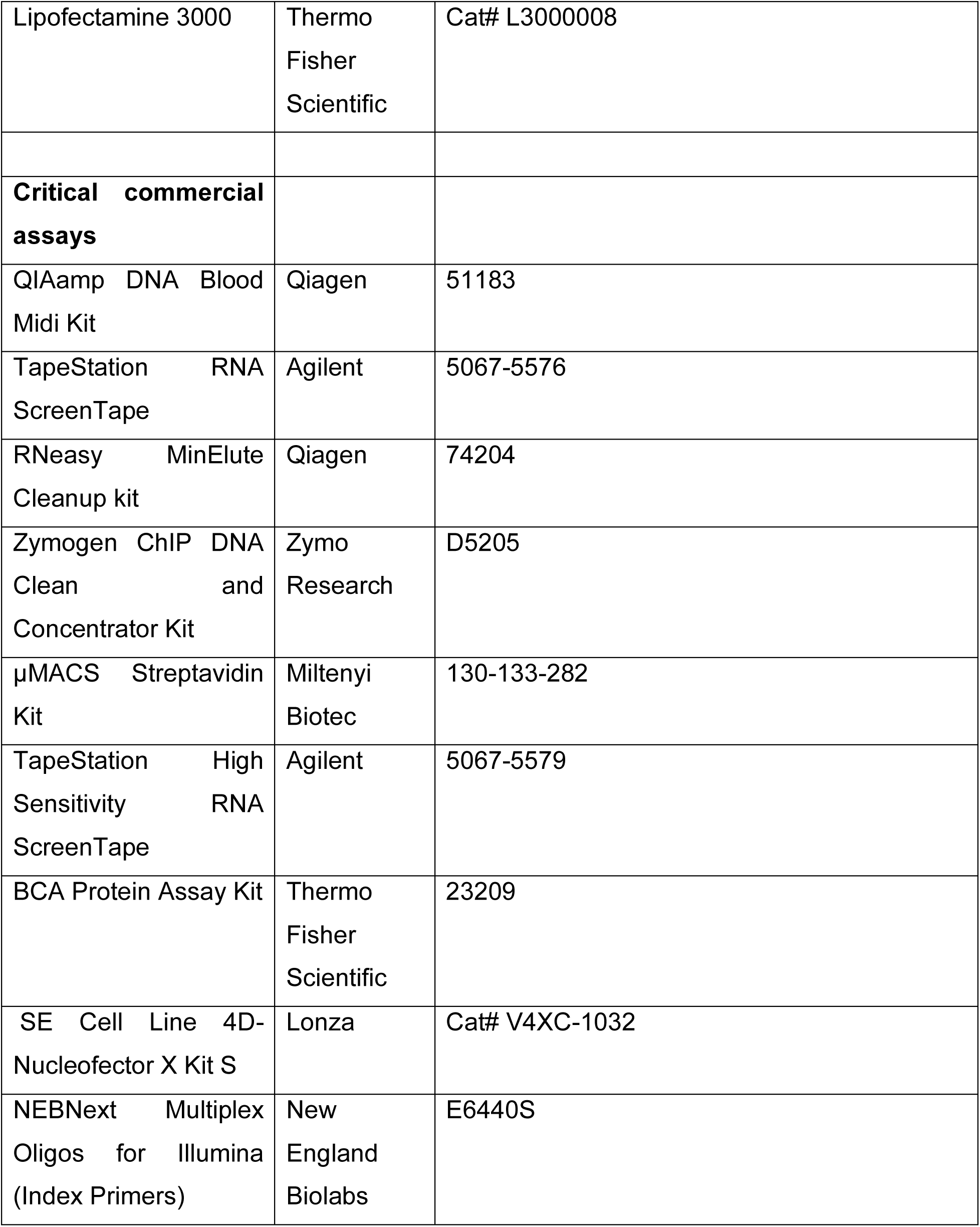
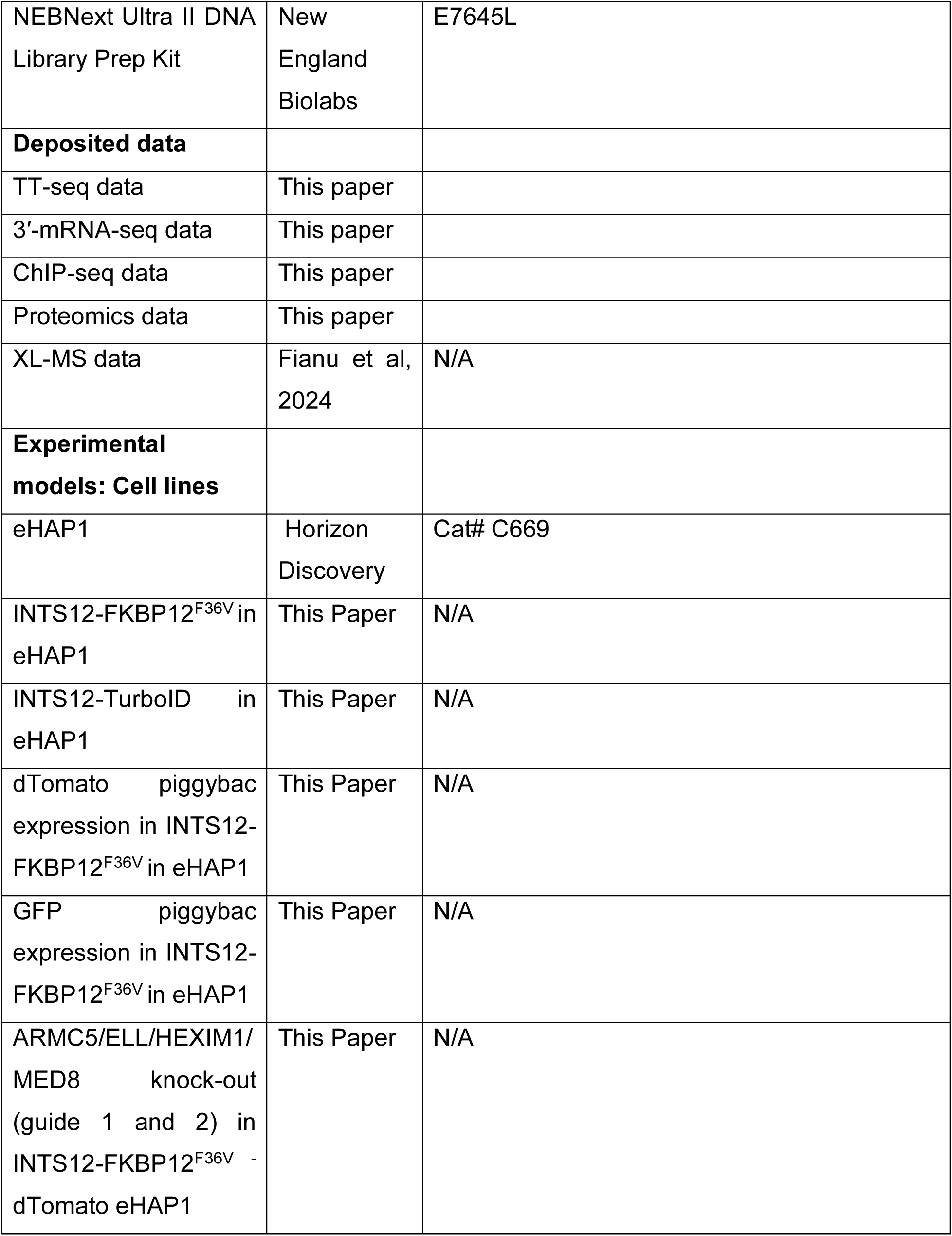
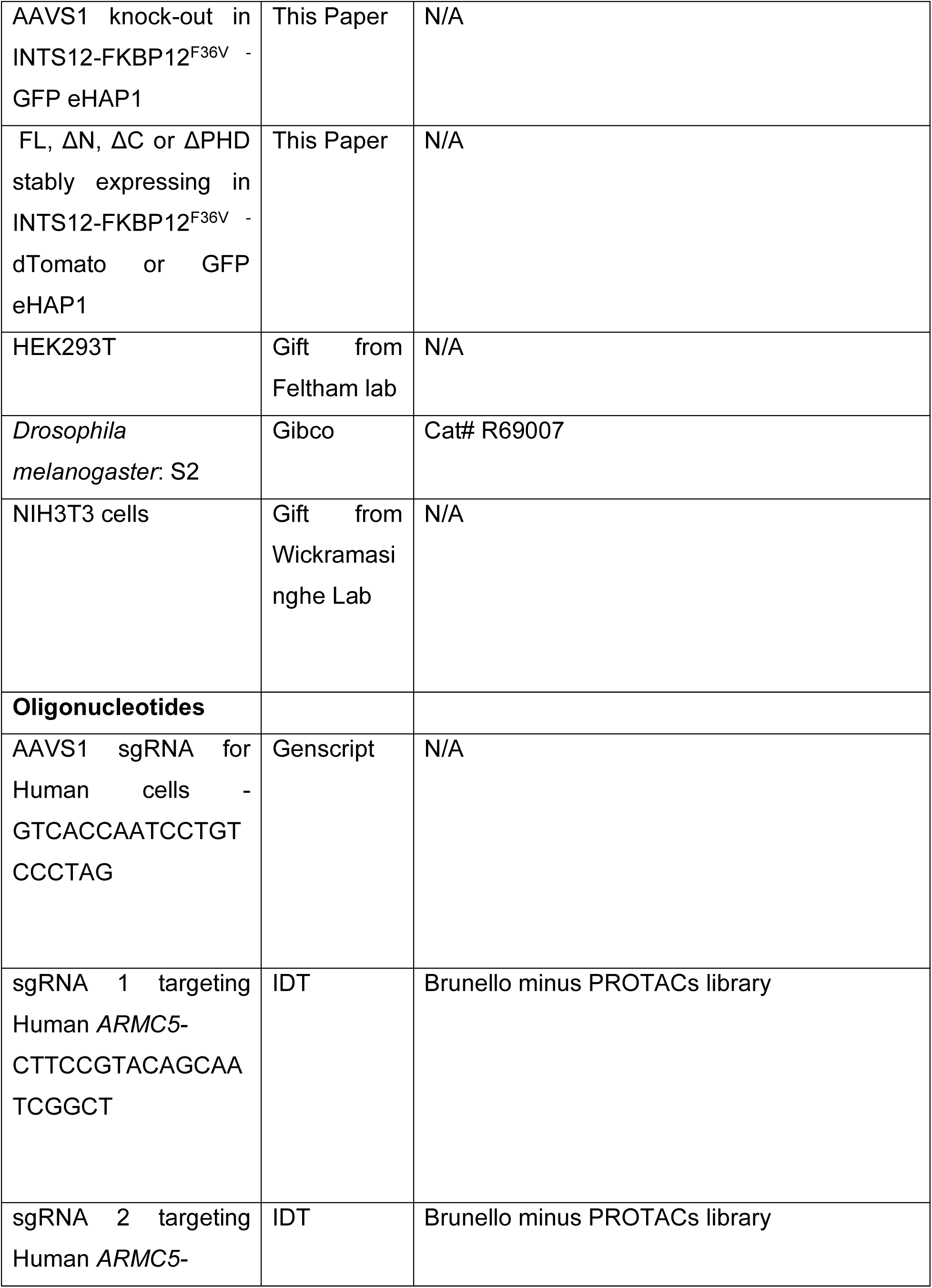
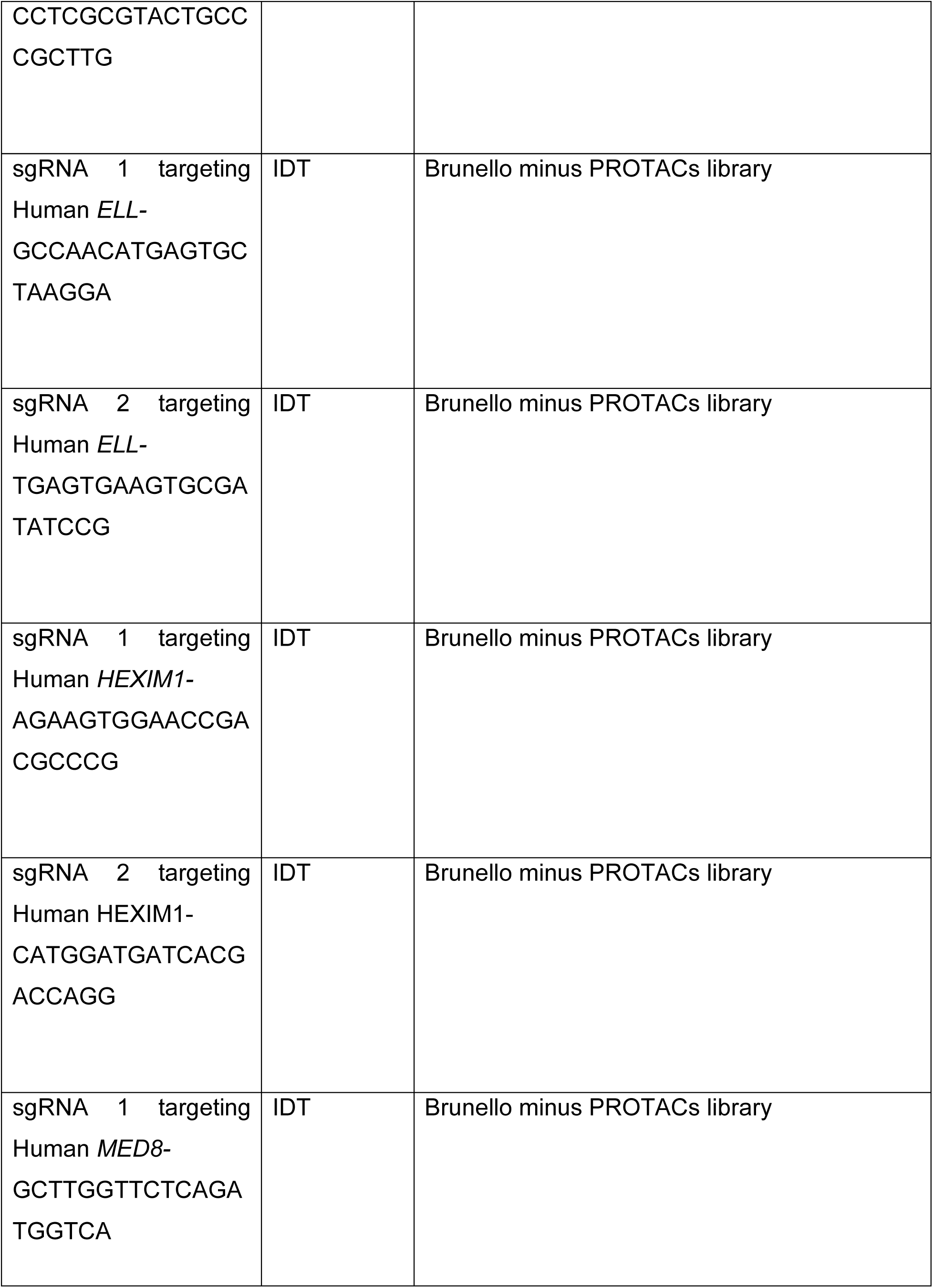
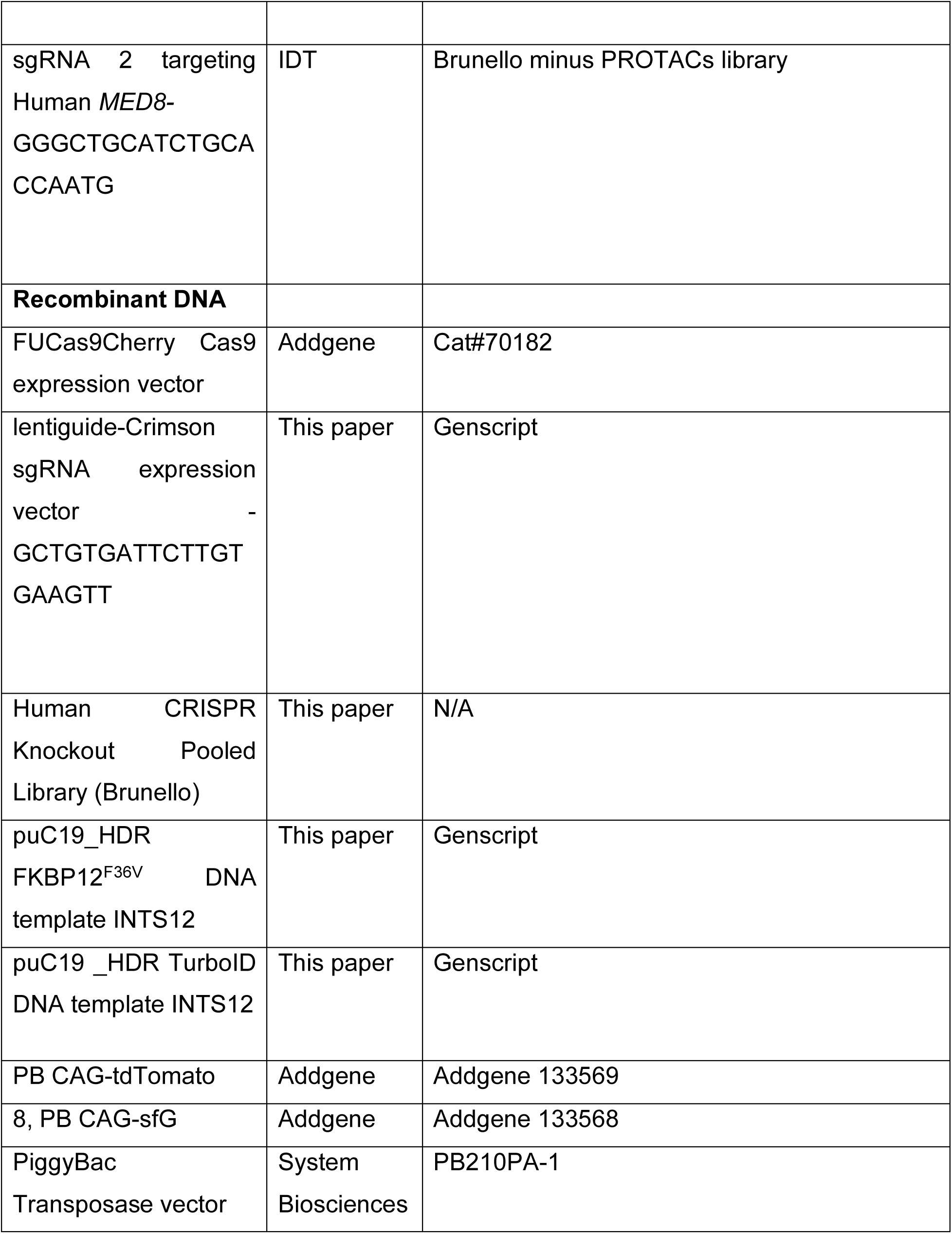
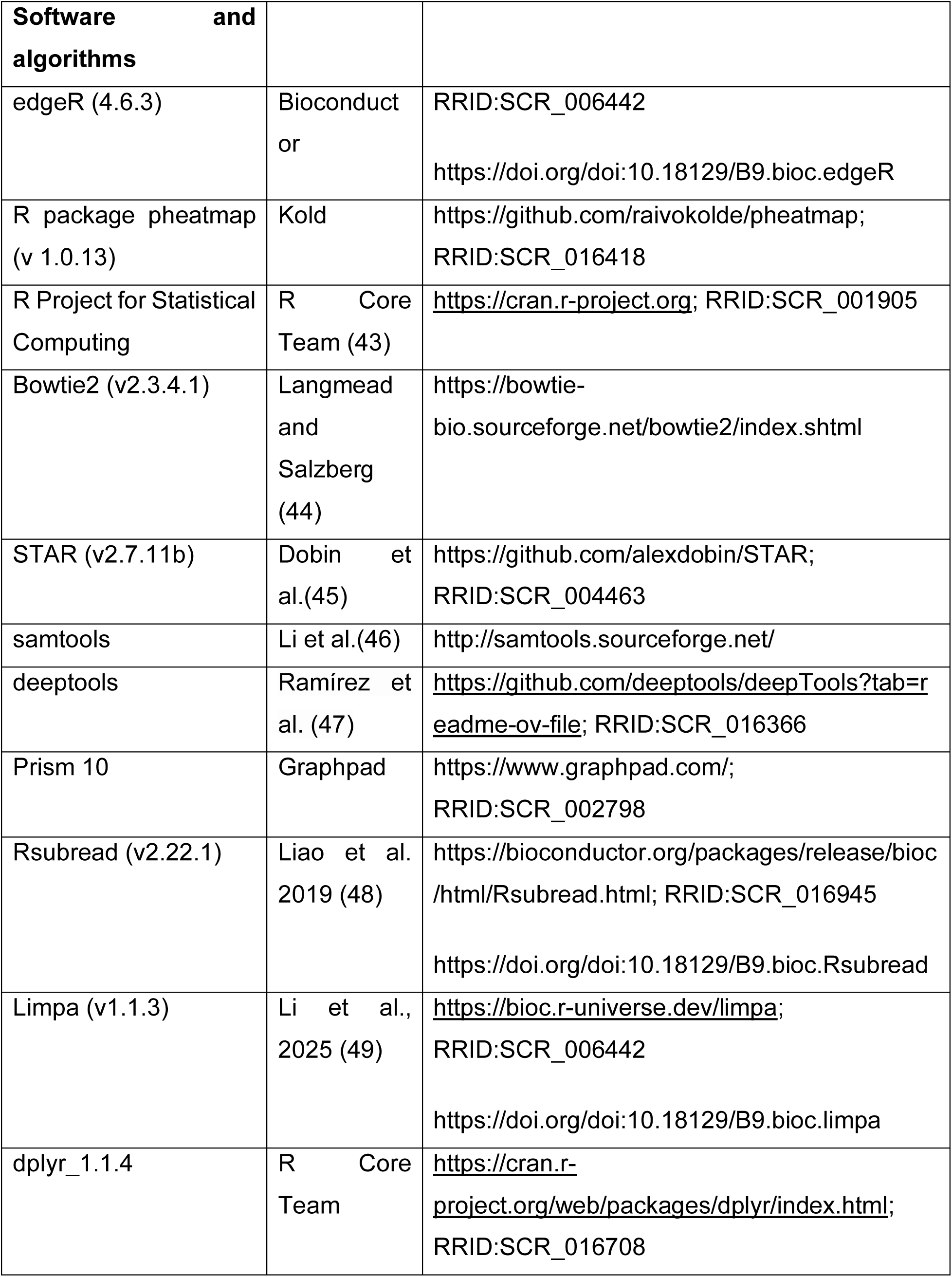
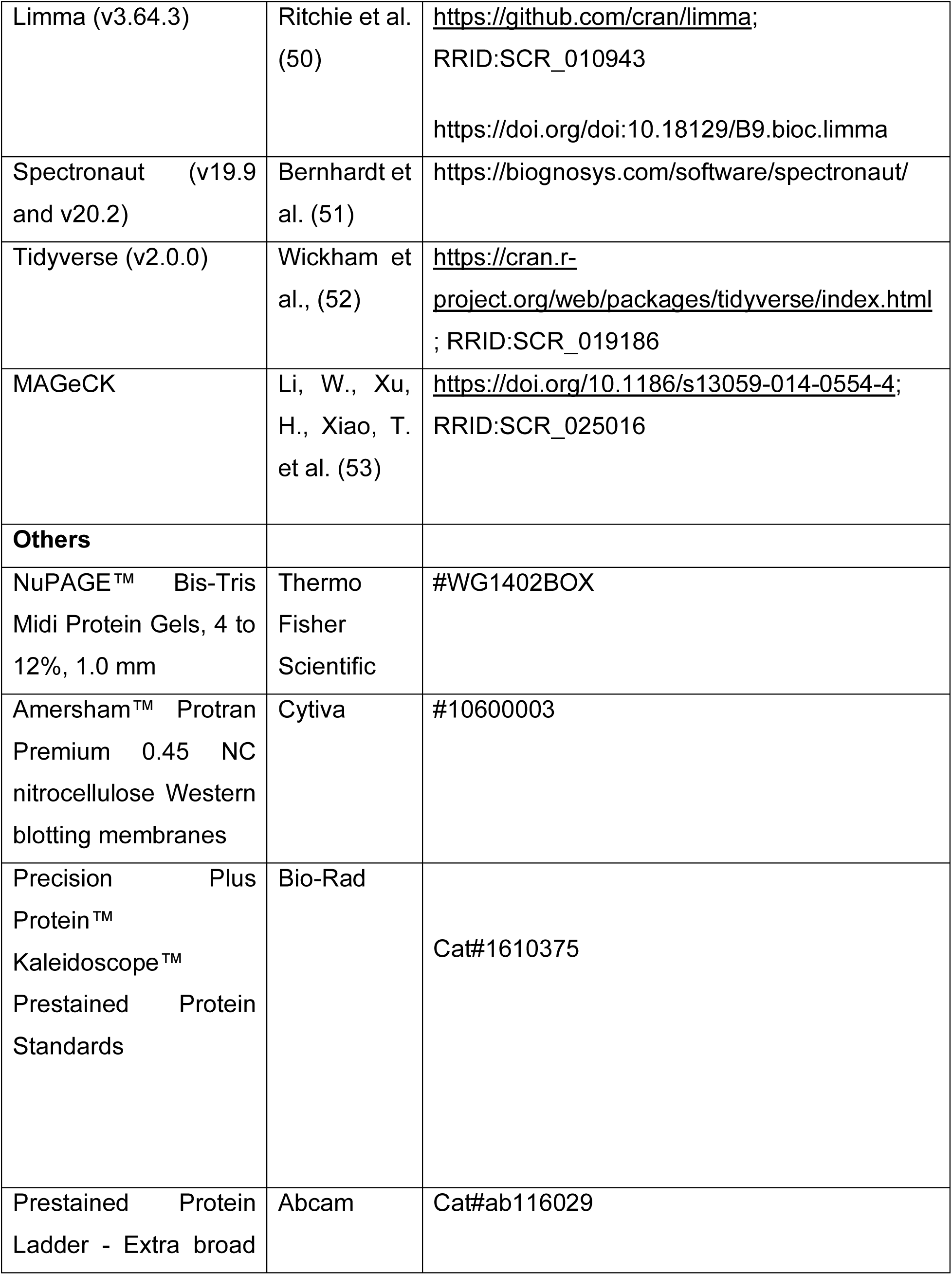
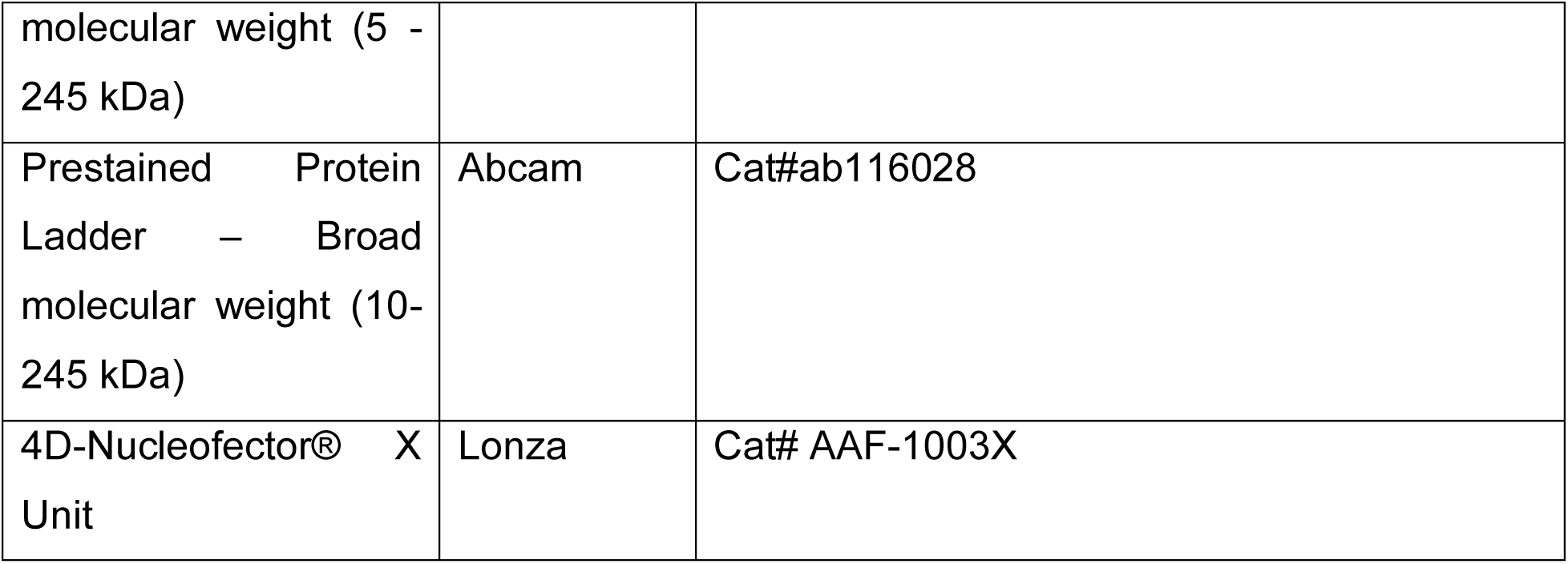

## Resource Availability

### Lead Contact

Further information and requests for resources and reagents should be directed to and will be fulfilled by the lead contact, Stephin J Vervoort.

## Materials Availability

### Data and code availability

Bioinformatics and proteomics datasets generated during this study are available from the NCBI Gene Expression Omnibus (link + GSEA number x) and the Proteomics Identifications Database (PRIDE, link + PXD number x) repositories respectively.

## Experimental Model and Subject Details

### Cell lines and culture conditions

eHAP1 wild-type and gene-edited cells were cultured at 37°C and 5 percent carbon dioxide (CO_2_) in IMDM supplemented with 10% fetal bovine serum (FBS), 100U/mL penicillin, 100 μg/mL streptomycin. Cells were authenticated using short-tandem-repeat (STR) profiling and were also tested for mycoplasma detection (Peter McCallum Cancer Centre service). HEK293T cells were cultured in DMEM medium. Drosophila melanogaster (*D*.*melanogaster*) Shneider 2 (S2) cells were cultured in Shneider’s Drosophila medium (ThermoFisher Scientific 21720) supplemented with 10% HI-FBS, 100U/mL penicillin; 100 μg/mL streptomycin and 2mM GlutaMax™ at room temperature and atmospheric CO_2_.

### Generation of INTS12-FKBP12^F36V^ and TurboID KI line

To generate the knock-in lines, the sgRNA and template plasmids were cloned and synthesised by Genscript. Cells 5x10^4 were seeded in a 6-well plate and transfected with Lipofectamine 3000 (7.5uL of lipofectamine and 5uL of P3000 per condition) the following day. Cas9:guide plasmids were added in a 2:1 ratio to a total amount of 2ug and 2ug of donor plasmid. Media was changed 16 hrs later. 46 hours later cells were selected with 2ug/mL of puromycin. Cells that were both, haploid and GFP positive were sorted and grown. For haploid sorting, cells were stained with Hoechst 33342 staining solution to differentiate between haploids and diploids. Cell lines were validated via immunoblotting. Cells were left in culture for few passages to allow them to become diploids and then sorted again for pure diploids and GFP positive cells.

### Generation of INTS12-FKBP12^F36V^ expressing dTomato piggybac vector

A total of 1x10^6^ INTS12-FKBP12^F36V^ cells were plated in 6-well plate. 24 hours later cells were transfected with the 0.2 ug PiggyBac Transposase vector (PB210PA-1), piggybac dTomato (Addgene #133569) plasmids and the transposase in 5:1 ratio for a 2.5ug total of plasmid delivered with lipofectamine 3000 following manufacture protocol with OptiMEM. After three days from transfection, cells were sorted for dTomato positivity.

### Generation of INTS12 mutants stable cells

To produce mutants stably expressing lines, lentiviral particles were produces in HEK293T cells with the INTS12 mutants. Plasmids expressing the mutants were ordered from Genscript. INTS12-FKBP12^F36V^ cells were transduced with the mutants viruses and sequabrene and then selected with blasticidin 10ug/mL for 10 days.

### Generation of screen validation cells KOs

400.000 cells INTS12-FKBP12^F36V^ cells/condition were nucleofected with the SE buffer Lonza kit following manufacture protocol. 180 pmol of 200uM sgRNA and 60pmol of recombinant Cas9 (IDT) were mixed together and incubated for 20 min at RT and then added to the cells and then nucleofected with 4D-Nucleofector® X Unit (Lonza)with the program DZ-113, Guides sequences are in the Key Resource Table.

## Method Details

### Design and synthesis of the PROTAC-compatible CRISPR library

The list of sgRNA spacer sequences in the Brunello Human CRISPR knockout pooled library (Addgene #73179) was used as a starting point. From this list, sgRNAs targeting known or potential mediators of dTAG/PROTAC-mediated degradation (63 genes in total) were manually removed. This resulted in a list of 77,189 spacer sequences. Briefly, the corresponding oligonucleotide pool was synthesised (Genscript), PCR amplified and cloned into the LentiGuide-Hygro vector (Addgene #139462) using the BsmBI (NEB) restriction sites. For a complete list of sgRNA sequences see Supplementary Table 1.

### Genome-wide CRISPR-Cas9 Screens

INTS12-FKBP12^F36V^ cells were engineered to stably express humanized *S.pyogenes* Cas9 endonuclease by lentiviral transduction with the FUCas9Cherry vector (Addgene 70182) and subsequent FACS-selection for mCherry-positive cells (eHAP1-Cas9).

For eHAP1-Cas9 survival screens, cells were transduced with the PROTAC-compatible genome-wide CRISPR library (77,189 sgRNAs) cat a MOI of 0.3 and a fold representation of 250 for each individual sgRNA.

Transduced cells were selected with hygromycin (300 μg/mL) for 7 days, at which time cells were split into relevant treatment conditions (DMSO, dTAG^V^-1 250nM) and cell pellets were collected and snap-frozen for T_0_ reference controls. Cells were kept in culture fro 20 days and samples were collected for final time point Genomic DNA was extracted using the QIAamp DNA Blood Midi Kit (Qiagen Cat.# 51185). PCRs were performed using the Ex Taq DNA polymerase (Takara Cat.# RR001C) following the protocol for PCR of sgRNAs from genomic DNA for Illumina sequencing from the BROAD Institute. Libraries were sequenced SE 100 at WEHI’s Genomics Facility.

### RNA Sequencing and Data Analysis

For all RNA-seq experiments, 5 x 10^5^ eHAP1s 12KI cells were seeded per well in a 6 well plate in 2 ml of complete growth medium one day prior to treatment. Cells were treated with 250 nM dTAG^V^-1 for 2h and/or 300 nM CDK9i for 1h (AZ5576 Batch 1). Following treatment, total RNA was isolated using the RNeasy MinElute Cleanup Kit (Qiagen) with on-column DNase digestion (according to manufacturer’s instructions). Isolated RNA samples were sent to Novogene for library preparation and sequencing. Strand-specific RNA-seq libraries were prepared using the NEBNext Ultra II directional RNA library prep kit for Illumina. Sequencing was performed on an Illumina NovaSeq X Plus platform to generate 150 bp paired-end reads.

Paired-end reads were aligned to the human reference genome (GRCh38) using Rsubread (v2.22.1). Gene-level counts were generated using the featureCounts function with GENCODE v45 annotations. Strand-specific counting was specified (strand-specific = 2) and reads overlapping exons were summarised at the gene level using the gene_name attribute. Differential gene expression analysis was performed using limma (v3.64.3).

### Competitive Proliferation and CTV Assays

For competition assays, WT and INTS12-KI, INTS12-KI mutants etc were plated 1:1 for a total of 40.000 cells in 12-well plates, 3 replicates/each condition and treated with 250 nM of dTAG^V^-1 and 100 nM of AZ5576. Every three days, cells were analysed by Flow Cytometry to check % of dTomato and GFP +ve/-ve cells. For CTV assays 5x10^6 WT and INTS12-KI cells were stained with CellTrace™ Violet dye following manufacturer’s instructions. Cells were sorted using the 405/450 nm channel. 4 x10^4 cells were seeded in 12-well plates, 3 technical replicates per condition. Cells were treated with 250 nM of dTAG^V^-1. Cells were analysed at day 3, 5 and 7 by Flow Cytometry.

### SDS page and Immunoblotting

For whole cell extracts, pellets were resuspended in 1X Laemmli Buffer (0.625M Tris-HCl, 0.07M (2%) SDS, 10% glycerol) and boiled at 98 °C for 10 minutes. Protein quantification has been done either with BCA or Qubit assay (Q33212, Thermo Fisher Scientific) and normalised with Laemmli buffer. 20 ug of protein got loaded on the gel and proteins were separated on 4-15%, 12% or 7.5 % gels (biorad) and transferred to a PVDF membrane with dry transfer (biorad) with specific program depending on the size of the protein. Membranes were then blocked with skimmed milk 5% in TBS-T buffer for 1h at RT and then incubated with the relative primary antibody (1:1000 dilution) diluted in 5% skimmed milk-TBSt () at 4°C overnight (O/N). The following day, the membrane was washed for 5 minutes 3 times with TBST and then incubated with HRP-conjugated secondary antibody diluted in 5% skimmed milk (1:10.000 dilution) (primary and secondary antibodies are listed in the Key Resource Table. Result was visualised by adding ECL components 1:1 and using the Chemidoc (BioRad).

### Glycerol Gradient Ultracentrifugation

For glycerol gradient analysis, INTS12-FKBP eHAP1 cell lines were plated in 15 cm dishes in IMDM supplemented with 10% FBS and 1% glutamax and maintained at 37°C until they were 70-90% confluent. Prior to collection, cells were treated with either 250 nM dTAG^V^-1 or the corresponding volume DMSO for two hours. Cells were then washed 3X with PBS, and resuspended in 2X cell pellet volume of RIPA lysis buffer (Millipore 20-188) supplemented with SDS (0.1%), protease inhibitor (1X), PhosSTOP (1X), and benzonase (1 µL/mL). Lysates were agitated either by end-to-end rotation or shaking (1500 rpm) for 1 hour at 4°C. Cell lysates were further mechanically disrupted by passing lysates through a 31G needle 3 times, and cell debris was spun down at 20,000 g for 10 minutes 4°C and discarded. Lysates were quantified with a BCA Assay and normalized to protein quantification prior to being loaded onto the glycerol gradients.

The 10–50% glycerol gradients were prepared by adding 2.5 mL of buffer A (50 mM HEPES pH 7.9, 200 mM KCl, 30 mM MgCl_2_, 0.2 mM EDTA, 1 mM TCEP, 1x cOmplete EDTA-free protease inhibitor (Roche), and 50% glycerol) to a 5 mL polypropylene tube (#326819, Beckman Coulter), followed by gently pipetting 2.5 mL of buffer B (50 mM HEPES pH 7.9, 200 mM KCl, 30 mM MgCl_2_, 0.2 mM EDTA, 1 mM TCEP, 1x cOmplete EDTA-free protease inhibitor (Roche), and 10% glycerol) to the top of the gradient on ice. The tube was gently tilted horizontally in a minute time to allow the glycerol layers to form a continuous gradient. The tube was kept in this position for 4 hours at 4 °C before returning to a vertical position for an hour to stabilise the gradient. A 200 µL of the FLAG-IP elution was layered on the top of the gradient. The gradient was then placed into a MLS-50 swinging-bucket rotor (Beckman Coulter) and ultracentrifuged at 48,000 rpm for 16 hours at 4 °C. Following centrifugation, 500 µL fractions were collected manually starting from the top of the gradient. The fractions were heated at 80 °C for 5 min with 1X Laemmli sample buffer (Bio-rad) and examined by Western Blotting.

### TurboID Labelling

Wild-type and INTS12-TurboID eHAP1 cell lines were plated in sets of four replicates in 15 cm dishes, in biotin-free IMDM that was supplemented with FBS and glutamax, such that they were 70-90% confluent upon biotin treatment. Biotin (100 mM) was added to both conditions and cells were maintained at 37°C for one hour. For the RNAPII-TurboID experiment, wild-type and RNAPII-TurboID INTS12-FKBP lines were plated as above. Cells were kept at 37°C and treated with 250 nM dTAG^V^-1 for one hour followed by 100 mM biotin for an additional hour. All cells were then immediately washed with ice-cold PBS five times and collected through manual scraping prior to lysis.

### Multi-Lysis Fractionation

Cell lysates were generated through a multi-lysis fractionation performed similar as previously described (18). All buffers were supplemented with PhosSTOP (Merck) and cOmplete™, EDTA-free Protease Inhibitor Cocktail (Merck) tablets. In brief, cell pellets were resuspended in excess of 10X cell pellet volume of hypotonic wash buffer (10 mM HEPES pH 7.5, 10 mM potassium chloride, 1.5 mM magnesium chloride (MgCl_2_), and incubated on ice for 25 minutes. Nuclei were pelleted at 1000 g for 15 minutes at 4°C, supernatant discarded, and washed in hypotonic buffer followed by immediately pelleting again at 1000 g for 15 minutes at 4 degrees. Nuclei were then resuspended in 2X original cell pellet volume of nucleoplasmic lysis buffer (20 mM HEPES pH 7.9, 1.5 mM MgCl_2_, 150 mM potassium acetate, 10% (v/v) glycerol, 0.05% (v/v) NP-40) and incubated on ice for 20 minutes. Chromatin was pelleted at 20,000 g for 20 minutes at 4°C and supernatant was collected as the nuclear fraction and stored on ice. Chromatin was resuspended in 2X original cell pellet volume of chromatin digestion buffer (20 mM HEPES pH 7.9, 1.5 mM MgCl_2_, 10% (v/v) glycerol, 0.05% (v/v) NP-40, 150 mM sodium chloride (NaCl) and benzonase (1:1000 enzyme:buffer)) and incubated on ice for 60 minutes. The low salt fraction was then centrifuged at 20,000 g for 20 minutes at 4°C and supernatants were pooled into their respective nuclear fraction. Pellets were resuspended with high salt chromatin digestion buffer (20 mM HEPES pH 7.9, 3 mM ethylenediaminetetraacetic acid (EDTA), 1.5 mM MgCl_2_, 10% (v/v) glycerol, 500 mM NaCl, and 0.1% (v/v) NP-40) at 2X original cell pellet volume and incubated on ice for 20 minutes. Six original cell pellet volumes of high salt dilution buffer (20 mM HEPES pH 7.9, 3 mM EDTA, 1.5 mM MgCl_2_, 10% (v/v) glycerol, 0.1% (v/v) NP-40) and cells were immediately centrifuged at 20,000 g for 20 minutes at 4°C and combined with previous nuclear fractions. The combined nuclear fraction lysate was snap frozen in liquid nitrogen and stored at -80°C until biotin enrichment.

### Protein Purification, Clean Up, and Digestion

Lysates were quantified using a BCA assay and protein quantities were normalized. Biotinylated proteins were enriched by incubating quantified lysates with high-capacity streptavidin agarose beads (Pierce) overnight at 4°C with end-to-end rotation. After incubation, beads were washed three times with 0.5% (w/v) SDS in PBS. Proteins were reduced with 100 mM DTT in 0.5% (w/v) SDS in PBS and then washed twice in UC Buffer (6M urea, 100 mM Tris-HCl pH 8.5). Beads were then captured with the Pierce Snap Cap Spin Columns (69725) and alkylated for 20 minutes with 50 mM iodoacetamide in UC buffer in the dark. Beads were then washed extensively with 1-minute spins at 1,000 g as follows: six times UC buffer, four times PBS, and then three times MS-grade milliQ water. Beads were resuspended in 50 mM ammonium bicarbonate (NH_4_HCO_3_) containing 2 ug of Trypsin and incubated overnight at 37°C. Digested peptides were collected by centrifugation at 1,000 g for 1-minute, and beads were washed once with 50 mM ammonium bicarbonate and elutants pooled. Peptide elutions were dried to completeness at 45°C on the SpeedVac, and resolubilised in mass spectrometry loading buffer (2% (v/v) ACN and 0.1% (v/v) formic acid) prior to injection.

### Endogenous IP

For the INTS12 immunoprecipitation, wild type eHAP1 cell lines were plated in sets of four replicates in 15 cm dishes in IMDM supplemented with 10% FBS and 1% glutamax and maintained at 37°C until they were 70-90% confluent. For the INTS1 immunoprecipitation, INTS12-FKBP eHAP1 cells were plated the same as the wild type cells, and treated with either 250 nM dTAG^V^-1 or the corresponding volume DMSO two hours prior to collection. Cells were washed with 2X in PBS and subsequently lysed with the multistep fractionation as described above. After lysis, lysates were quantified using a BCA assay and protein quantities were normalised to approximately 500 ug of protein. Two micrograms of either Rabbit IgG (Bethyl Laboratories), INTS12 (Proteintech 16455-1-AP), or INTS1 (A300-361A Lot#2) antibody were added to their respective sample and incubated with end-to-end rotation at 4°C overnight. Following antibody/IgG incubation, Protein A/G beads (Pierce) were washed 2X with PBS and resuspended in IP Wash Buffer (150 mM NaCl, 20 mM Tris-HCl pH 7.5, 1.5 mM MgCl_2_, 3 mM EDTA, 10% Glycerol, 0.1% NP-40). Beads were added to samples and incubated at 4°C with end-to-end rotation for four hours. Beads were then washed 5X with IP Wash Buffer and eluted in 50 uL of Elution Buffer (100 mM Tris-HCl pH 7.5, 1% SDS, 0.5 mM EDTA) at 65°C for three minutes. Two more elutions of 30 uL were performed the same way and elutants were pooled and snap frozen prior to Filter Assisted Sample Preparation (FASP) cleanup. For Western blot samples, SDS loading dye was added to beads and boiled at 95°C for 10 minutes for elution.

### SLF-Biotin Pulldown

Wild type or INTS12-FKBP eHAP1 cells were plated and lysed as described for the endogenous immunoprecipitations. Dynabeads™ Streptavidin Magnetic Beads (Invitrogen) were washed 2X in the nuclear lysis buffers (1:1:1:3 of Nucleoplasmic Lysis Buffer:Chromatin Digest Buffer:High Salt Digestion Buffer:High Salt Dilution Buffer). Beads were either pre-incubated with 10 mM SLF-biotin or DMSO for three hours at 4°C with end-to-end rotation. Beads were then washed with IP Wash Buffer three times, added to cell lysates, and incubated overnight at 4°C with end-to-end rotation. Beads were subsequently washed 5X with IP Wash Buffer and eluted as described above.

### FASP (Filter Assisted Sample Preparation)

Protein digestion was carried out using a modified FASP protocol [4]. All centrifugation steps were performed at 14,000 × g for 10–15 min at room temperature using Vivacon 500 filter units with a 30 kDa molecular weight cut-off (Sartorius, VN01H22). Protein elutions were loaded onto FASP columns and were solubilised in Urea/Tris buffer (stock: 12 M Urea in 100 mM Tris-HCl pH 7.6) and reduced with tris(2-carboxyethyl)phosphine (TCEP, final concentration 10 mM) for 30 min at room temperature. Columns were centrifuged and washed with a Urea/Tris buffer. Proteins were alkylated using 100 ul of 50 mM iodoacetamide (IAA) prepared in Urea/Tris buffer, followed by incubation in the dark for 20 minutes at room temperature. Columns were washed three times with Urea/Tris buffer and subsequently three times with 50 mM ammonium bicarbonate (AmBic). Flow-through was discarded after each wash. Trypsin digestion was performed by adding 1 µg Trypsin Gold (Promega) and 0.5 ug Lys-C (Fujifilm Wako, Cat# 129-02541) in 50 mM AmBic to each column and incubated overnight at 37°C. Following digestion, columns were transferred to fresh collection tubes and centrifuged. An additional 50 µL of 50 mM AmBic was added to recover remaining peptides, followed by a final centrifugation. Peptides were acidified in 1% formic acid and lyophilized to dryness using a CentriVap (Labconco) before being reconstituted in 30 µL (200 µL for NELF and SPT5 IPs) of 0.1% Formic Acid (FA)/2% acetonitrile (ACN) prior to mass spectrometry analysis.

### Mass Spectrometry

Endogenous INTS12 IP and INTS12-SLF biotin were acquired on timsTOF using DIA mode. Reconstituted peptides were separated by reverse-phase liquid chromatography on a 15 cm C18 fused silica column with an integrated emitter tip (IonOpticks, ID 75 µm, OD 360 µm, 1.6 µm C18 beads) using a custom nano-flow HPLC system (Thermo Ultimate 300 RSLC Nano-LC, PAL systems CTC autosampler). The HPLC was coupled to a timsTOF Pro (Bruker) equipped with a CaptiveSpray source. Peptides were loaded directly onto the column at a flow rate of 600 nL/min with buffer A (99.9% Milli-Q water, 0.1% FA) and eluted at 400nL/min on a 30-min linear analytical gradient of increasing buffer B (90% ACN, 0.1% FA) from 2 to 34%. The timsTOF Pro (Bruker) was operated in diaPASEF mode using Compass Hystar 5.1. The settings on the TIMS analyzer were as follows: Lock Duty Cycle to 100% with equal accumulation and ramp times of 100 ms, and 1/K0 Start 0.6 V.·/cm2 End 1.6 V·s/cm2, Capillary Voltage 1400V, Dry Gas 3 l/min, Dry Temp 180°C. The DIA methods were set up using the instrument firmware (timsTOF control 2.0.18.0) for data-independent isolation of multiple precursor windows within a single TIMS scan. The method included two windows in each diaPASEF scan, with window placement overlapping the diagonal scan line for doubly and triply charged peptides in the m/z – ion mobility plane across 16 × 25 m/z precursor isolation windows (resulting in 32 windows) defined from m/z 400 to 1,200, with 1 Da overlap, and CID collision energy ramped stepwise from 20 eV at 0.8 V·s/cm2 to 59eV at 1.3 V·s/cm2.

Endogenous INTS1 IP -/+ dTAG^V^-1 was acquired on Eclipse using DIA mode. Reconstituted peptides were separated using reverse-phase liquid chromatography on a 15 cm C18 fused silica column with an integrated emitter tip (IonOpticks, ID 75 µm, OD 360 µm, 1.6 µm C18 beads) using a nano-flow HPLC system (Thermo Ultimate 3000 RSLC Nano-LC) that is coupled to a Orbitrap Eclipse Tribrid Mass Spectrometer (Thermo Scientific) using Easy nLC source and electro sprayed directly into the mass spectrometer. Peptides were loaded directly onto the column at a flow rate of 600 nL/min with buffer A (99.9% Milli-Q water, 0.1% FA) and eluted at 400nL/min on a 30 min linear analytical gradient of increasing buffer B (90% ACN, 0.1% FA) from 2 to 34%. Data was acquired in a data independent (DIA) mode. MS1 spectra were acquired in the Orbitrap (R = 120k; normalised AGC target = 300%; MaxIT = custom; RF Lens = 40%; scan range = 375–1500; profile data). Dynamic exclusion was employed for 30 s excluding all charge states for a given precursor. MS2 spectra were collected in the Orbitrap (R = 30k; first mass = 120 m/z; normalised AGC target = 100%; MaxIT = 54 ms).

For NELFE and SPT5 IP experiments, the peptides were separated using reverse-phase liquid chromatography on a 15 cm C18 fused silica column with an integrated emitter tip (IonOpticks, ID 75 µm, OD 360 µm, 1.6 µm C18 beads) on a Thermo Scientific NeoVanquish LC coupled to Thermo Scientific Orbitrap Astral. Peptides were analysed on a 20 min linear analytical gradient of increasing buffer B (80% ACN, 0.1% FA) from 4 to 34%. Data was acquired in a data independent (DIA) mode. The MS1 settings were as follows: Orbitrap resolution 240,000; scan range m/z 380-980; AGC target 500%. The DIA parameters were as follows: isolation window: custom; HCD collision energy: 25%; scan range: m/z 145-1450; maximum injection time: 3 ms; AGC target: 800%.

Raw DIA data for the INTS12 endogenous IP and INTS12-SLF pulldown were analysed on DIA-NN (54). All other data was analysed on Spectronaut (55) version 19.9 using BGS factory settings of directDIA analysis with cysteine carbamidomethylation as fixed modification and N-terminal acetylation and methionine oxidations as variable modifications. Result filters m/z was set to 1800 (max) and 300 (min) with a relative intensity set to 5. DIA analysis identification had all precursor Qvalue cutoff, precursor PEP cutoff, protein Qvalue cutoff (experiment and run), and protein PEP cutoff set to 0.01. Human (Homo sapiens) reference proteome was used for database searching.

### Proteomics Analysis

Raw MS data were searched by DIA-NN (Version 1.8.1) in library-free mode with match-between-run (MBR) enabled. Precursor-level data from the second-pass search were used in the analysis for endogenous INTS12 IP and INTS12 SLF-AP experiments. Precursor intensities with ‘Global.Q.Value’ or ‘Lib.Q.Value’ greater than 0.01 were filtered out. Non-proteotypic precursors, single precursor protein groups, and compound protein groups were filtered out in preprocessing. Precursor-level intensities were log2-transformed. For the INTS1 immunoprecipitation, INTS12 TurboID, and RNAPII TurboID experiments, raw MS data were searched by Spectronaut (Version 19.9). ‘EG.TotalQuantity (Settings)’ was used for elution group intensities.

Log2 transformation was applied on precursor and elution group intensities in pre-processing. Differential expression (DE) analysis was performed using the limpa R package(49). Briefly, precursors (or elution groups) were summarized into proteins using the dpcQuant() function, which eliminates missing data from the protein-level data by representing the missing values using the detection probability curve (DPC)(49, 56). Unless otherwise specified, cyclic loess normalisation was applied on the protein level(57) using the normalizeCyclicLoess() function in limma(50). DE analysis was performed using the dpcDE() pipeline as implemented in the limpa^1^ package. A false discovery rate (FDR) threshold of 5% was used to define differential expression.

All data analysis was performed using LIMPA data analysis (49) and subsequently visualized in R.

### Chromatin Immunoprecipitation (ChIP) Sequencing

INTS12-FKBP cells were incubated with DMSO or dTAG^V^-1 (250nM) for 2 hours. ChIP-seq was performed as previously described (Vervoort et al. 2021 Cell). Cells were washed once with cold PBS prior to cross-linking at room temperature for 10 minutes with 1/10^th^ volume fresh formaldehyde solution (11% formaldehyde, 0.5mM EGTA, 1mM EDTA, 100mM NaCl, 50mM HEPES-KOH pH7.5) in RT PBS. To quench the cross-linking reaction, 1/20^th^ volume of 2.5M glycine was added and cells were incubated at room temperature for a further 5 minutes. Cells were washed with cold PBS, scraped and spun down. From scraping to the end of the washes, buffers were supplemented with cOmplete^TM^, EDTA-free Protease Inhibitor Cocktail and PhosSTOP^TM^ (Roche). Nuclei were isolated by three successive 10 minute incubations on ice with cold nuclear extraction buffer (20mM Tris-HCl pH 8, 10mM NaCl, 0.5% IGEPAL CA-630, 2mM EDTA). Cell nuclei were resuspended in sonication buffer (20mM Tris-HCl pH 7.5, 150mM NaCl, 2mM EDTA, 0.3% SDS, 1% IGEPAL CA-630) and sonicated for 8 minutes using the Covaris ME220 (Peak power 75; Duty Factor 15%; Cycles/Burst 1,000; Avg. Power 11.3). Fragmented chromatin was spun at top speed on a benchtop centrifuge for 20 minutes at 4 degrees. The supernatant was diluted 1:1 with dilution buffer (20mM Tris-HCl pH 8, 150mM NaCl, 2mM EDTA, 1% Triton X-100). Samples were quantified with the Qubit dsDNA BR assay (Invitrogen) and S2 (Drosophila) or NIH-3T3 (Mouse) chromatin was spiked into each sample. For each immunoprecipitation, diluted chromatin was rotated overnight at 4C with Pierce Protein A/G Magnetic beads (Thermo Scientific) and the indicated antibodies (Key Resource Table). The following day, beads were washed once each with ChIP buffer (20mM Tris-HCl pH 8, 150mM NaCl, 2mM EDTA, 0.15% SDS, 1% Triton X-100), ChIP wash buffer 1 (20mM Tris-HCl pH 8, 500mM NaCl, 2mM EDTA, 0.1% SDS, 1% Triton X-100) and ChIP wash buffer 2 (20mM Tris-HCl pH 8, 250mM LiCl, 2mM EDTA, 0.5% deoxycholate, 0.5% IGEPAL CA-630) and washed twice with TE buffer (10mM Tris-HCl pH 7.5, 1mM EDTA). Washed beads were incubated with shaking at 55C for 1 hour in reverse crosslinking buffer (200mM NaCl, 100mM NaHCO3, 1% SDS, 300µg/mL Proteinase K) followed by incubation of the supernatant overnight at 65C. DNA was isolated using the Zymogen ChIP DNA Clean and Concentrator Kit (Zymo Research D5205) following the manufacturer’s instructions. Sequencing libraries were prepared using the NEBNext Ultra II DNA Library Prep Kit (NEB E7645).

### ChIP-seq Data Analysis

Paired-end ChIP-seq reads were aligned to the human reference genome (GRCh38) using bowtie2 (v2.5.3). Alignments were performed in local mode with sensitive settings (--local –sensitive-local), not allowing mixed and discordant alignments (--no-mixed –no-discordant). Fragment sizes were constrained between 10 and 700 bp. Files were sorted and indexed using samtools and PCR duplicates were removed with Picard. Deduplicated files were then indexed, and quality was assessed using samtools (flagstat). Reads were aligned to mm10 or dm6 (GCA_000001215.4) in the same manner for spike-in control. For RNA pol II marks (3E10 and 4H8) CPM normalised bigwigs were generated using bamCoverage function from deeptools. Coverage was calculated using a bin size of 10bp. All other marks (INTS12, INTS1, NELFE, and NELFC) were normalised to an exogenous reference genome. Normalisation factors were calculated as α = 1/Nd (where Nd = drosophila or mm10 reads in millions) and applied in –scaleFactor in the bamCoverage function to generate bigwig files (Orlando et al., 2014).

To visualise enrichment around TSS, normalised bigwigs were analysed using computeMatrix in reference-point mode. Signal was centred at TSS (referencePoint center) and calculated in 10 bp bins with upstream and downstream windows of 2000 bp. For snRNA a smaller window of 1000 bp upstream and downstream was used. Missing data were treated as zero. The reference bed files that were used were all expressed genes, all protein coding genes, ints12 enriched genes and snRNAs.

### Transient Transcriptome Sequencing

For both standard and DRB TT-seq experiments, 5.5 x 10^6^ eHAP1s 12KI cells were seeded one day prior to treatment in 10 cm dishes containing 10 ml of complete growth medium.

For standard TT-seq, INTS12 degradation was induced by treatment with 250 nM dTAG^V^-1 for 1.5h Cells were additionally treated with 300 nM (batch 1 for replicate 1 and 2) or 150 nM (batch 1 for replicate 3 and 4) of AZ5576 during the final 30 mins of incubation (total 1.5h). Nascent RNA was labelled by the addition of 1 mM 4-thiouridine (4sU) for the final 15 mins.

For DRB TT-seq experiments, cells were treated with 250 nM dTAG^V^-1 for 1h, followed by the addition of 100 uM DRB for a total treatment time of 3.5h. Cells were then washed 2x with PBS and transcriptional release was performed for 15, 30 mins or 45 mins. Nascent RNA was labeled with 1 mM 4sU during the final 15 mins of each release.

Total RNA was isolated using TRIzol reagent according to the manufacturer’s instructions. RNA concentration was measured using Qubit RNA BR and NanoDrop. Samples were sonicated on a Covaris ME220 machine with the following settings: Duration, 30 seconds; Pear power, 75; Duty %, 25; Cycles/burst, 10,000; Average, 18.8; Temperature, 4-10 ∼6.4C. Fragmentation efficiency was assed using Agilent TapeStation for RNA.

For biotinylation of 4sU-labeled RNA, approximately 100 ug of sonicated RNA were incubated with MTSEA biotin-XX linker in biotinylation buffer (TRIS-HCL pH 7.4, EDTA) at 25C for 30 mins while shaking in the dark. Excess biotin reagent was removed by two sequential phenol:chloroform:isoamyl alcohol (pH 8) extractions, followed by isopropanol precipitation. RNA pellets were washed with 85% ethanol and resuspended in RNase-free water.

Incorporation of 4sU was validated by dot-blot analysis following the publicly available TT-chem-seq protocol (58). Briefly, diluted biotinylated RNA was immobilised on a Hybond-N membrane, UV crosslinked, and probed with HRP-conjugated streptavidin.

Signal was detected using enhanced chemiluminescence and total RNA loading was assessed by methylene blue stain. Biotinylated RNA was separated from unlabelled RNA using uMACS streptavidin Microbeads (Miltenyi; cat 130-133-282). RNA samples were denatured at 65C, rapidly cooled on ice and incubated with the beads at room temperature for 30 minutes. The mix of beads and RNA was then applied to the uMACS columns previously placed on the magnet. Columns were then washed with high-salt wash buffer (100 mM Tris-HCl, pH 7.4, 10 mM EDTA, 1 M NaCl and 0.1% (vol/vol) Tween 20.) at 65C and labeled RNA was eluted using freshly prepared dithiothreitol (DTT). Eluted RNA was purified using RNeasy MinElute spin columns (Qiagen, cat 74204), with modified binding conditions to retain short RNA fragments and further eluted in RNase-free water. RNA concentration was quantified using Qubit RNA HS kit.

Strand specific TT-seq libraries were prepared using the NEBNext Ultra II Directional RNA library prep kit for Illumina, following rRNA-depleted FFPE RNA protocol. Libraries were sequenced on an illumina NovaSeq X plus platform to generate 150 bp paired-end reads.

### TT-seq Analysis

STAR genome indices were generated using GENCODE v41 annotations. Reads were mapped to the human genome (GRCh38) using STAR (v2.7.11b) in paired end mode with two pass mapping enabled (--twopassMode Basic). Transcriptome alignments and gene level read counts were generated (–quantMode TranscriptomeSAM GeneCounts). Unmapped reads were excluded and PCR duplicates were identified and marked with Picard. Aligned reads were sorted and indexed using samtools.

Strand specific RPKM normalised bigwig files were generated using deepTools (bamCoverage --effectiveGenomeSize 2913022398 –binSize –extendReads – ignoreDuplicates –filterRNAstrand). Protein-coding genes expressed in the INTS12 dataset were separated by strand and used as input bed files for generating signal matrices (computeMatrix scale-regions) with 500 bp upstream and downstream, and a scaled gene body of 5kb. Signal was summarised in 10 bp bins, skipping regions with zero coverage. Forward and reverse strands were processed separately. For the resulting metagene profiles, mean signal per gene was computed across bins within each sample followed by averaging across all samples. Genes with extremely low (log2 mean < -5) or extremely high (>5) average signal were removed. The resulting matrix was filtered to retain protein coding genes between 100kb and 1mb for analysing processivity. To avoid bias from genes directly responsive to INTS12 degradation, INTS12 enriched genes were removed from DMSO/ dTAG^V^-1 experiments. For each sample the metagene profile was generated by dividing the raw smoothed signal at each bin by the sample specific median (within sample median normalisation) to emphasize relative signal distribution along gene bodies.

The processivity index was calculated from the median normalised metagene profiles as the ratio of signal in the first 1.25 kb (125 bins from TSS) and the last 1.25kb (125 bins from TES) of the gene body (5kb). Statistical significance between DMSO and dTag conditions was evaluated using a paired t-test with bin position used as the pairing factor.

The same workflow was implemented for CDK9i/dual experiments. INTS12 enriched genes were not removed.

### DRB TT-seq Analysis

Paired-end reads were processed in a similar manner to TT-seq. Duplicates were marked with picard, without removal. Strand-specific metagene matrices were computed using deeptools computeMatrix in reference-point mode, centred on TSS. Signal was extended 5kb upstream of the TSS and 150kb downstream, using 100 bp bins. Zeros were skipped and missing data were treated as zero.

Log2(dTAG/DMSO) genome wide signal tracks were generated using deeptools bamCompare. For each timepoint dTAG^V^-1 samples were compared to DMSO. Coverage was computed in 100bp bins and normalised to CPM for bam file corresponding to each condition. Log2 ratios were then calculated using a pseudocount value of 0.01 to avoid undefined values in low coverage regions. The resulting bigwigs represented genome wide log2(dtag/dmso) signal that was visualised using plotHeatmap from deeptools.

For the TSS centred profile plots, only protein coding genes with a total length between 100kb and 300kb were retained. Further filtering was applied to retain genes with a log2 mean signal between 1 and 6, based on overall signal distribution, and low distal end signal (last 50kb) at 15 minutes (<=1). Thes genes were excluded from all timepoints as they arise either from background or residual signal. For each sample, the mean signal across all bins was calculated and each bin was divided by the sample specific mean to emphasize positional differences along the gene body.

### Evolution Analysis

To investigate which INTS12 homologs could have similar functions to human INTS12, we aligned this gene family using PRANK and investigated the sequence conservation of each gene, comparing it to the human INTS12 protein.

### Quantification and Statistical Analysis

The number of replicates for each experiment (number of independent experiments is indicated in the figure legends. To determine statistical significance 2-way ANOVA.

## Acknowledgments

S.J.V. was supported by a CSL Centenary Fellowship and a Snow Medical Fellowship; L.D.C. was supported by an Australian Government Research Training Scholarship. This research was supported by the Snow Medical Research Foundation [SMRF2021-SF346]. This work was supported by the NHMRC Investigator grants to W.T. (GNT 2026635) and S.S. (GNT 2016827). The funders were not involved in the design of the study, collection, analysis, and interpretation of the data, the writing of this report, or the decision to submit the article for publication. We greatly thank the Walter and Eliza Hall Institute (WEHI) Genomics, Proteomic and FACS Facilities for data collection of samples (Vineet Vaibhav and Laura Dagley for proteomics facility). We thank the members of the Vervoort lab for critical discussions, and in particular Estelle Cui, Maede Salehi and Olivia Voulgaris for helping with experiments execution. The Brown laboratory at the Peter MacCallum Cancer Center (PMCC) for ATF3 antibody.

## Author contributions

Conceptualization: S.J.V; methodology: L.D.C. and S.J.V.; software: S.J.V., G.K.S., A.A.H., I.S.R., O.O. and C.A.N.G.; validation: L.D.C., I.S.R., A.A.H., C.A.N.G. and S.J.V.; formal analysis: S.J.V., L.D.C., I.S.R., A.A.H., C.A.N.G., M.L., L.N.; investigation: L.D.C., I.S.R., A.A.H., C.A.N.G., K.T., S.W., W.T., S.S., J.K.C.; resources: S.J.V.; data curation: SJ.V., L.D.C., I.S.R., A.A.H., C.A.N.G.; writing-original draft: S.J.V. (lead), L.D.C. (equal), A.A.H. (equal); writing-review and editing: SJ.V., L.D.C., I.S.R., A.A.H., C.A.N.G.; visualisation: S.J.V., L.D.C., I.S.R., A.A.H., C.A.N.G., O.O, L.N.; supervision: S.J.V., G.K.S., AMcL; project administration: S.J.V.; funding acquisition: S.J.V.

## Declaration of Interests

There are no conflicts of interests to declare.

## Notes

### Competing Interest Statement

The authors have declared no competing interest.

