## Additional_Information for "INTS12 Bridges Integrator and NELF to Prevent the Release of Non-processive RNA Polymerase II Complexes"

**INTS12 Left Homology Arm:**

TTTACTTTATTTTTTTAGACATCCACAGTTATTTCAGGAAATTCTTCTAGTGCCAGCGTTTCCTCGTCAGTAACTAGTGGCTTAACTGGATGGGCAGCTTTTGCAGCCAAAACTTCCTCTGCTGGTCCTTCAACAGCAAAATTGAGTTCAACAACACAAAACAATACTGGGAAACCTGCTACTTCGTCAGCTAACCAGAAACCTGTGGGTTTGACTGGTCTGGCAACATCATCCAAAGGTGGAATAGGTTCCAAAATAGGTTCCAATAACAGCACTACGCCCACTGTACCTTTAAAACCACCTCCACCTCTAACCTTGGGTAAAACTGGCCTTAGTCGCTCAGTTAGTTGTGACAATGTCAGCAAAGTAGGTCTTCCTAGTCCAAGTAGTTTAGTTCCAGGAAGCAGCAGCCAACTAAGTGGGAATGGAAATAGTGGAACATCAGGACCTAGTGGAAGTACTACCAGCAAAACTACTTCAGAATCCAGCAGCTCTCCCTCAGCATCCCTTAAAGGACCAACTTCACAAGAATCACAGCTCAATGCTATGAAGCGATTACAGATGGTCAAGAAGAAAGCTGCCCAAAAGAAACTCAAGAAG

**INTS12 Right Homology Arm:**

TAATGTGGCCAAGTAGGTTTTTGTATCATATTAGCCTAAAGATGAAAGGCTTATTATTATGATATAATCTGTAATACACTGTAATTTAATAAAAGTCTTCATAATCAAATTTCTATTGTTTCTTACCTTAAAATGCTAAATATTATATAACAAAACTTGGAAACTTTCATTTTAAATCAATCTTCAACATTTTTATTCTAATGATTTCCATAGTGTGTGAAAACAATGATTAGAATGACTGGTTCTCAATGAGATCTTCAGAAGATTGAAAGCATCTGTGAATTTAAGGAAAAAGAAATAGTTCATCTTCTAGCAAAGCCCCAAGAGTTGATCCTGATGAATGAGGTGCCTTATGTGGCGGCAGGGACGACCCAACATAGTTTTCTATTTTCCTCCCCGTATTCTGGGTGTTCATTTTCTGTGCTAGCCCCATTTCAACTGACTCTCCCTCTCCCTCCCAATTATTTCATCTTTTTCAAAATTATTTTTTCATTATTGATTTCTCATATCAATGGATTTTCCTAATTGATTTATCATACTTCATATTATATCTTTACCTTTTTTATCACCTCATTTCAATTATTTAAGGGTATATCTTTTTCC

**GS-linker-FKBP12^F36V^-GS-linker-HA-T2A-PuroR-T2A-EGFP insert:**

ATGGGAGTGCAGGTGGAAACCATCTCCCCAGGAGACGGGCGCACCTTCCCCAAGCGCGGCCAGACCTGCGTGGTGCACTACACCGGGATGCTTGAAGATGGAAAGAAAGTTGATTCCTCCCGGGACAGAAACAAGCCCTTTAAGTTTATGCTAGGCAAGCAGGAGGTGATCCGAGGCTGGGAAGAAGGGGTTGCCCAGATGAGTGTGGGTCAGAGAGCCAAACTGACTATATCTCCAGATTATGCCTATGGTGCCACTGGGCACCCAGGCATCATCCCACCACATGCCACTCTCGTCTTCGATGTGGAGCTTCTAAAACTGGAAGGAGGTTCAGGAGGTGGATCATACCCCTACGACGTACCTGATTACGCTGAGGGCAGAGGAAGTCTGCTAACATGCGGTGACGTCGAGGAGAATCCTGGCCCATCCGGAATGACCGAGTACAAGCCCACGGTGCGACTTGCAACAAGAGATGATGTACCTAGAGCTGTTAGAACTTTGGCAGCTGCATTTGCTGATTATCCTGCAACTAGACATACAGTAGATCCAGATCGACACATCGAGCGGGTCACCGAGCTGCAAGAACTCTTCCTCACGCGCGTCGGGCTCGACATCGGAAAAGTTTGGGTTGCAGATGATGGTGCTGCAGTTGCTGTTTGGACAACTCCAGAATCTGTTGAAGCTGGAGCAGTGTTCGCTGAAATCGGACCTCGCATGGCCGAGTTGAGCGGATCAAGACTGGCTGCTCAACAACAGATGGAAGGCCTCCTGGCGCCGCACCGGCCCAAGGAGCCCGCGTGGTTCCTGGCTACTGTTGGAGTTTCTCCTGATCATCAAGGAAAAGGTCTTGGTTCTGCTGTAGTACTTCCAGGTGTTGAAGCAGCAGAACGAGCTGGAGTACCAGCTTTCCTGGAGACCTCCGCGCCCCGCAACCTCCCCTTCTACGAGCGGCTCGGCTTCACCGTCACCGCCGACGTCGAGGTGCCCGAAGGACCGCGCACCTGGTGCATGACCCGCAAGCCCGGTGCCTCCGGAGAGGGCAGAGGAAGTCTGCTAACATGCGGTGACGTCGAGGAGAATCCTGGCCCAATGGTGAGCAAGGGCGAGGAGCTGTTCACCGGGGTGGTGCCCATCCTGGTCGAGCTGGACGGCGACGTAAACGGCCACAAGTTCAGCGTGTCTGGCGAGGGCGAGGGCGATGCCACCTACGGCAAGCTGACCCTGAAGTTCATCTGCACCACCGGCAAGCTGCCCGTGCCCTGGCCCACCCTCGTGACCACCCTGACCTACGGCGTGCAGTGCTTCAGCCGCTACCCCGACCACATGAAGCAGCACGACTTCTTCAAGTCCGCCATGCCCGAAGGCTACGTCCAGGAGCGCACCATCTTCTTCAAGGACGACGGCAACTACAAGACCCGCGCCGAGGTGAAGTTCGAGGGCGACACCCTGGTGAACCGCATCGAGCTGAAGGGCATCGACTTCAAGGAGGACGGCAACATCCTGGGGCACAAGCTGGAGTACAACTACAACAGCCACAACGTCTATATCATGGCCGACAAGCAGAAGAACGGCATCAAGGCGAACTTCAAGATCCGCCACAACATCGAGGACGGCAGCGTGCAGCTCGCCGACCACTACCAGCAGAACACCCCCATCGGCGACGGCCCCGTGCTGCTGCCCGACAACCACTACCTGAGCACCCAGTCCGCCCTGAGCAAAGACCCCAACGAGAAGCGCGATCACATGGTCCTGCTGGAGTTCGTGACCGCCGCCGGGATCACTCTCGGCATGGACGAGCTGTAC

GS-linker-TurboID-GS-linker-HA-T2A-PuroR-T2A-EGFP insert:

GGGAGGTTCTGGTGGAGGTTCTaaggacaataccgtgcctctcaagctgatcgccctgctggccaacggcgagttccacagcggcgagcagctgggagagacactgggcatgtctagagccgccatcaacaagcacatccagaccctgagagattggggcgtggacgtgttcaccgtgcccggcaagggctacagcctgccagaacccatccccctgctgaacgccaagcaaatcctgggccagctggatggcggatctgtggccgtgctgccagtggtcgacagcaccaaccagtacctgttggatagaatcggcgagctgaaaagcggagatgcttgcatcgccgagtaccagcaggccggcagaggtagccggggcagaaagtggttcagcccttttggcgctaatctgtatttgagcatgttctggcggctgaaaagaggacctgccgctatcgggctgggccctgtgatcggcatcgtgatggccgaggccctgcggaaactgggcgccgacaaggtgcgggtgaagtggcctaacgacctgtacctgcaagacagaaagctggctggcatcctggtggaactggccggcatcacaggcgacgccgcccagattgtgatcggcgctggcatcaacgtggccatgcggagagtggaagagagcgtggtgaaccagggatggatcacactgcaggaggccggaattaacctggaccggaacaccctggctgccaccctgatccgcgaactgagagccgctctggaactgtttgagcaggagggactggccccttacctgcctagatgggagaagctcgacaacttcatcaatagacctgtcaagctgatcatcggagataaggaaatcttcggcatttctagaggcatcgacaaacagggcgccctgctgctggaacaggacggcgttatcaagccctggatgggcggcgagatctccctgaggtccgccgaaaagGGAGGTTCAGGAGGTGGATCATACCCCTACGACGTACCTGATTACGCTGAGGGCAGAGGAAGTCTGCTAACATGCGGTGACGTCGAGGAGAATCCTGGCCCATCCGGAATGACCGAGTACAAGCCCACGGTGCGACTTGCAACAAGAGATGATGTACCTAGAGCTGTTAGAACTTTGGCAGCTGCATTTGCTGATTATCCTGCAACTAGACATACAGTAGATCCAGATCGACACATCGAGCGGGTCACCGAGCTGCAAGAACTCTTCCTCACGCGCGTCGGGCTCGACATCGGAAAAGTTTGGGTTGCAGATGATGGTGCTGCAGTTGCTGTTTGGACAACTCCAGAATCTGTTGAAGCTGGAGCAGTGTTCGCTGAAATCGGACCTCGCATGGCCGAGTTGAGCGGATCAAGACTGGCTGCTCAACAACAGATGGAAGGCCTCCTGGCGCCGCACCGGCCCAAGGAGCCCGCGTGGTTCCTGGCTACTGTTGGAGTTTCTCCTGATCATCAAGGAAAAGGTCTTGGTTCTGCTGTAGTACTTCCAGGTGTTGAAGCAGCAGAACGAGCTGGAGTACCAGCTTTCCTGGAGACCTCCGCGCCCCGCAACCTCCCCTTCTACGAGCGGCTCGGCTTCACCGTCACCGCCGACGTCGAGGTGCCCGAAGGACCGCGCACCTGGTGCATGACCCGCAAGCCCGGTGCCTCCGGAGAGGGCAGAGGAAGTCTGCTAACATGCGGTGACGTCGAGGAGAATCCTGGCCCAATGGTGAGCAAGGGCGAGGAGCTGTTCACCGGGGTGGTGCCCATCCTGGTCGAGCTGGACGGCGACGTAAACGGCCACAAGTTCAGCGTGTCTGGCGAGGGCGAGGGCGATGCCACCTACGGCAAGCTGACCCTGAAGTTCATCTGCACCACCGGCAAGCTGCCCGTGCCCTGGCCCACCCTCGTGACCACCCTGACCTACGGCGTGCAGTGCTTCAGCCGCTACCCCGACCACATGAAGCAGCACGACTTCTTCAAGTCCGCCATGCCCGAAGGCTACGTCCAGGAGCGCACCATCTTCTTCAAGGACGACGGCAACTACAAGACCCGCGCCGAGGTGAAGTTCGAGGGCGACACCCTGGTGAACCGCATCGAGCTGAAGGGCATCGACTTCAAGGAGGACGGCAACATCCTGGGGCACAAGCTGGAGTACAACTACAACAGCCACAACGTCTATATCATGGCCGACAAGCAGAAGAACGGCATCAAGGCGAACTTCAAGATCCGCCACAACATCGAGGACGGCAGCGTGCAGCTCGCCGACCACTACCAGCAGAACACCCCCATCGGCGACGGCCCCGTGCTGCTGCCCGACAACCACTACCTGAGCACCCAGTCCGCCCTGAGCAAAGACCCCAACGAGAAGCGCGATCACATGGTCCTGCTGGAGTTCGTGACCGCCGCCGGGATCACTCTCGGCATGGACGAGCTGTAC

Full Length (FL) INTS12- 3X T7-tag- P2A -Hygro:

ATGGCTGCTACTGTGAACTTGGAACTTGATCCCATTTTTTTGAAAGCACTAGGTTTCTTGCATTCAAAGAGTAAAGATTCTGCTGAAAAGCTAAAAGCACTGCTTGATGAATCTTTGGCTCGGGGCATTGATTCCAGTTACCGTCCATCTCAAAAGGATGTGGAGCCACCCAAAATTTCAAGCACAAAAAACATTTCCATTAAGCAAGAGCCCAAAATATCATCCAGTCTTCCTTCTGGTAATAATAATGGCAAGGTCCTCACAACTGAAAAGGTAAAGAAGGAAGCTGAAAAGAGACCTGCTGATAAAATGAAATCAGACATCACTGAAGGAGTTGATATTCCAAAGAAACCTAGATTGGAGAAACCAGAAACACAGTCATCTCCCATTACTGTCCAAAGTAGCAAGGATTTACCTATGGCTGACCTTTCCAGTTTTGAGGAGACCAGTGCTGATGATTTTGCCATGGAGATGGGATTGGCCTGCGTTGTTTGTAGGCAAATGATGGTGGCATCTGGCAATCAATTAGTAGAATGTCAGGAGTGCCATAATCTCTACCACCGAGATTGTCATAAACCCCAGGTGACAGACAAGGAAGCGAATGACCCTCGCCTGGTGTGGTATTGTGCCCGATGTACCAGACAAATGAAAAGAATGGCTCAAAAAACTCAGAAACCACCGCAGAAACCAGCCCCTGCAGTTGTTTCTGTAACTCCAGCTGTCAAAGATCCATTGGTTAAGAAACCAGAAACTAAACTGAAACAAGAGACAACTTTTCTAGCGTTTAAGAGAACAGAAGTCAAGACATCCACAGTTATTTCAGGAAATTCTTCTAGTGCCAGCGTTTCCTCGTCAGTAACTAGTGGCTTAACTGGATGGGCAGCTTTTGCAGCCAAAACTTCCTCTGCTGGTCCTTCAACAGCAAAATTGAGTTCAACAACACAAAACAATACTGGGAAACCTGCTACTTCGTCAGCTAACCAGAAACCTGTGGGTTTGACTGGTCTGGCAACATCATCCAAAGGTGGAATAGGTTCCAAAATAGGTTCCAATAACAGCACTACGCCCACTGTACCTTTAAAACCACCTCCACCTCTAACCTTGGGTAAAACTGGCCTTAGTCGCTCAGTTAGTTGTGACAATGTCAGCAAAGTAGGTCTTCCTAGTCCAAGTAGTTTAGTTCCAGGAAGCAGCAGCCAACTAAGTGGGAATGGAAATAGTGGAACATCAGGACCTAGTGGAAGTACTACCAGCAAAACTACTTCAGAATCCAGCAGCTCTCCCTCAGCATCCCTTAAAGGCCCAACTTCACAAGAATCACAGCTCAATGCTATGAAGCGATTACAGATGGTCAAGAAGAAAGCTGCCCAAAAGAAACTCAAGAAGATGGCCAGCATGACCGGCGGACAACAGATGGGAGGCTCTAGCGCCTCTATGGCTAGCATGACCGGCGGCCAGCAGATGGGCGGAAGCAGCGCCAGAATGGCCAGCATGACAGGCGGCCAGCAGATGGGCGGCTCCGGCgcaacaaacttctctctgctgaaacaagccggagatgtcgaagagaatcctggaccgatgaaaaagcctgaactcaccgcgacgtctgtcgagaagtttctgatcgaaaagttcgacagcgtctccgacctgatgcagctctcggagggcgaagaatctcgtgctttcagcttcgatgtaggagggcgtggatatgtcctgcgggtaaatagctgcgccgatggtttctacaaagatcgttatgtttatcggcactttgcatcggccgcgctcccgattccggaagtgcttgacattggggagttcagcgagagcctgacctattgcatctcccgccgtgcacagggtgtcacgttgcaagacctgcctgaaaccgaactgcccgctgttctgcagccggtcgcggaggccatggatgcgatcgctgcggccgatcttagccagacgagcgggttcggcccattcggaccgcaaggaatcggtcaatacactacatggcgtgatttcatatgcgcgattgctgatccccatgtgtatcactggcaaactgtgatggacgacaccgtcagtgcgtccgtcgcgcaggctctcgatgagctgatgctttgggccgaggactgccccgaagtccggcacctcgtgcacgcggatttcggctccaacaatgtcctgacggacaatggccgcataacagcggtcattgactggagcgaggcgatgttcggggattcccaatacgaggtcgccaacatcttcttctggaggccgtggttggcttgtatggagcagcagacgcgctacttcgagcggaggcatccggagcttgcaggatcgccgcggctccgggcgtatatgctccgcattggtcttgaccaactctatcagagcttggttgacggcaatttcgatgatgcagcttgggcgcagggtcgatgcgacgcaatcgtccgatccggagccgggactgtcgggcgtacacaaatcgcccgcagaagcgcggccgtctggaccgatggctgtgtagaagtactcgccgatagtggaaaccgacgccccagcactcgtccgagggcaaaggaataa

C-term deletion (ΔC) INTS12- 3X T7-tag- P2A -Hygro:

ATGGCTGCTACTGTGAACTTGGAACTTGATCCCATTTTTTTGAAAGCACTAGGTTTCTTGCATTCAAAGAGTAAAGATTCTGCTGAAAAGCTAAAAGCACTGCTTGATGAATCTTTGGCTCGGGGCATTGATTCCAGTTACCGTCCATCTCAAAAGGATGTGGAGCCACCCAAAATTTCAAGCACAAAAAACATTTCCATTAAGCAAGAGCCCAAAATATCATCCAGTCTTCCTTCTGGTAATAATAATGGCAAGGTCCTCACAACTGAAAAGGTAAAGAAGGAAGCTGAAAAGAGACCTGCTGATAAAATGAAATCAGACATCACTGAAGGAGTTGATATTCCAAAGAAACCTAGATTGGAGAAACCAGAAACACAGTCATCTCCCATTACTGTCCAAAGTAGCAAGGATTTACCTATGGCTGACCTTTCCAGTTTTGAGGAGACCAGTGCTGATGATTTTGCCATGGAGATGGGATTGGCCTGCGTTGTTTGTAGGCAAATGATGGTGGCATCTGGCAATCAATTAGTAGAATGTCAGGAGTGCCATAATCTCTACCACCGAGATTGTCATAAACCCCAGGTGACAGACAAGGAAGCGAATGACCCTCGCCTGGTGTGGTATTGTGCCCGATGTATGGCCAGCATGACCGGCGGACAACAGATGGGAGGCTCTAGCGCCTCTATGGCTAGCATGACCGGCGGCCAGCAGATGGGCGGAAGCAGCGCCAGAATGGCCAGCATGACAGGCGGCCAGCAGATGGGCGGCTCCGGCgcaacaaacttctctctgctgaaacaagccggagatgtcgaagagaatcctggaccgatgaaaaagcctgaactcaccgcgacgtctgtcgagaagtttctgatcgaaaagttcgacagcgtctccgacctgatgcagctctcggagggcgaagaatctcgtgctttcagcttcgatgtaggagggcgtggatatgtcctgcgggtaaatagctgcgccgatggtttctacaaagatcgttatgtttatcggcactttgcatcggccgcgctcccgattccggaagtgcttgacattggggagttcagcgagagcctgacctattgcatctcccgccgtgcacagggtgtcacgttgcaagacctgcctgaaaccgaactgcccgctgttctgcagccggtcgcggaggccatggatgcgatcgctgcggccgatcttagccagacgagcgggttcggcccattcggaccgcaaggaatcggtcaatacactacatggcgtgatttcatatgcgcgattgctgatccccatgtgtatcactggcaaactgtgatggacgacaccgtcagtgcgtccgtcgcgcaggctctcgatgagctgatgctttgggccgaggactgccccgaagtccggcacctcgtgcacgcggatttcggctccaacaatgtcctgacggacaatggccgcataacagcggtcattgactggagcgaggcgatgttcggggattcccaatacgaggtcgccaacatcttcttctggaggccgtggttggcttgtatggagcagcagacgcgctacttcgagcggaggcatccggagcttgcaggatcgccgcggctccgggcgtatatgctccgcattggtcttgaccaactctatcagagcttggttgacggcaatttcgatgatgcagcttgggcgcagggtcgatgcgacgcaatcgtccgatccggagccgggactgtcgggcgtacacaaatcgcccgcagaagcgcggccgtctggaccgatggctgtgtagaagtactcgccgatagtggaaaccgacgccccagcactcgtccgagggcaaaggaataa

N-term deletion (ΔN) INTS12- 3X T7-tag- P2A -Hygro:

ATGTTGGCCTGCGTTGTTTGTAGGCAAATGATGGTGGCATCTGGCAATCAATTAGTAGAATGTCAGGAGTGCCATAATCTCTACCACCGAGATTGTCATAAACCCCAGGTGACAGACAAGGAAGCGAATGACCCTCGCCTGGTGTGGTATTGTGCCCGATGTACCAGACAAATGAAAAGAATGGCTCAAAAAACTCAGAAACCACCGCAGAAACCAGCCCCTGCAGTTGTTTCTGTAACTCCAGCTGTCAAAGATCCATTGGTTAAGAAACCAGAAACTAAACTGAAACAAGAGACAACTTTTCTAGCGTTTAAGAGAACAGAAGTCAAGACATCCACAGTTATTTCAGGAAATTCTTCTAGTGCCAGCGTTTCCTCGTCAGTAACTAGTGGCTTAACTGGATGGGCAGCTTTTGCAGCCAAAACTTCCTCTGCTGGTCCTTCAACAGCAAAATTGAGTTCAACAACACAAAACAATACTGGGAAACCTGCTACTTCGTCAGCTAACCAGAAACCTGTGGGTTTGACTGGTCTGGCAACATCATCCAAAGGTGGAATAGGTTCCAAAATAGGTTCCAATAACAGCACTACGCCCACTGTACCTTTAAAACCACCTCCACCTCTAACCTTGGGTAAAACTGGCCTTAGTCGCTCAGTTAGTTGTGACAATGTCAGCAAAGTAGGTCTTCCTAGTCCAAGTAGTTTAGTTCCAGGAAGCAGCAGCCAACTAAGTGGGAATGGAAATAGTGGAACATCAGGACCTAGTGGAAGTACTACCAGCAAAACTACTTCAGAATCCAGCAGCTCTCCCTCAGCATCCCTTAAAGGCCCAACTTCACAAGAATCACAGCTCAATGCTATGAAGCGATTACAGATGGTCAAGAAGAAAGCTGCCCAAAAGAAACTCAAGAAGATGGCCAGCATGACCGGCGGACAACAGATGGGAGGCTCTAGCGCCTCTATGGCTAGCATGACCGGCGGCCAGCAGATGGGCGGAAGCAGCGCCAGAATGGCCAGCATGACAGGCGGCCAGCAGATGGGCGGCTCCGGCgcaacaaacttctctctgctgaaacaagccggagatgtcgaagagaatcctggaccgatgaaaaagcctgaactcaccgcgacgtctgtcgagaagtttctgatcgaaaagttcgacagcgtctccgacctgatgcagctctcggagggcgaagaatctcgtgctttcagcttcgatgtaggagggcgtggatatgtcctgcgggtaaatagctgcgccgatggtttctacaaagatcgttatgtttatcggcactttgcatcggccgcgctcccgattccggaagtgcttgacattggggagttcagcgagagcctgacctattgcatctcccgccgtgcacagggtgtcacgttgcaagacctgcctgaaaccgaactgcccgctgttctgcagccggtcgcggaggccatggatgcgatcgctgcggccgatcttagccagacgagcgggttcggcccattcggaccgcaaggaatcggtcaatacactacatggcgtgatttcatatgcgcgattgctgatccccatgtgtatcactggcaaactgtgatggacgacaccgtcagtgcgtccgtcgcgcaggctctcgatgagctgatgctttgggccgaggactgccccgaagtccggcacctcgtgcacgcggatttcggctccaacaatgtcctgacggacaatggccgcataacagcggtcattgactggagcgaggcgatgttcggggattcccaatacgaggtcgccaacatcttcttctggaggccgtggttggcttgtatggagcagcagacgcgctacttcgagcggaggcatccggagcttgcaggatcgccgcggctccgggcgtatatgctccgcattggtcttgaccaactctatcagagcttggttgacggcaatttcgatgatgcagcttgggcgcagggtcgatgcgacgcaatcgtccgatccggagccgggactgtcgggcgtacacaaatcgcccgcagaagcgcggccgtctggaccgatggctgtgtagaagtactcgccgatagtggaaaccgacgccccagcactcgtccgagggcaaaggaataa

PHD deletion (ΔPHD) INTS12- 3X T7-tag- P2A -Hygro:

ATGGCTGCTACTGTGAACTTGGAACTTGATCCCATTTTTTTGAAAGCACTAGGTTTCTTGCATTCAAAGAGTAAAGATTCTGCTGAAAAGCTAAAAGCACTGCTTGATGAATCTTTGGCTCGGGGCATTGATTCCAGTTACCGTCCATCTCAAAAGGATGTGGAGCCACCCAAAATTTCAAGCACAAAAAACATTTCCATTAAGCAAGAGCCCAAAATATCATCCAGTCTTCCTTCTGGTAATAATAATGGCAAGGTCCTCACAACTGAAAAGGTAAAGAAGGAAGCTGAAAAGAGACCTGCTGATAAAATGAAATCAGACATCACTGAAGGAGTTGATATTCCAAAGAAACCTAGATTGGAGAAACCAGAAACACAGTCATCTCCCATTACTGTCCAAAGTAGCAAGGATTTACCTATGGCTGACCTTTCCAGTTTTGAGGAGACCAGTGCTGATGATTTTGCCATGGAGATGGGAACCAGACAAATGAAAAGAATGGCTCAAAAAACTCAGAAACCACCGCAGAAACCAGCCCCTGCAGTTGTTTCTGTAACTCCAGCTGTCAAAGATCCATTGGTTAAGAAACCAGAAACTAAACTGAAACAAGAGACAACTTTTCTAGCGTTTAAGAGAACAGAAGTCAAGACATCCACAGTTATTTCAGGAAATTCTTCTAGTGCCAGCGTTTCCTCGTCAGTAACTAGTGGCTTAACTGGATGGGCAGCTTTTGCAGCCAAAACTTCCTCTGCTGGTCCTTCAACAGCAAAATTGAGTTCAACAACACAAAACAATACTGGGAAACCTGCTACTTCGTCAGCTAACCAGAAACCTGTGGGTTTGACTGGTCTGGCAACATCATCCAAAGGTGGAATAGGTTCCAAAATAGGTTCCAATAACAGCACTACGCCCACTGTACCTTTAAAACCACCTCCACCTCTAACCTTGGGTAAAACTGGCCTTAGTCGCTCAGTTAGTTGTGACAATGTCAGCAAAGTAGGTCTTCCTAGTCCAAGTAGTTTAGTTCCAGGAAGCAGCAGCCAACTAAGTGGGAATGGAAATAGTGGAACATCAGGACCTAGTGGAAGTACTACCAGCAAAACTACTTCAGAATCCAGCAGCTCTCCCTCAGCATCCCTTAAAGGCCCAACTTCACAAGAATCACAGCTCAATGCTATGAAGCGATTACAGATGGTCAAGAAGAAAGCTGCCCAAAAGAAACTCAAGAAG ATGGCCAGCATGACCGGCGGACAACAGATGGGAGGCTCTAGCGCCTCTATGGCTAGCATGACCGGCGGCCAGCAGATGGGCGGAAGCAGCGCCAGAATGGCCAGCATGACAGGCGGCCAGCAGATGGGCGGCTCCGGCgcaacaaacttctctctgctgaaacaagccggagatgtcgaagagaatcctggaccgatgaaaaagcctgaactcaccgcgacgtctgtcgagaagtttctgatcgaaaagttcgacagcgtctccgacctgatgcagctctcggagggcgaagaatctcgtgctttcagcttcgatgtaggagggcgtggatatgtcctgcgggtaaatagctgcgccgatggtttctacaaagatcgttatgtttatcggcactttgcatcggccgcgctcccgattccggaagtgcttgacattggggagttcagcgagagcctgacctattgcatctcccgccgtgcacagggtgtcacgttgcaagacctgcctgaaaccgaactgcccgctgttctgcagccggtcgcggaggccatggatgcgatcgctgcggccgatcttagccagacgagcgggttcggcccattcggaccgcaaggaatcggtcaatacactacatggcgtgatttcatatgcgcgattgctgatccccatgtgtatcactggcaaactgtgatggacgacaccgtcagtgcgtccgtcgcgcaggctctcgatgagctgatgctttgggccgaggactgccccgaagtccggcacctcgtgcacgcggatttcggctccaacaatgtcctgacggacaatggccgcataacagcggtcattgactggagcgaggcgatgttcggggattcccaatacgaggtcgccaacatcttcttctggaggccgtggttggcttgtatggagcagcagacgcgctacttcgagcggaggcatccggagcttgcaggatcgccgcggctccgggcgtatatgctccgcattggtcttgaccaactctatcagagcttggttgacggcaatttcgatgatgcagcttgggcgcagggtcgatgcgacgcaatcgtccgatccggagccgggactgtcgggcgtacacaaatcgcccgcagaagcgcggccgtctggaccgatggctgtgtagaagtactcgccgatagtggaaaccgacgccccagcactcgtccgagggcaaaggaataa
